# Pandemic Coronavirus Genome Packaging Relies on Multiple Dispersed Packaging Signals

**DOI:** 10.64898/2026.06.18.733228

**Authors:** Taha Y. Taha, Nikesh Patel, Sam Clark, Julia Rosecrans, Abdullah Syed, Rebecca Chandler-Bostock, Erik R. Farquhar, Lekshmi B.G. Nair, Abid Javed, Jennifer Doudna, Peter G. Stockley, Melanie Ott, Reidun Twarock

## Abstract

Packaging of viral genomes into progeny virions is a critical step in the viral life cycle. Coronaviruses such as SARS-CoV-2 possess unusually large RNA genomes (∼30 kb), yet the mechanism by which these genomes are selectively condensed and incorporated into virions remains poorly understood. Here, we demonstrate that the SARS-CoV-2 genome contains multiple dispersed RNA structures, termed packaging signals (PSs), which cooperate with a dominant central PS to direct genomic RNA incorporation into infectious particles. We identify the dominant PS within the coding region of the *nsp15* gene, downstream of a packaging signal previously described in Embecoviruses. Using virus-like particles (VLPs), we investigate its role in virion assembly and selective genomic RNA packaging, and show that it promotes the formation of ribonucleoprotein (RNP) complexes with the viral nucleocapsid protein (N), as revealed by mass photometry. Notably, we uncover a unique N–induced conformational rearrangement of the PS RNA, from an extended structure to a double stem-loop architecture. This dominant PS acts together with a nearby stem-loop element to assemble a higher-order RNP complex containing 12 N-protein dimers. Using a SARS-CoV-2 replicon system, we further demonstrate functional cooperativity between the dominant PS, its proximal partner stem-loop, and additional packaging elements located approximately 10 kb upstream. Collectively, our findings support a highly dynamic and cooperative mechanism of SARS-CoV-2 genome packaging that relies on multiple dispersed packaging signals organized around a dominant central PS. These insights provide a mechanistic framework for understanding coronavirus genome packaging and reveal new opportunities for antiviral intervention through disruption of this process. They also have important implications for vaccine development and may enable the design of membrane-based vector systems capable of efficiently delivering large nucleic acid cargoes, expanding the potential of bionanotechnology and genetic medicine.

---

Since Crick and Watson’s seminal paper reporting the discovery of highly symmetric virus architectures 70 years ago^1^, virus assembly has been considered the archetype of biological self-assembly. In their simplest form, viruses consist of a spherical protein shell surrounding the genomic nucleic acid, which is typically < 10 kb in length. In these viruses, that also comprise major viral pathogens such as Picornaviruses^2,3^ and Hepatitis B virus^4^, highly selective packaging of the genomic RNA is achieved against a backdrop of cellular RNAs via multiple dispersed packaging signals^5^, that act collectively in virion formation^6^. The mechanism has also evolved under directed evolution in non-viral protein containers^7^. However, until now it has not been fully characterised in larger and more complex viruses, such as the membrane-enclosed coronaviruses.

Coronaviruses have positive-sense, single-stranded genomic RNAs (gRNAs) around 30kb in length. Akin to eukaryotic mRNAs, they are endowed with a 5′ cap and a 3′ polyadenylated tail, and act as messenger RNAs during the infection. Due to their large genome sizes, the genome packaging and delivery strategies for viruses in this family differ from those of simple RNA viruses. Instead of a protein shell, they envelope their gRNAs into membranes studded by glycoproteins, which include the receptor-binding Spike (S) protein, as well as Membrane (M) and Envelope (E) proteins^8^. Nucleocapsid (N) proteins interact with the genome, forming Ribonucleoprotein (RNP) complexes to facilitate genome compaction. In the *Embecovirus* subgenus, one of the five subgenera of the *Betacoronaviruses*, a packaging signal (PS) is known to mediate the contact between an RNP complex containing six N protein dimers, and M protein, thus anchoring the RNP complex at the inner membrane to enable the selective incorporation of the gRNA into the virion^9,10^.

The *Embecovirus* PS is the most thoroughly characterised coronavirus PS, such as mouse hepatitis virus^11^ (MHV, Fig. 1a), with similar signals characterised also in bovine coronavirus (BCoV)^12^, and the endemic human coronaviruses (HCoVs) HCoV-OC43 and HCoV-HKU1. Severe acute respiratory syndrome coronavirus 2 (SARS-CoV-2), the causative agent of COVID-19, SARS-CoV, and Middle East respiratory syndrome coronavirus (MERS-CoV), are *Sarbecoviruses*, a subgenus of the *Betacoronavirus*es. In contrast to the extensively studied non-pandemic coronaviruses, *Sarbecoviruses* have a deletion in the coding sequence of the *nsp15* gene in *orf1b*^*1*3^ (Fig. S1a) where the MHV PS is located, thereby complicating the identification of the SARS-CoV-2 PS by analogy to the MHV PS on the basis of sequence alignment.

**Fig. 1.**
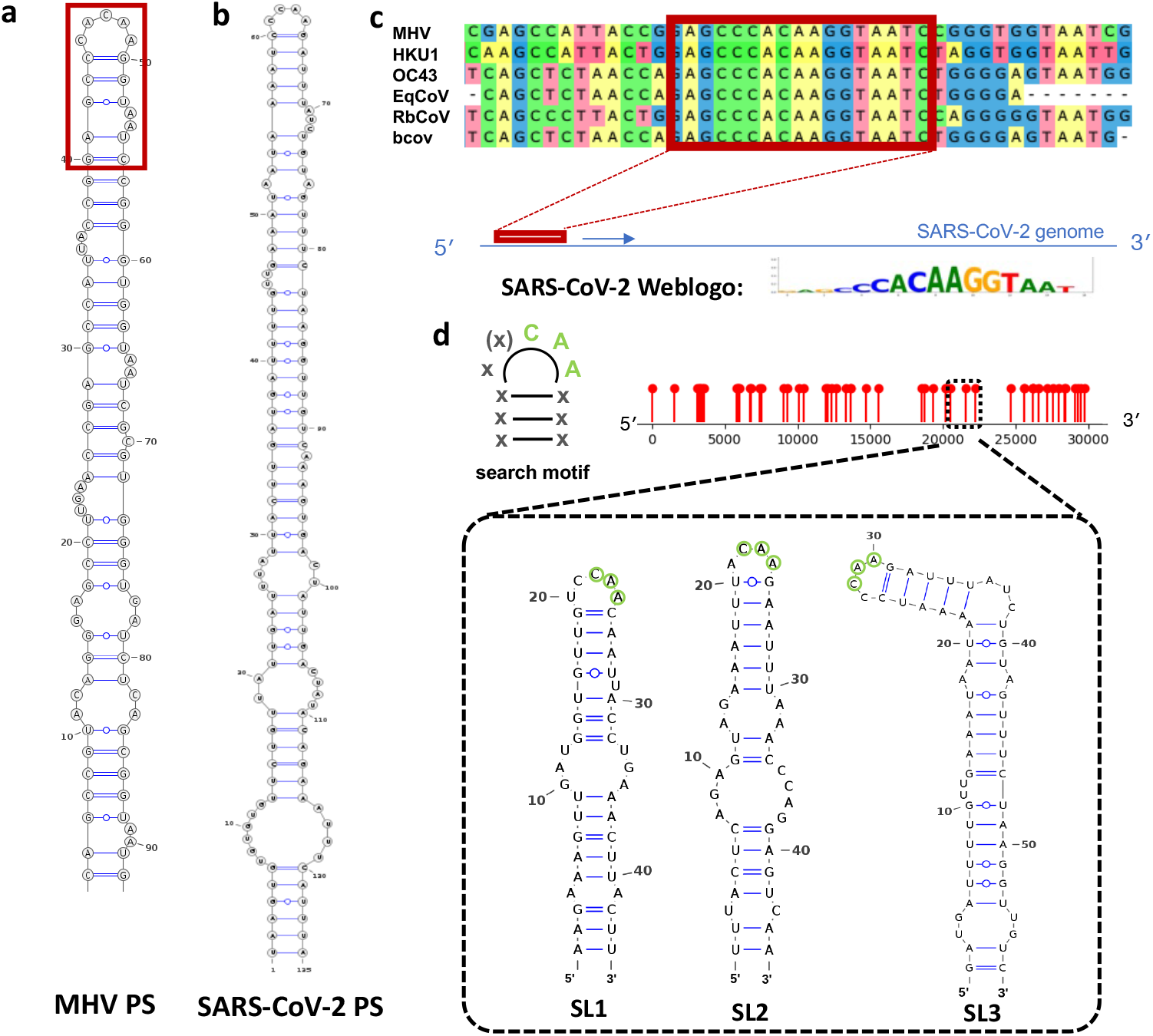
Identification of the SARS-CoV-2 PS. **a** The packaging signal (PS) in MHV, and **b** the predicted SARS-CoV-2 PS. **c** Alignment of the sequences corresponding to the top portions of the known PSs in different beta-coronaviruses identifies a 17 nt-long conserved sequence (red box). **c** A bioinformatics search for contiguous sequences of at least 5nt length in the SARS-CoV-2 genome matching this sequence identifies 249 hits; their alignment as a Weblogo^14^ reveals a highly conserved ‘CAA’ motif. **d** The implied search motif (left) and the locations of such sites in the SARS-CoV-2 genome shown as lollipops (right); the secondary structures of the three sites in the PS9 fragment are shown (dashed box).

Here we identify and characterise the *Sarbecovirus* PS using a bioinformatics search informed by the 6-fold degenerate structure of an RNP complex. As in smaller non-enveloped RNA viruses, where the 60-fold symmetric repeat of the PS binding site at the inner capsid surface was the key to unravelling the multiple PS packaging code in these viruses, the symmetry of an RNP complex is the key to identifying an equivalent mechanism in the more complex coronaviruses here. In this highly interdisciplinary effort, combining theory with mutagenesis in a VLP and replicon systems as well as mass photometry, we uncover a complex and highly dynamic process that may serve as a drug target or blueprint for packaging large genomes into viral vectors.

## Identification of the SARS-CoV-2 PS

As a means of determining a common motif in the known *Embecovirus* PSs, we aligned their sequences (Fig. 1c). This identified a contiguous 17nt-long, fully conserved fragment encompassing the loop portion of the MHV PS (red box). We slid this motif in increments of 1nt along the SARS-CoV-2 genome (NC_045512.2), recording any contiguous matches at least 5 nts in length. This resulted in 249 hits, that we used to build a Weblogo^14^, revealing the most frequent match to include a ‘CAA’ motif. As this motif overlaps with the loop of the known MHV PS (Fig. 1a), the SARS-CoV-2 PS should also contain a ‘CAA’ loop motif. We therefore identified all sites in the SARS-CoV-2 genome with an ‘xxCAA’ or ‘xCAA’ loop followed by three base-pairs (*cf*. search motif in Fig. 1d).

In other RNA viruses presenting a multiple packaging signal mediated assembly mechanism, there is typically a dominant PSs that is distinguished, e.g., by its folding free energy^6^. To identify such a PS site in SARS-CoV-2, we took additional criteria, such as stem-loop length and folding free energy, into account. We have previously established a virus-like particle (VLP) system to assess the packaging efficiency of different SARS-CoV-2 genome fragments^15^. This identified a sequence (PS9) overlapping with the *nsp15* gene as the fragment with the strongest packaging phenotype, suggesting that the dominant SARS-CoV-2 PS should be located within this fragment. We therefore focused on the three sites matching our search motif within this fragment (Fig. 1d). Only one of them can be extended, by base-pairing, into a stem-loop of similar length to the MHV PS, making it the likely candidate for the (dominant) SARS-CoV-2 PS (Fig. 1b). Mutants of PS9 (*cf*. **Mutations - Set 1** in the SI) presenting these stem-loops with different frequencies in the folding ensemble showed better packaging for mutants presenting a ‘CCAA’ motif, consistent with the SARS-CoV-2 PS identified here (Fig. S2a). Akin to the MHV PS, the sequence of the putative SARS-CoV-2 PS (Fig. 1b) corresponds to the coding sequence of an external loop in the *nsp15* gene^13^, albeit at a site further downstream (Fig. S1b), presenting a further analogy to the MHV PS. Taken together, these characteristics make this stem-loop a likely candidate for the SARS-CoV-2 PS.

### Experimental validation of the predicted SARS-CoV-2 PS

As validation of the predicted SARS-CoV-2 PS we created several mutants of this stem-loop in the context of a subsequence of PS9 containing the putative PS (the **control**; genome positions 20158-20824) and assessed their impact on packaging in the context of the VLP system^15^. Six experiments (**Mut1** – **Mut6**) correspond to in-frame truncations of different lengths, replacing all nucleotides between, and including, the encircled nucleotides (Fig. 2a) by ‘UUU’ (*cf*. **Mutations - Set 2** in the SI).

**Fig. 2.**
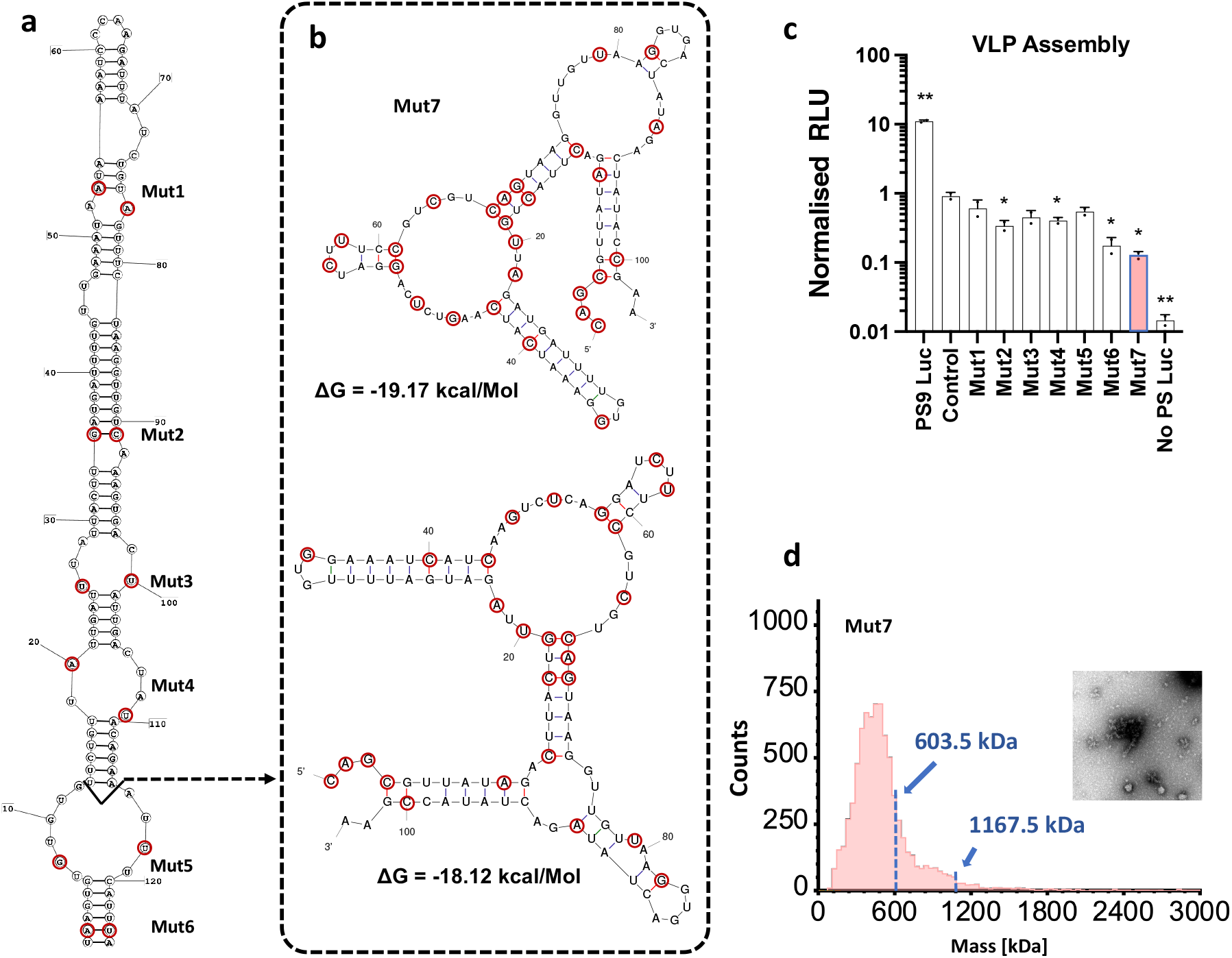
Experimental validation of the SARS-CoV-2 PS. **a** In frame truncations of the SARS-CoV-2 PS of increasing length were generated (**Mut1**-**Mut6**), replacing the sequence between, and including, the encircled nucleotides (red) by ‘UUU’. **b** Secondary structure of the PS after 27 synonymous mutations (**Mut7**, red circles), ablating the PS secondary structure as shown. **c** All mutations resulted in a reduction in packaging in the context of the VLP system^15^, with the strongest effect seen for **Mut7** (pink). Data are presented as mean +/-SD of two independent biological replicates. *, p<0.05; **, p<0.01 by two-sided Student’s t-test. **d** Mass photometry and imaging for **Mut7** shows a range of assembly products; masses corresponding to a single and double RNP complex are indicated by arrows.

The largest drop in packaging was observed for **Mut7** (Fig. 2C). In this experiment, 27 synonymous mutations were introduced to ablate the PS fold (Fig. 2b), with a mutation rate of ∼26% (27 mutations over 105 nts; indicated by red dots). However, this did not fully ablate assembly, as was also the case for the MHV PS where an equivalent experiment was performed with a mutation rate of 21% (20 in 95) to establish the function of the MHV PS^11^. This suggests that both in SARS-CoV-2 and MHV the packaging mechanism does not only rely on a single PS, as we discuss in more detail in the following sections.

As the VLP system considers packaging in the context of all structural proteins, we also performed *in vitro* assembly of N protein around the mutant sequences to interrogate the role of different mutations in RNP complex formation. Mass photometry of the nucleic acid/N-protein complexes for **Mut7** (Fig. 2d) shows a range of different assembly products, including a subspecies corresponding to the weight of an RNP complex (603.5 kDa, *cf*. Methods). The latter confirms that RNP complex formation is still taking place, albeit inefficiently, explaining why assembly has not been fully ablated in these experiments. A similar scenario is observed for **Mut1** to **Mut 6** (Fig. S7). Taken together, there seems to be a correlation between RNP complex formation and packaging, in support of the hypothesis that the dominant PS is involved in nucleation of particle formation.

### RNP complex formation and PS dynamics

The occurrence of packaging even after full truncation of the SARS-CoV-2 PS (**Mut6**) points to a complex packaging mechanism, potentially also involving other sites in the genome. To address this, we start by dissecting the role of the dominant PS in RNP complex formation, and then identify other sites that could play a similar role.

Results for MHV reveal an interplay of N- and M-protein with the PS, but the full determinants of this interaction are not known at present^10^. Cryo-electron tomography (cryo-ET) and subtomogram averaging (STA) of a SARS-CoV-2 RNP complex reveal six N protein dimers in a ring-like formation^8^. We therefore started by identifying putative N-binding sites at the appropriate spacings in the gRNA that could act in tandem with the PS. Sequence-specific binding of N protein has been probed by RNA SELEX^16^, identifying a selected library of RNA sequences with affinity for N-protein. In that reference, these sequences were interrogated for a shared sequence motif. However, as PSs are sequence/structure motifs in which sequence-specific binding motifs are typically presented in the single-stranded parts (such as bulges and loops) of the PS fold, we reanalysed the published selected library from that reference for such motifs. For this, we sampled the secondary structures of the selected library via a sliding window approach (*cf*. Methods), extracting the bulge sequences from any stem-loop, and recording on which side of the stem (5’ or 3’) they occur. We then identified any recurrent sequence motifs for the 5’, respectively 3’ side, recording their frequency of occurrence across the stem-loop ensemble. In contrast to the previous analysis^16^, this revealed a preference for an A/U-rich (bulge) sequence motif (Fig 3a).

**Fig. 3.**
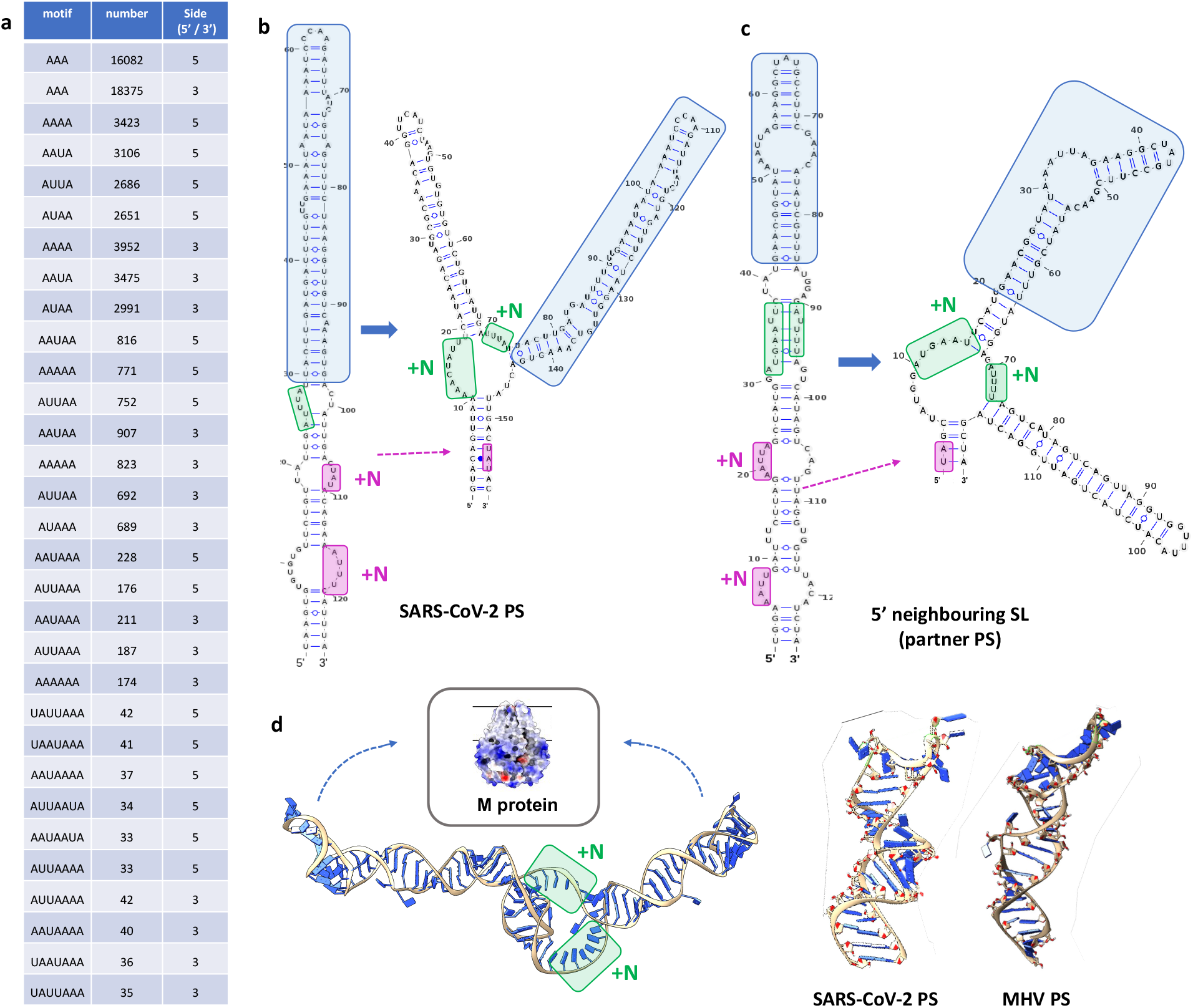
PS dynamics. **a** Analysis of published SELEX data^16^ for N-binding sequence motifs in the bulges of stem-loops in the selected library reveals a preference for A/U-rich motifs (column 1); their frequencies in the ensemble (column 2) and locations in a bulge on the 5’ or 3’ side of a stem (column 3) are also indicated. **b** The SARS-CoV-2 PS presents A/U-rich motifs in its two loops from the bottom (pink). N-protein triggers a conformational switch to a double-stem architecture presenting further N-binding motifs (green). **c** A second stem-loop, located 113 nts upstream from the PS (or 51 nts in the double-stem-loop conformation) exhibits similar characteristics, suggesting a similar role in RNP complex formation. **d** The teritiary structure of the double-loop conformation rendered with the *RNAComposer*^*19*^; N binding to the central loop (green boxes) could reorient the stem-loops towards M-protein in the membrane. The tertiary structure of the SARS-CoV-2 PS in the long conformation exhibits similar structural characteristics to the MHV PS (right); its loop is the likely contact with M-protein^18^, its receptor.

The two bottom-most bulges of the SARS-CoV-2 PS present such A/U-rich motifs at the 3’ side of the stem (Fig 3b; pink boxes). In order to determine the likely change in secondary structure of the PS when those areas are engaged in N binding, we performed a constrained secondary structure prediction in which the sequence fragment encompassing both alleged N-binding sequences (pink) is constrained to be unavailable for base-pairing^17^. This resulted in the double stem-loop conformation (right), which differs in folding free energy by only 8.9kcal/M (-26.1 compared to -35), consistent with reasonable estimates for the binding free energy of N protein. The dimensions of this alternative fold are in excellent agreement with the imaged RNP complex^18^ (*cf*. Fig. S6A in this reference), which reveals an internal density of 16nm in height, corresponding to approximately 53 bps (assuming ∼0.3 nm/bp). Interestingly, the lower part of this density was assigned to the N-terminal domain of N protein (N_NTD)^18^. Our secondary structure fold reveals further N-binding motifs in the loop connecting the two stem-loops (Fig. 3b, green box), consistent with this observation. Tertiary structure prediction of the double stem-loop conformation using *RNAComposer*^*19*^ (Fig. 3d, left) shows the two stem-loops pointing outwards. It is possible that N-binding to the sites indicated by green boxes flips them into a parallel conformation pointing upwards (as indicated by the arrows), matching the RNA density seen at the centre of the RNP complex. In summary, the double stem-loop conformation likely corresponds to the internal density in the RNP complex, anchoring it at the inner membrane via contact with M protein at the loop portion of one or both of the SLs in the double stem-loop fold (Fig. 3d, right).

A computational search for a second stem-loop of similar size and with similar structural characteristics, that can also undergo this conformational change in response to N protein binding, located such a stem-loop (Fig. 3c, the ‘partner PS’) at a distance of 51 nts 5’ from the (dominant) PS in its double-loop conformation. Comparison with RNA secondary structure elements identified previously via icSHAPE^20^ structure probing of the SARS-CoV-2 genome (*cf*. Fig. 3 in that reference) reveals a match with the double stem-loop conformation of the partner PS. It also indicates a double stem-loop with a slightly different fold in the position of the dominant PS. Interestingly, the reactivities indicated are also consistent with our variant of the fold in this area (the double stem-loop structure in Fig. 3b), confirming that our results are consistent with secondary structure predictions in the literature.

A computational search for a second stem-loop of similar size and with similar structural characteristics, that can also undergo this conformational change in response to N protein binding, located such a stem-loop (Fig. 3c, the ‘partner PS’) at a distance of 51 nts 5’ from the (dominant) PS in its double-loop conformation. Comparison with RNA secondary structure elements identified previously via icSHAPE^20^ structure probing of the SARS-CoV-2 genome (*cf*. Fig. 3 in that reference) reveals a match with the double stem-loop conformation of the partner PS. It also indicates a double stem-loop with a slightly different fold in the position of the dominant PS. Interestingly, the reactivities indicated are also consistent with our variant of the fold in this area (the double stem-loop structure in Fig. 3b), confirming that our results are consistent with secondary structure predictions in the literature.

As the published secondary structures are based on gRNA in the absence of protein, we also interrogated the structure of the RNP complexes in the *in vitro* experiments using X-ray synchrotron footprinting, a technique we pioneered for the analysis of packaged viral genomes^21^. Structure prediction based on these data is complicated by the fact that low reactivity values are indicative of both N-binding and RNA base-pairing, which are difficult to deconvolute in the context of an RNP complex. However, a contiguous stretch of low reactivities is indicative of the *location* of an RNP complex. The XRF results for transcript in complex with N-protein (Fig. S4) not only reveal low reactivities in a 264-nt long fragment encompassing the PS, but also in an adjacent fragment upstream where the partner PS is located. This is indicative of additional N-binding events around the partner PS, suggestive of a double RNP complex.

### Cooperative action of the dominant and partner PS

To better understand the cooperative action of the dominant PS and its partner, which together form the likely nucleation site for packaging and particle assembly, we created a set of mutants (*cf*. **Mutations - Set 3** in the SI) that explore the contributions of different aspects of the PS dynamics (Fig. 3) to packaging and virion assembly. We created mutants stabilising the PSs in their extended form, preventing their transition into the double stem-loop conformation (Fig. 4b, **X3**). We also ablated the double stem-loop fold of both PSs (**X7**) and mutated the regions containing the N-binding motifs not overlapping with the PSs (**X8**). All three scenarios impacted on packaging compared with control (**X1**), confirming the importance of these different aspects of RNP complex formation for gRNA packaging (Fig. 4a). We also used mass photometry to characterize RNP complex formation for the different scenarios. This revealed the presence of many other species over and above a single and double RNP complex, supporting the hypothesis that packaging efficiency is linked with successful RNP complex formation.

**Fig. 4.**
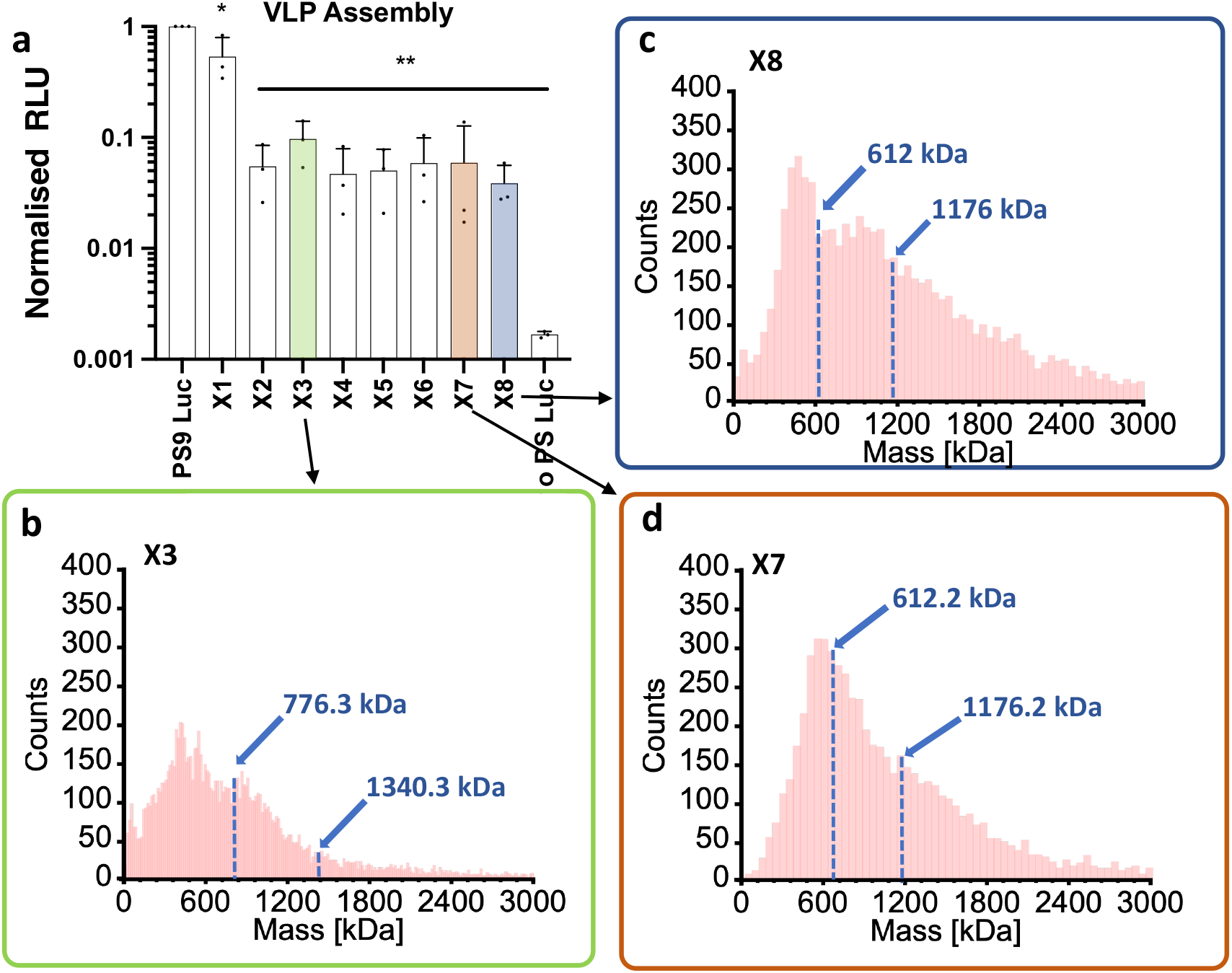
Cooperative action of the dominant and partner PS. **a** Mutations performed in the control fragment (**X1**) reduce packaging in the VLP system^15^: PSs stabilized, preventing dynamics (**X3**); dominant and partner PS mutated (**X7**); exterior N-binding motifs mutated (**X8**). Data are presented as mean +/-SD of two independent biological replicates. *, p<0.05; **, p<0.0001 by two-sided Student’s t-test. **b-d** Mass photometry reveals a local peak around the mass corresponding to a single RNP complex for X3 and X7, in contrast to X8, consistent with their superior performance in packaging.

### Formation of a double RNP complex

To develop a model of complex formation around the dominant PS and its partner, we next profiled the locations of putative N-binding sites, identifying A/U-rich motifs consistent with the sequence determinants of the binding sites. Using a sliding window approach, we located all contiguous 20nt-long gRNA fragments with over 75% A/U content along the gRNA (lollipops in Fig. 5a), identifying a cluster of such sites in proximity to the PS and its partner (Fig. 5b). Their positions relative to these two stem-loops, shown here schematically in the double-stem conformation as a ‘Y’ shape (with the stem-loop corresponding to the extended form of the PS indicated by an asterisk), reveals 12 putative N-binding sites (Fig. 5c & green highlights in Fig. S5). If each such site was associated with an N protein dimer, this would be consistent with the formation of two RNP complexes around these two stem-loops as illustrated in the cartoon (Fig. 5d). Cryo-EM of N-protein assembling around the control (*cf*. the ‘long fragment’ in Fig. 5c) shows evidence of such double-RNP complex formation in support of this hypothesis (Fig. 5e & Fig. S6). Mass photometry of this fragment (Fig. 5f) moreover shows a signal at 1339.7 kDa, the expected mass of the double RNP complex (see Methods) consistent with this interpretation. By contrast, the corresponding signal (1176 kDa) is largely absent in the shorter fragment encompassing only five N-binding sites around the dominant PS (Fig. 5g). In this case, mass photometry reveals larger numbers of single RNP complexes (at 612 kDa), with a peak at intermediates with five N-protein dimers as expected given that the fragment covers only 5 of the identified N-binding sites. These mass photometry results demonstrate that our identification of the N-binding sites and their roles in RNP complex formation matches the masses of the structures seen, in support of our model of RNP double complex formation.

**Fig. 5.**
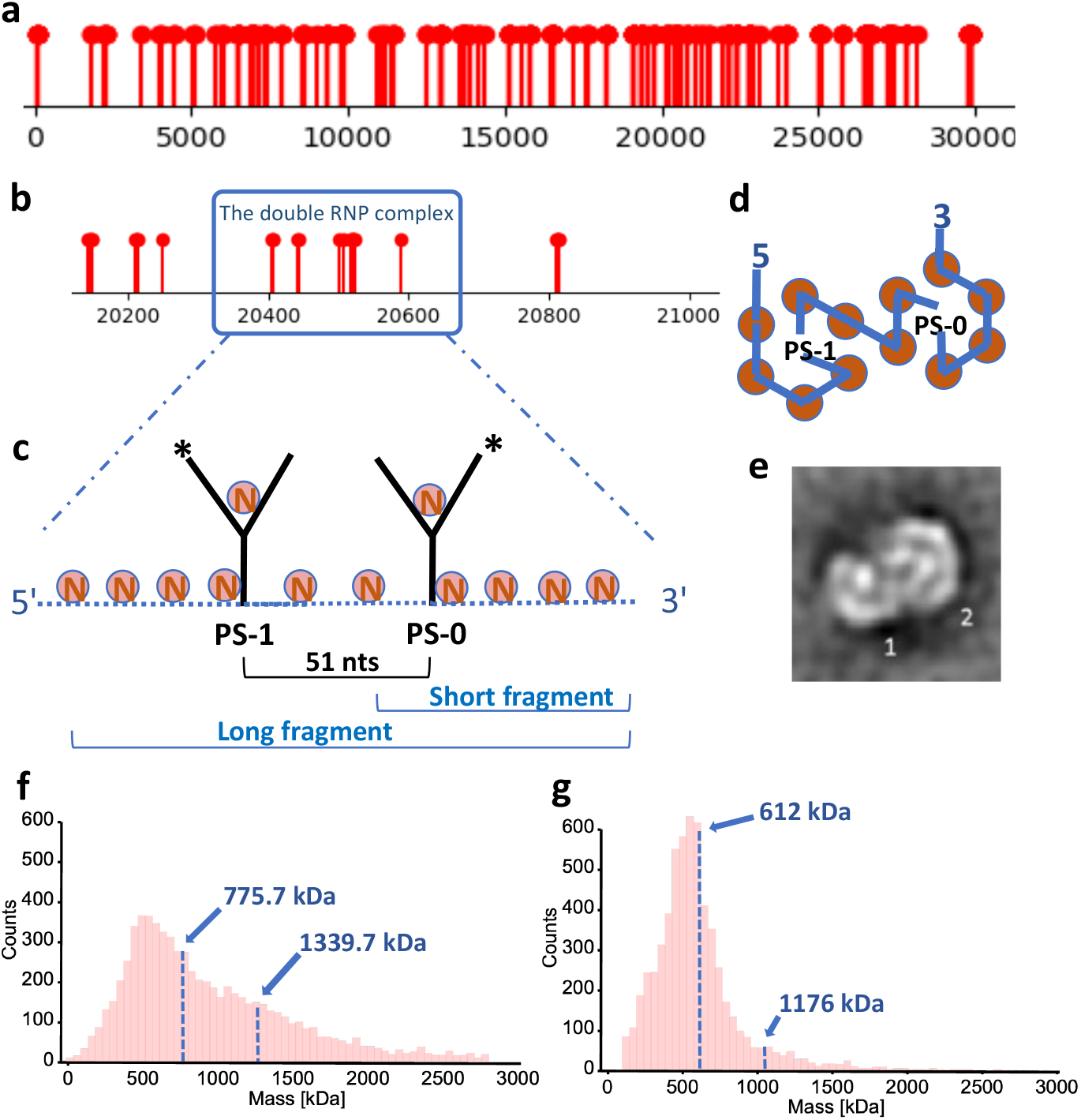
Nucleation Complex Formation. **a** Computational search for contiguous 20nt-fragments with over 75% A/U content identifies multiple locations across the genome, and **b** a cluster of such sites in proximity of the dominant PS and its partner. **c** Schematics showing the positions of these clusters (encircled Ns) in context of the two SLs in the double-loop conformation (Y shapes); the stem-loop corresponding to the top of the PS in the extended conformation is indicated by a star. **d** Schematic of how the unit in **c** could organize into a double RNP complex, if each site was associated with an N-protein dimer, matching negative stain 2D class average of this fragment in complex with N-protein (**e**). **f** Mass photometry of the longer fragment, and **g** of a shorter fragment, overlapping with the unit as indicated in **c**; the masses corresponding to a single and double RNP complex are indicated by arrows.

### Genome-wide cooperativity indicates a multiple packaging signal-mediated packaging mechanism

The cooperative action of the dominant PS with its partner is reminiscent of the multiple packaging signal mediated assembly mechanism in smaller RNA viruses^5^. As cryo-ET of SARS-CoV-2 virions reveals multiple RNA complexes in contact with the inner membrane in similar orientations^18^, it is likely that they are anchored at the membrane in similar ways. This suggests that there could be other PS-like sites along the genomic RNA that support the formation of RNP complexes and their ‘anchoring’ to M-protein akin to the mechanism characterised above, suggestive of a genome-wide, multiple packaging signal mediated packaging mechanism (Fig. 6a).

**Fig. 6.**
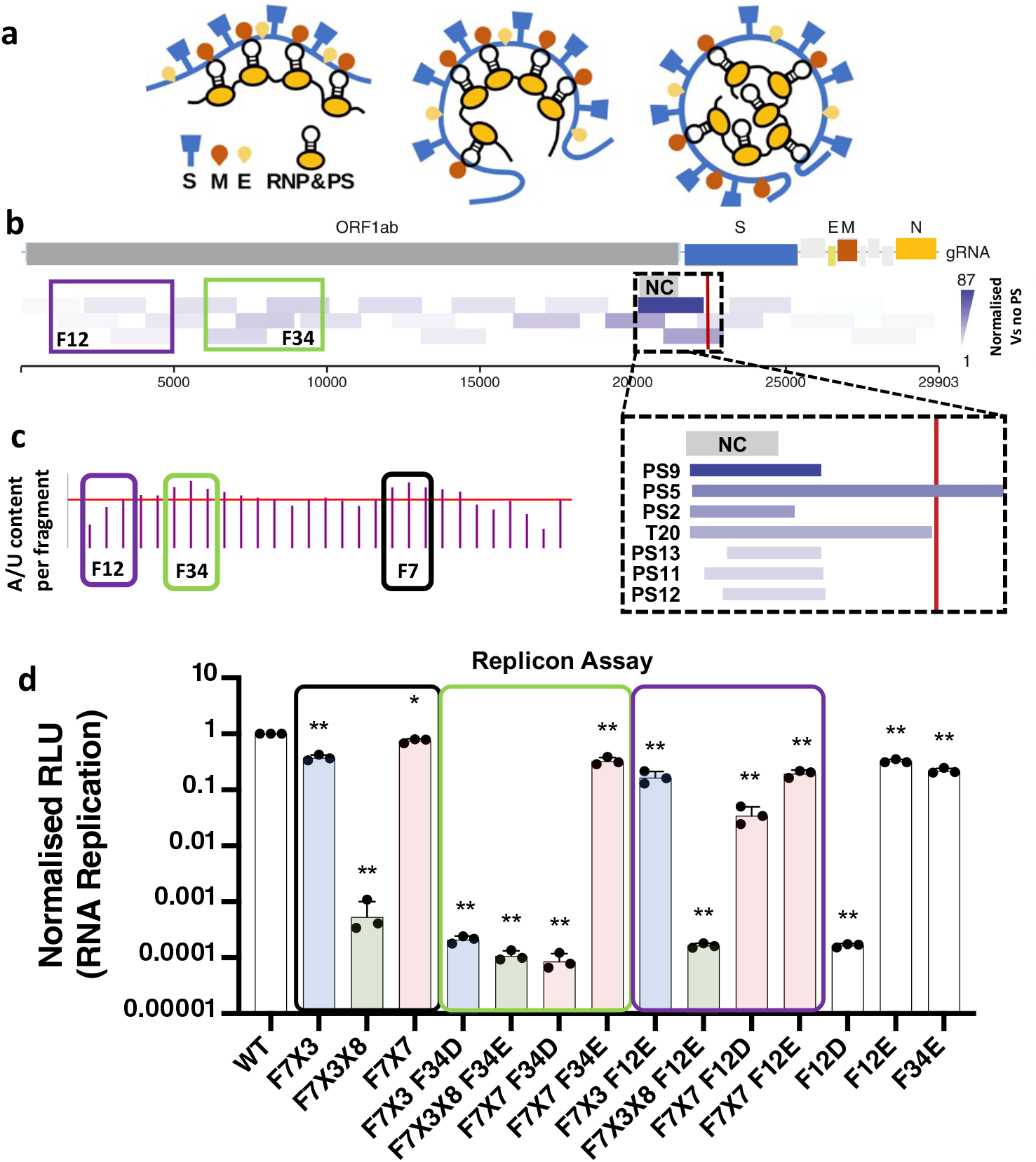
Genome-wide cooperativity of the PS-mediated packaging mechanism. **a** Cartoon illustrating how multiple dispersed RNP complexes could be anchored at the inner membrane via multiple PS containing RNP complexes. **b** 2000nt-long fragments covering the SAR-CoV-2 genome exhibit different packaging efficiencies (indicated as shades of purple, adapted from Ref. ^15^); the dashed black box indicates the three fragments with the highest packaging efficiency, with the middle one (‘T20’, darkest shade) corresponding to the strongest packaging phenotype. The inset shows relative packaging efficiencies of truncations and extensions of this fragment, including PS9, together with the location of the nucleation cassette (NC). The occurrence of another PS candidates at nt position 22244 (*cf*. Table S3) explains the superior performance of PS5 over T20. **c** Fragments exhibiting higher packaging efficiency (F34) also have above average A/U content. **d** Mutations performed in the replicon systems: X referring to the experiments in Fig. 4a, and F34 and F12 covering 4026 nts long fragments each, with ‘U’s replaced by ‘G’s for the dyscoded (D) version and ‘U’s maximised in the encoded (E) version. Data are presented as mean +/-SD of two independent biological replicates. *, p<0.05; **, p<0.0001 by two-sided Student’s t-test.

To probe cooperativity of the double RNP complex – the ‘nucleation cassette’ (NC) – with other sites in the genome, we refer to our previous study of packaging efficiency of different genomic fragments^15^. Fig. 6b, adapted from this reference, shows packaging efficiencies of different genomic fragments as shades of dark purple, with darker shades indicating higher packaging efficiency. The fragment overlapping with the dominant and partner PS (dashed box; fragment F7) shows the highest packaging, but a local maximum of lower peak height occurs in fragments F3 and F4 (green box; F34). As the latter is in an area where stem-loops with PS characteristics have been characterised in some *Embecoviruses*^*9*^, we reason that there could be secondary sites important for packaging also in the *Sarbecoviruses*. To test this in a scenario as close as possible to a wild-type infection, we now work with the full replicon system rather than the VLP system (*cf*. **Mutations - Set 4** in the SI). In the replicon system, the Spike coding sequence is replaced with secreted nanoluciferase and EGFP allowing single-round replication assays^22^. Mutant replicon plasmids are co-transfected with a Spike expression vector to produce single-round SARS-CoV-2 particles that are then used for infection of receiver cells. This assay therefore enables assessment of packaging efficiency and replication across the entire genome.

Before testing cooperativity between the nucleation cassette in F7 and putative PS sites in F34, we first repeat the packaging experiment ablating only the dominant PS and its partner in F7 in the context of the replicon system to establish a benchmark for comparison (Fig. 6d, black box). This revealed the strongest reduction in packaging efficiency for a combination of **X3** and **X8** (*cf*. F7X3X8), which simultaneously prevents formation of the double stem-loop conformation in the dominant PS and its partner and ablates the N-binding motifs. Preventing formation of the double stem-loop conformation in isolation (*cf*. F7X3) or ablating the PS folds in isolation (*cf*. F7X7) also impacts on packaging, but to a much smaller extent.

To test cooperativity between the nucleation cassette in F7 and additional sites in F3/4, we note that both fragments show the highest percentage of A/U content (Fig. 6c), as expected from the A/U rich nature of the N-binding motif. Therefore, instead of targeting the PS itself, we are mutating the N protein binding sites associated with RNP complex formation. For this, we either minimised or maximised the occurrence of the A/U motifs as follows. A/U-rich sites were ‘dyscoded’ (F34D) by replacing ‘U’s with ‘G’s synonymously were possible, thus eliminating the N-binding sites. We also ‘encoded’ (F34E) more binding sites by maximising the occurrence of ‘U’s synonymously, thus ablating the position-specific encoding of the N-binding sites via the generation of additional sites. We then combined these mutations (F34D or F34E) with those in the control experiments in F7. All combinations of mutations in F7 and F34 resulted in a decrease in packaging efficiency (green box), demonstrating cooperativity between the nucleation cassette in F7 and sites in F34. The strongest drop in packaging efficiency was seen in the dyscoded case (*cf*. F7X7F34D), compared with only a small decrease for the encoded variant (*cf*. F7X7F34E). This shows that ablating binding sites is more detrimental to packaging than creating additional ones. As a control, we also created analogous synonymously dyscoded and encoded versions of a genomic fragment further upstream (F12; purple box), that has the same length as F34, but has lower packaging efficiency (Fig. 6b) and A/U content (Fig. 6c). In this case, combining F12D with F7X7 reduced packaging significantly less than in combination with F34D (*cf*. F7X7F12D), demonstrating that sites in F34 are more important for packaging than those in F12. Taken together, this indicates cooperativity between the nucleation cassette in F7 and putative N-binding sites in F34 in packaging, in support of a multiple packaging signal mediated assembly and genome packaging mechanism in SARS-CoV-2.

## Discussion

Our study has located the *Sarbecovirus* PS in the coding region of an external loop of the *nsp15* gene, downstream from, and in proximity to, the known *Embecovirus* PS. It also identified a second stem-loop with similar structural characteristics (the partner PS) upstream from the dominant PS and provided evidence for their cooperative action in the formation of a double RNP complex. Our data characterise the dynamic action of the PS as a two-stage process, with an N-binding mediated conformational change from the extended to a double stem-loop conformation. It is possible that the two stem-loops in the double stem-loop conformation each contact M protein, bringing two copies of M protein into close proximity and defining their relative orientation. This would create a specific intrinsic curvature between adjacent M proteins in the envelope. It had previously been demonstrated that such intrinsic curvature is essential for virus budding^23^, pointing towards another function for the double stem-loop conformation beyond RNP complex formation.

The double RNP complex around the dominant PS and its partner could explain the superior packaging efficiency of genomic fragments overlapping with this nucleation cassette^15^. Ablating formation of the double RNP complex in isolation does not fully ablate virion formation in the context of the replicon system, echoing previous experiments in MHV^11^. We show here that this is due to cooperative action with other sites in the SARS-CoV-2 genome. Shorter stem-loops with PS-like characteristics have previously been reported in *Embecoviruses* in a similar region of the RbCoV genome^10^, but their cooperativity with the dominant packaging signal has not been investigated. Our study is thus the first report of a multiple packaging signal-mediated assembly/packaging mechanism in coronaviruses, providing evidence for genome-wide cooperativity in genome packaging and virion assembly.

A previous study displacing the dominant PS in MHV demonstrated its functionality at an ectopic site in the genome, further supporting the hypothesis that PS function is independent of location^11^. This study also showed that PS containing fragments have a selective packaging advantage over sequences lacking a PS^11^. Notably, both the double RNP complex and the additional sites upstream are located in coding regions of the non-structural proteins. This favours the highly selective packaging of the full-length gRNA over the abundant subgenomic fragments overlapping with the coding regions of the structural proteins, thus reducing the occurrence of incompletely packaged particles. A recent study reported a structured RNA element within *nsp15* that contributes to SARS-CoV-2 genome encapsidation, but the actual PS structure and mechanism of function remained unidentified^24^. This 667 nt long sequence overlaps with one of our motifs in Fig. S3. We established (Fig. S9) that this fragment can fold into a double-stem loop configuration containing a stem-loop overlapping with one of our motifs, providing further validation for our analysis.

The detailed mechanistic insights afforded by our study open up novel opportunities in medicine and bionanotechnology. They point to novel drug targets for small molecular weight compounds that could act as assembly/packaging inhibitors^25^. A previous study identified highly stable RNA secondary structures in the SARS-CoV-2 genome that lend themselves as potential drug targets for RNA binding small molecules^26^. RNA structure probing moreover identified RNA secondary structure with significant covariation among SARS-CoV-2 strains and other coronaviruses. One of these is our partner PS in the double stem-loop conformation^27^. Secondary structure-restrained 3D modelling of its structure identified a putative druggable pocket (Fig. S11 in that reference). Based on our results, we hypothesize that molecules targeting this and equivalent pockets in multiple dispersed PSs could be potent assembly inhibitors.

Our analysis also suggests novel ways of engineering membrane-based drug delivery systems for large mRNA cargoes. In particular, the determinants of RNP complex formation and additional PS sites could be superimposed on guest RNAs to improve their packaging into glycoprotein-studded membranes. Such systems could offer extensions beyond the canonical AAV and lentiviral-based vector systems in gene therapy towards the delivery of large genetic cargoes.

## Acknowledgement

RT acknowledges funding via an EPSRC Established Career Fellowship (EP/R023204/1), that also funded her two visits to the Gladstone Institutes in 2024, and a Royal Society Wolfson Fellowship (RSWF\R1\180009). RT & PGS also acknowledge support from the Joint Wellcome Trust Investigator Awards (110145, 110146, 224509/A/21/Z). M.O. received support from the Roddenberry Foundation, the James B. Pendleton Charitable Trust, and P. and E. Taft, and is supported by the Gladstone Institutes. M.O. is a Biohub San Francisco Investigator and the Nick and Sue Hellmann Distinguished Professor. This research used resources of the National Synchrotron Light Source II, a U.S. Department of Energy (DOE) Office of Science User Facility operated for the DOE Office of Science by Brookhaven National Laboratory under Contract No. DE-SC0012704. RT would like to thank the Ott Lab for their wonderful hospitality during her two extended visits to the Gladstone Institutes (and for looking well after her scooter). We thank Prof Fred Antson and his team for the provision of SARS-CoV-2 N-protein for the experiments carried out in Leeds. We would like to thank Louie Aspinall, Yehuda Halfon and Neil Ranson at University of Leeds for help with Astbury Bioscience Electron Microscopy facility access and cryo-EM data collection.

## Methods

### Cells

BHK-21, 293T cells were obtained from ATCC and grown in standard media: DMEM (Corning) supplemented with 10% fetal bovine serum (FBS) (GeminiBio), 1x Glutamax (Corning), 1x non-essential amino acids (NEAA) (Corning), and 1x penicillin-streptomycin (Corning) at 37 °C, 5% CO_2_. Vero cells stably expressing human ACE2 and TMPRSS2 (VAT) (gifted from A. Creanga and B. Graham at NIH) were maintained in standard media with the addition of 10 μg/mL of puromycin. All cell lines are confirmed mycoplasma-free by quarterly testing.

### SARS-CoV-2 VLPs

VLP experiments were conducted as described previously with minor modifications^15^. Briefly, 5.0x10^5^ HEK293T cells were seeded in 6-well plates. The next day, cells were transfected with 500 ng packaging signal luciferase construct, 333 ng N expression vector, 166 ng M IRES E expression vector, and 10 ng Spike expression vector using X-tremegene 9 transfection reagent (Sigma-Aldrich) according to manufacturer’s recommendations. After 4-6 hours, the media was changed with 1.5 mL fresh growth medium. After 72 hours, the supernatant was collected and 0.2 μm filtered. 250 μL of supernatant was mixed with 4.0x10^4^ VAT cells in white 96-well plates and incubated for 24 hours. The cells were then washed once with PBS and lysed with 20 μL passive lysis buffer (Promega) for 15 minutes at room temperature. 50 μL of reconstituted luciferase assay reagent was added and luminescence measurement recorded on a Tecan plate reader.

### SARS-CoV-2 replicons

Replicon plasmids were constructed as described previously with some modifications.^22^ Briefly, encoded and dyscoded fragments were purchased as double-stranded DNA from Twist Biosciences and cloned into genome fragments using HiFi DNA assembly master mix (NEB). The fragments were then assembled using Golden Gate assembly into full length replicons, amplified, and sequenced using whole plasmid sequencing from Plasmidsaurus. Replicon assays were conducted as described previously.^28^

### Mass photometry

In vitro SARS-CoV-2 nucleocapsid (N) protein - RNA binding reactions were performed in a buffer containing 25 mM HEPES (pH 7.5), 250 mM NaCl, 2 mM TCEP, 0.01% Tween 20, and 1 U murine RNase inhibitor (New England Biolabs), using 10 nM RNA and 200 nM N protein in a total volume of 100 µL and incubated at 37 °C for 30 min. RNA was heat annealed in 25 mM MES (pH 7.0), 50 mM NaCl by heating to 70 °C for 10 min followed by cooling to room temperature. For transmission electron microscopy, 3 µL of sample was applied to glow discharged, formvar coated copper 300 mesh grids (Agar scientific), negatively stained with 2% (w/v) uranyl acetate, and imaged on a Talos 120C microscope at 56,000x magnification. In parallel, particle mass distributions were measured by mass photometry using a Refeyn OneMP instrument (Refeyn Ltd.); binding reactions were diluted 1:60 into filtered phosphate buffer immediately prior to acquisition on cleaned glass coverslips and data was collected for 1 min. Data were analysed using Discover software (Refeyn Ltd.) to resolve mass populations corresponding to SARS-CoV-2 N protein:RNA assemblies, and extracted and plotted using custom scripts leveraging the Matplotlib python library. Scripts can be found at (https://github.com/nikesh86-2/mass-photometry-gaussian-analysis). RNA masses were calculated using the oligonucleotide analyser from Integrated DNA Technologies (https://eu.idtdna.com/calc/analyzer). The N protein masses were calculated using https://web.expasy.org/protparam/, inputting the N protein sequence for the SARS-Cov-2 reference strain described. The total masses for protein:RNA complexes are predicted in table X, assuming a potential 1, 2, 6, 12 or 24 N proteins to 1 RNA ratio.

### SELEX analysis

The selected library from the 3^rd^ round of SELEX (SARS-CoV-2 wildtype)^16^ was reanalysed by us for motifs occurring in the single-stranded portions of any stem-loops presented in local folds of 150 nt long segments. For this, all stem-loops with a free energy of below -4 kcal/mol were identified and their bulge sequences recorded. For each bulge with a length of up to 7 nts on either flank, bulge sequences occurring with a frequency >75% of the maximal frequency bulge for that length were recorded. This revealed a preference for A/U-rich bulge motifs (Fig. 3a).

### Sample preparation for in vitro experiments

Mutant and control RNAs (cf. list of sequences in the SI) was pre-annealed in 25 mM MES pH 7.0, 50 mM NaCl at 70°C for 10 minutes, and then cooled to room temperature. Binding assays were carried out with a buffer containing 25 mM HEPES at pH 7.5 with 250 nM NaCl, 2 mM TCEP, 1 unit RNase inhibitor (Murine), and 0.01% Tween20. Concentration in 100 µL was 10 nM RNA and 200 nM N Protein. These were incubated for 30 minutes at 37 °C. For the imaging, 3 ul of sample were applied to formvar coated, glow discharged copper grids and stained using 2% (w/v) uranyl acetate. The sample was imaged using a Talos 120C microscope at 56,000x magnification.

### Cryo-EM analysis of RNPs

3 µl of reconstituted SARS-CoV-2 N with RNA (10nM RNA, 200nM N protein was applied to glow-discharged holey lacey grids (Agar Scientific) and plunge frozen in liquid ethane using Vitrobot IV (thermos scientific). The cryo-EM grids were imaged on Krios1 equipped with Falcon4i + Selectris, with a pixel size of 0.74 Å/pix. 3000 movies were collected that were pre-processed using RELION 4. Movies were motion corrected to generate aligned micrographs (using relion motioncorr implementation) and CTF estimation performed using CTFFIND4. From 500 micrographs, a subset of particles was manually picked to train Topaz particle picker model, and the resulting trained model was used to pick 122,455 particles. Particles were boxed and 2D class averages were generated with 100 iterations and 20 class groups, from which distinct classes assigned as either ‘doubles’ or ‘singles’ were selected for further interpretation.

### X-ray synchrotron footprinting

XRF experiments were as described previously^21^. Samples were exposed in duplicate to X-ray pulses of 50 ms at the National Synchrotron Light Source II, beamline 17-BM XFP at Brookhaven National Laboratory (Upton, NY). Nucleotide modification propensity (reactivity) is directly related to residue mobility and thus reflects base pairing and inter-molecular contacts. Reactivity was quantified by capillary electrophoresis sequencing using dye-labeled primers, and data analysed using purposed designed algorithms^21^ based on the QuShape software^29^. Pairwise Pearson correlation coefficients (PCCs) for normalized replicates at 50 ms exposure were 0.927 for transcript and 0.951 in the presence of N-protein for a duplicate run.

