## Supplementary Material for "Pandemic Coronavirus Genome Packaging Relies on Multiple Dispersed Packaging Signals"

#### Supplementary Figures

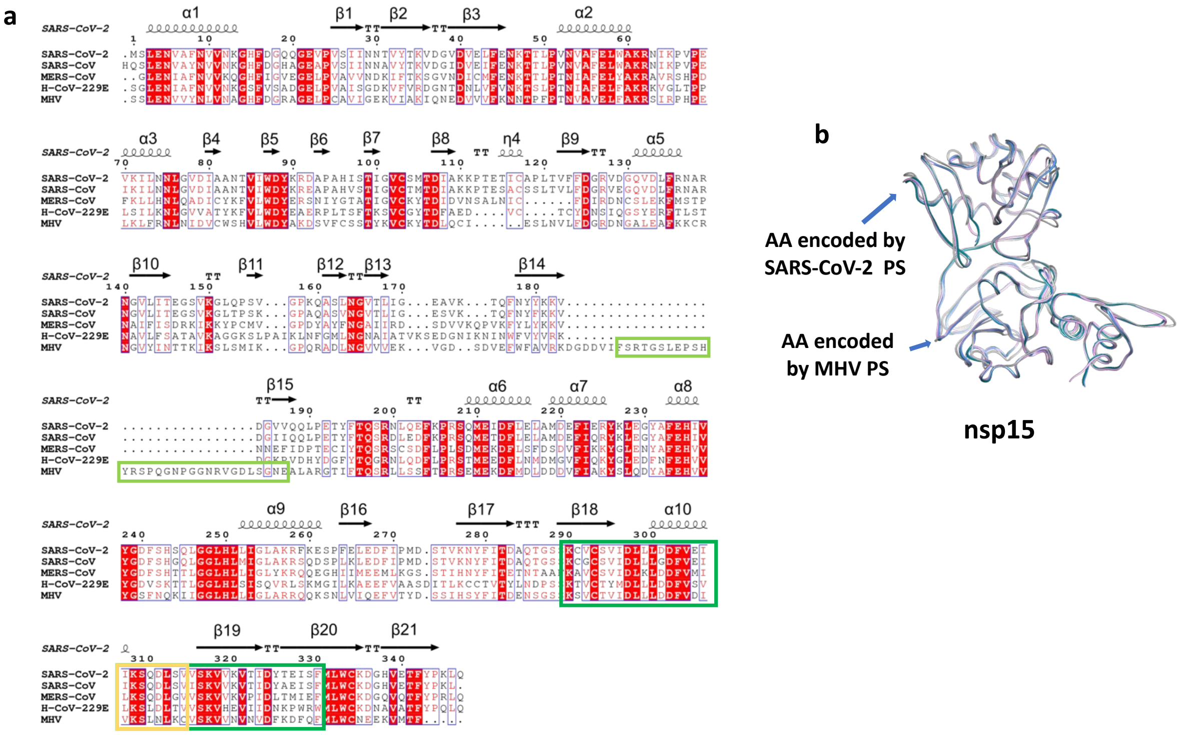


**Fig. S1 | Location of the SARS-CoV-2 PS in an alignment of different coronavirus nsp15 coding sequences. a** The MHV PS (light green box, superimposed on a published alignment) is located in an area of the genome where the other coronaviruses have a deletion. The SARS-CoV-2 PS identified here is located further 3' (dark green box). **b** The amino acids encoded by the PS sequences correspond to surface loops of nsp15.

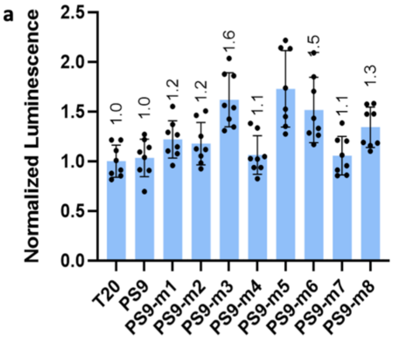


b

| SL | Folding propensity | | | | | | | | |
|---|---|---|---|---|---|---|---|---|---|
| | PS9 | PS9-m1 | PS9-m2 | PS9-m3 | PS9-m4 | PS9-m5 | PS9-m6 | PS9-m7 | PS9-m8 |
| 1 | 372 | 12 | 15 | 334 | 20 | 179 | 193 | 21 | 25 |
| 2 | 2 | 2 | 4 | 1 | 0 | 2 | 6 | 3 | 0 |
| 3 | 111 | 115 | 0 | 0 | 1 | 121 | 125 | 0 | 1 |

**Fig. S2 | Frequencies of different stem-loops (SLs) with a 'CAA' loop motif in different mutants. a** Packaging efficiencies of different sequences in *Mutant Set 1* established in the VLP system. **b** Frequencies of SL1-3 (*cf.* Fig 1) in the folding ensemble of different mutant sequences.

| Motif 'CCAA' | | |
|---|---|---|
| 5990, 'AGCAACCAATTGAT' | 9066, 'TGATACCAATGTAC' | 12657, 'AAGGGCCAATTCTG' |
| 20538, 'AAATCCCAAGATTT' | 22244, 'ATTTGCCAATAGGT' | 25607, 'ACTCTCCAAGGGTG' |
| 26170, 'TGGAACCAATTTAT' | 28393, 'CGGCCCCAAGGTTT' | 29297, 'AAGATCCAAATTTC' |
| **Motif 'ACAA'** | | |
| 3269, 'TTCAAACAATTGTT' | 5862, 'TTTCTACAAAGAAA' | 5949, 'GTTGGACAATTATT' |
| 6815, 'GAAGTACAAATTCT' | 7375, 'CTTGTACAAATGGC' | 9279, 'ACTTAACAATGATT' |
| 13309, 'TATGTACAAATACC' | 13687, 'AAGAAACAATTTAT' | 18534, 'ATTGTACAAATGTT' |
| 18768, 'GTTCAACAATGGGG' | 20214, 'AATTTACAAGAATT' | 24661, 'CTTGGACAATCAAA' |
| 26217, 'TAAGCACAAGCTGA' | 27160, 'CAGTGACAATATTG' | 29482, 'TCCAAACAATTGCA' |
| **Motif 'xCCAA'** | | |
| 3124, 'GATTACCAAGGTAA' | 3441, 'TGCAGCCAATGTTT' | 10443, 'GAGGCCCAATTTCA' |
| 15539, 'CACGGCCAATGTTA' | 20172, 'GTTGTCCAACAATT' | 25607, 'ACTCTCCAAGGGTG' |
| 26170, 'TGGAACCAATTTAT' | 27938, 'TTTCACCAAGAATG' | 28404, 'TTTACCCAATAATA' |
| **Motif 'xACAA'** | | |
| 21, 'AGGTAACAAACCAA' | 1512, 'CCATAACAAGTGTG' | 3229, 'GGTCAACAAGACGG' |
| 3582, 'CGGACACAATCTTG' | 5819, 'TACTTACAAAGTCC' | 6713, 'TTAGTACAACTACT' |
| 7497, 'GTGTTACAAACGTA' | 10118, 'GTGGTACAACTACA' | 11943, 'GTTACACAATGACA' |
| 12066, 'GCTGGACAACAGGG' | 12350, 'TGCAGACAATGCTT' | 13687, 'AAGAAACAATTTAT' |
| 14681, 'TTTTAACAAAGACT' | 19353, 'TTAAAACAATTACC' | 21554, 'AACGAACAATGTTT' |
| 26570, 'CTTGAACAATGGAA' | 27542, 'TGATAACAAATTTG' | 29070, 'AGCATACAATGTAA' |
| 29762, 'AGTGAACAATGCTA' | | |

**Fig. S3 | Identification of stem-loops matching the search motif in Fig. 1.** Position of every 17-nt sequence matching the search motif, organized by loop motif. Sequences with loops closed by 4 consecutive base-pairs are shown in orange.

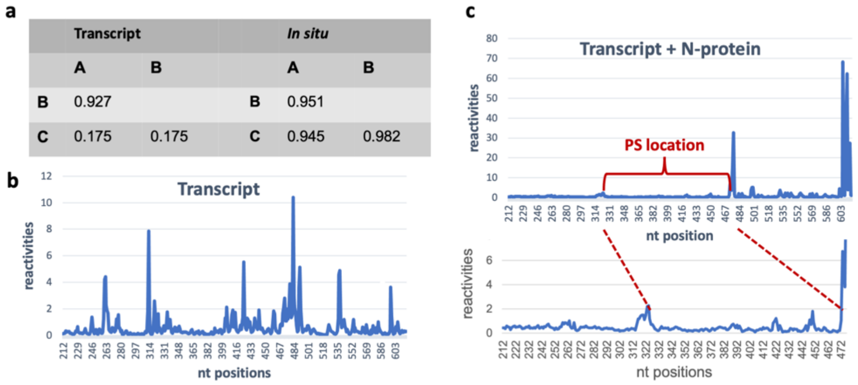

| | Transcript | | | *In situ* | |
|---|---|---|---|---|---|
| | A | B | | A | B |
| B | 0.927 | | B | 0.951 | |
| C | 0.175 | 0.175 | C | 0.945 | 0.982 |



**Fig. S4 | X-ray synchrotron footprinting (XRF) of the long, PS containing, fragment in complex with N protein. a** Pairwise Pearson correlation coefficients (PCCs) for normalized replicates at 50 ms exposure from triplicate runs show an excellent correlation for samples A and B in the N protein free case (transcript), and of all samples in the presence of N protein (*in situ*). Data were processed following protocols established earlier (*cf.* Ref. 15). **b** The reactivities for transcript. **c** Reactivity values are low in the sequence corresponding to the dominant PS, indicating a combination of RNA base-paring and N protein binding.

GGTTTACAACCATCTGTAGGTCCCAAACAAGCTAGTCTTAATGGAGTCACATTAATTGGAGAAGC

CGTAAAAACACAGTTCAATTATTATAAGAAAGTTGATGGTGTTGTCCAACAATTACCTGAAACTTAC

TTTACTCAGAGTAGAAATTTACAAGAATTTAAACCCAGGAGTCAAATGGAAATTGATTTCTTAGAAT

PS-1 TAGCTATGGATGAATTCATTGAACGGTATAAATTAGAAGGCTATGCCTTCGAACATATCGTTTATGG

AGATTTTAGTCATAGTCAGTTAGGTGGTTTACATCTACTGATTGGACTAGCTAAACGTTTTAAGGAA

TCACCTTTTGAATTAGAAGATTTTATTCCTATGGACAGTACAGTTAAAAACTATTTCATAACAGATGC

PS 0 GCAAACAGGTTCATCTAAGTGTGTGTGTTCTGTTATTGATTTATTACTTGATGATTTTGTTGAAATAA

TAAAATCCCAAGATTTATCTGTAGTTTCTAAGGTTGTCAAAGTGACTATTGACTATACAGAAATTTCA

TTTATGCTTTGGTGTAAAGATGGCCATGTAGAAACATTTTACCCAAAATTACAATCTAGTCAAGCGT

GGCAACCGGGTGTTGCTATGCCTAATCTTTACAAAATGCAAAGAATGCTATTAGAAAAGTGTGACC

TTCAAAATTATGGTGATAGTGCAACATTACCTAAAGGCATAATGATGAATGTCGCAAAATATACTCA

ACTGTGTCAATATTTAAACACATTAACATTAGCTGTACCCTATAATATGAGAGTTATACATTTTG

**Fig. S5 | Locations of the N-binding sites associated with the double RNP complex.** The predicted N-binding sites (green) are shown in context with the two PSs (yellow); the fragment shown corresponds to the cartoon in Fig. 5c.

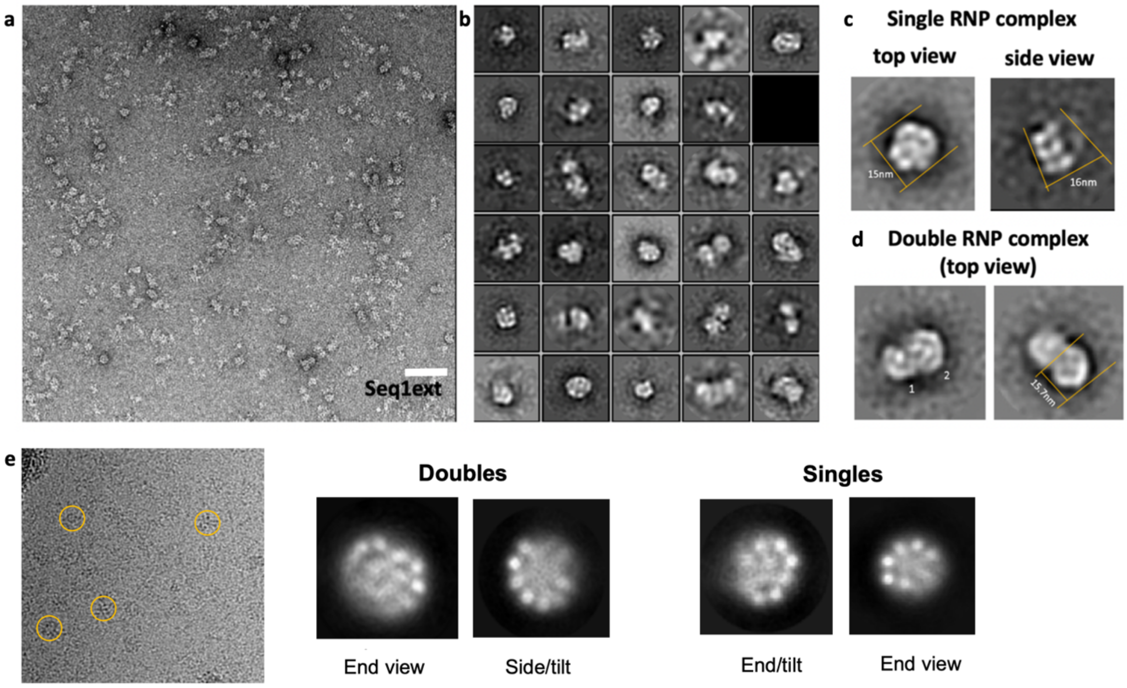


**Fig. S6 | TEM analysis reveals a double RNP complex. a** 2% Uranyl acetate stained SARS-CoV-2 N micrograph of the longer control fragment (Seq1ext), imaged on TEM F20 (120kV) with CCD camera. Size marker = 50 nm. **b** 2D classification (10 images per class) reveals different oligomeric SARS-CoV-2 N states, including a single (**c**) and double (**d**) RNP complex. **e** Images collected on a Krios1 (Falcon4i + Selectris). 2D classification (10 images per class) based on 122,455 particles picked from 3,000 micrographs reveals different oligomeric SARS-CoV-2 N states.

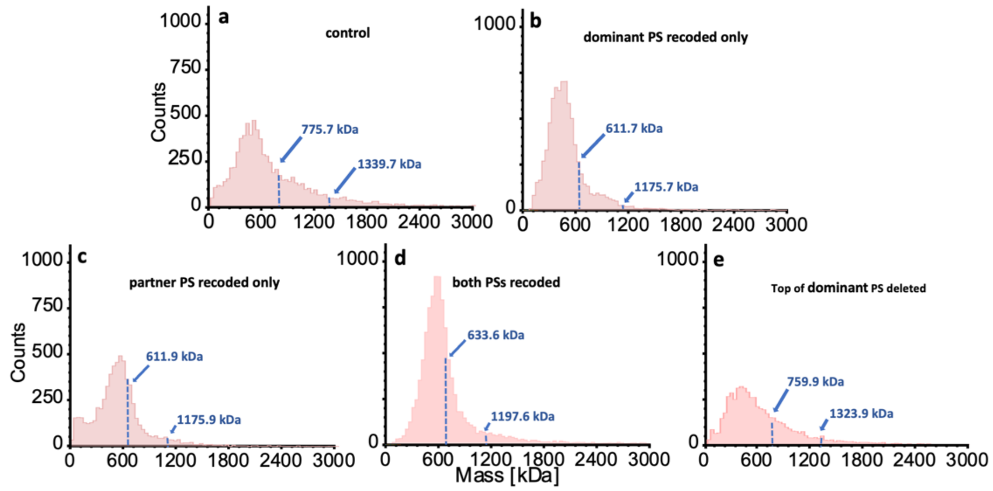


**Fig. S7 | Mass Photometry for different mutants from Mutant Set 2.** The profile of assembly products for the control sequence (**a**) changes if the dominant packaging signal (PS0, **b**) its 5’ partner (PS-1, **c**) or both (**d**) are dyscoded synonymously, or if the top of the packaging signal is deleted (**e**).

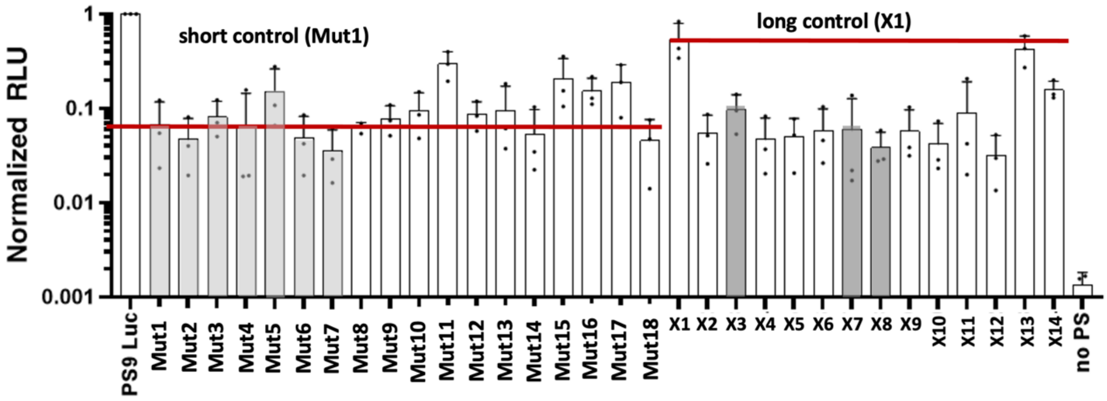


**Fig. S8 | Overview of all experiments performed in the VLP system.** Data also shown in the main text figures are coloured in grey; the corresponding mutant sequences are Mutant Set 2 (Mut1 – Mut18) and Mutant Set 3 (X1 – X14).

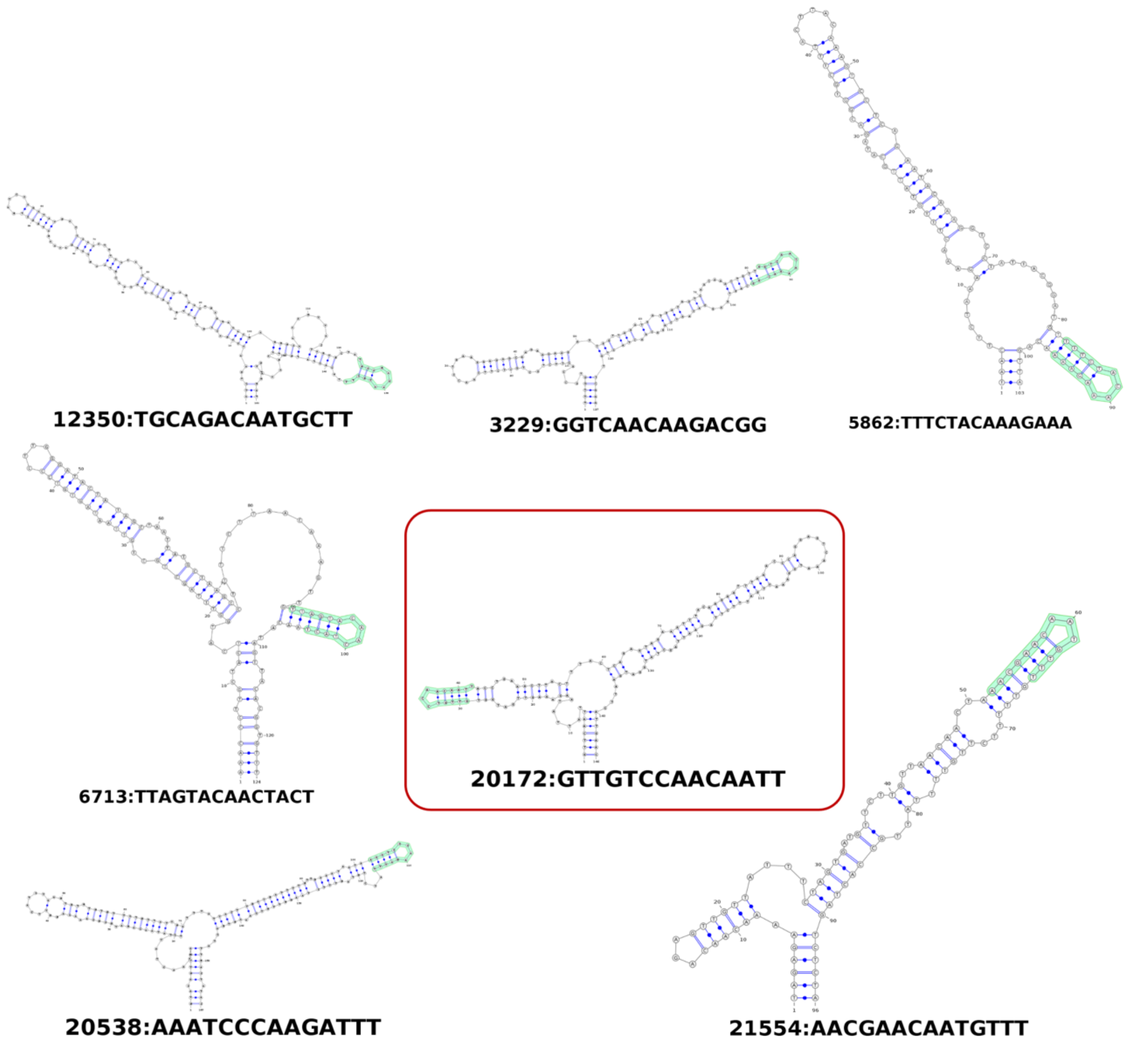


**Fig. S9 | Double stem-loop configurations in the SARS-CoV-2 genome.** There are seven double stem-loop configurations in the SARS-CoV-2 genome presenting a motif from Fig. S3 coincident with the loop portion of one of the stem-loops (green highlights). Interestingly, they are located in areas of the genome known to present higher packaging efficiency according to Fig. 6b (*cf.*, Ref. 15). The element overlapping with a 667-nt fragment implicated in genome packaging is indicated by a red box.

#### Sequences

### Mutations - Set 1

**PS9-m1**

AACAAGCTAGTCTTAATGGAGTCACATTAATTGGAGAAGCCGTAAAAACACAGTTCAATTATTATAAGAAAGTTGATGGT
GTTGTCCAACAATTACCTGAAACTTACTTTACTCAGAGTAGAAATTTACAAGAATTTAAACCCAGGAGTCAAATGGAAATT
GATTTCTTAGAATTAGCTATGGATGAATTCATTGAACGGTATAAATTAGAAGGCTATGCCTTCGAACATATCGTTTATGGA
GATTTTAGTCATTTTCAGTTAGGTTTTTTACATCTACTGATTGGACTAGCTAAACGTTTTAAGGAATCACCTTTTGAATTAGA
AGATTTTATTCCTATGGACAGTACAGTTAAAAACTATTTCATAACAGATGCGCAAACAGGTTCATCTAAGTGTGTGTGTTCT
GTTATTTTTTTATTACTTTTTTTTTTTGTTGAAATAATAAAATCCCAAGATTTATCTGTAGTTTCTAAGGTTGTCAAAGTGACT
ATTGACTATACAGAAATTTCATTTATGCTTTGGTGTAAAGATGGCCATGTAGAAACATTTTACCCAAAATTACAATCTAGTC
AAGCGTGGCAACCGGGTGTTGCTATGCCTAATCTTTACAAAATGCAAAGAATGCTATTAGAAAAGTGTGACCTTCAAAATT
ATGGTGATAGTGCAACATTACCTAAAGGCATAATGATGAATGTCGCAAAATATACTCAACTGTGTCAATATTTAAACACAT
TAACATTAGCTGTACCCTATAATATGAGAGTTATACATTTTTTTGCTGGTTCTTTTAAAGGAGTTGCACCAGGTACAGCTGT
TTTAAGACAGTGGTTGCCTACGGGTACGCTGCTTGTCGATTCAGATCTTTTTGACTTTGTCTCTTTTGCAGATTCAACTTTG
ATTGGTGATTGTGCAACTGTACATACAGCTAATAAATGGGATCTCATTATTAGTGATATGTACGACCCTAAGACTAAAAAT
GTTACAAAAGAAAATGACTCTAAAGAGGGTTTTTTCACTTACATTTGTGGGTTTATACAACAAAAGCTAGCTCTTGGAGGT
TCCGTGGCTATAAAGATAACAGAACATTCTT

**PS9-m2**

AACAAGCTAGTCTTAATGGAGTCACATTAATTGGAGAAGCCGTAAAAACACAGTTCAATTATTATAAGAAAGTTGATGGT
GTTGTCCAACAATTACCTGAAACTTACTTTACTCAGAGTAGAAATTTACAAGAATTTAAACCCAGGAGTCAAATGGAAATT
GATTTCTTAGAATTAGCTATGGATGAATTCATTGAACGGTATAAATTAGAAGGCTATGCCTTCGAACATATCGTTTATGGA
GATTTTAGTCATTTTCAGTTAGGTTTTTTACATCTACTGATTGGACTAGCTAAACGTTTTAAGGAATCACCTTTTGAATTAGA
AGATTTTATTCCTATGGACAGTACAGTTAAAAACTATTTCATAACAGATGCGCAAACAGGTTCATCTAAGTGTGTGTGTTCT
GTTATTTTTTTATTACTTTTTTTTTTTGTTGAAATAATAAAATCCCAAGATTTATCTGTAGTTTCTAAGGTTGTCAAAGTGACT
ATTGACTATACAGAAATTTCATTTATGCTTTGGTGTAAAGATGGCCATGTAGAAACATTTTACCCAAAATTACAATCTAGTC
AAGCGTGGCAACCGGGTGTTGCTATGCCTAATCTTTACAAAATGCAAAGAATGCTATTAGAAAAGTGTGACCTTCAAAATT
ATGGTGATAGTGCAACATTACCTAAAGGCATAATGATGAATGTCGCAAAATATACTCAACTGTGTCAATATTTAAACACAT
TAACATTAGCTGTACCCTATAATATGAGAGTTATACATTTTTTTGCTGGTTCTTTTAAAGGAGTTGCACCAGGTACAGCTGT
TTTAAGACAGTGGTTGCCTACGGGTACGCTGCTTGTCGATTCAGATCTTAATGACTTTGTCTCTGATGCAGATTCAACTTTG
ATTGGTGATTGTGCAACTGTACATACAGCTAATAAATGGGATCTCATTATTAGTGATATGTACGACCCTAAGACTAAAAAT
GTTACAAAAGAAAATGACTCTAAAGAGGGTTTTTTCACTTACATTTGTGGGTTTATACAACAAAAGCTAGCTCTTGGAGGT
TCCGTGGCTATAAAGATAACAGAACATTCTT

**PS9-m3**

AACAAGCTAGTCTTAATGGAGTCACATTAATTGGAGAAGCCGTAAAAACACAGTTCAATTATTATAAGAAAGTTGATGGT
GTTGTCCAACAATTACCTGAAACTTACTTTACTCAGAGTAGAAATTTACAAGAATTTAAACCCAGGAGTCAAATGGAAATT
GATTTCTTAGAATTAGCTATGGATGAATTCATTGAACGGTATAAATTAGAAGGCTATGCCTTCGAACATATCGTTTATGGA
GATTTTAGTCATTTTCAGTTAGGTTTTTTACATCTACTGATTGGACTAGCTAAACGTTTTAAGGAATCACCTTTTGAATTAGA
AGATTTTATTCCTATGGACAGTACAGTTAAAAACTATTTCATAACAGATGCGCAAACAGGTTCATCTAAGTGTGTGTGTTCT
GTTATTGATTTATTACTTGATGATTTTGTTGAAATAATAAAATCCCAAGATTTATCTGTAGTTTCTAAGGTTGTCAAAGTGA
CTATTGACTATACAGAAATTTCATTTATGCTTTGGTGTAAAGATGGCCATGTAGAAACATTTTACCCAAAATTACAATCTAG
TCAAGCGTGGCAACCGGGTGTTGCTATGCCTAATCTTTACAAAATGCAAAGAATGCTATTAGAAAAGTGTGACCTTCAAA
ATTATGGTGATAGTGCAACATTACCTAAAGGCATAATGATGAATGTCGCAAAATATACTCAACTGTGTCAATATTTAAACA
CATTAACATTAGCTGTACCCTATAATATGAGAGTTATACATTTTGGTGCTGGTTCTGATAAAGGAGTTGCACCAGGTACAG
CTGTTTTAAGACAGTGGTTGCCTACGGGTACGCTGCTTGTCGATTCAGATCTTAATGACTTTGTCTCTGATGCAGATTCAAC
TTTGATTGGTGATTGTGCAACTGTACATACAGCTAATAAATGGGATCTCATTATTAGTGATATGTACGACCCTAAGACTAA
AAATGTTACAAAAGAAAATGACTCTAAAGAGGGTTTTTTCACTTACATTTGTGGGTTTATACAACAAAAGCTAGCTCTTGG
AGGTTCCGTGGCTATAAAGATAACAGAACATTCTT

**PS9-m4**
AACAAGCTAGTCTTAATGGAGTCACATTAATTGGAGAAGCCGTAAAAACACAGTTCAATTATTATAAGAAAGTTGATGGT
GTTGTCCAACAATTACCTGAAACTTACTTTACTCAGAGTAGAAATTTACAAGAATTTAAACCCAGGAGTCAAATGGAAATT
GATTTCTTAGAATTAGCTATGGATGAATTCATTGAACGGTATAAATTAGAAGGCTATGCCTTCGAACATATCGTTTATGGA
GATTTTAGTCATAGTCAGTTAGGTGGTTTACATCTACTGATTGGACTAGCTAAACGTTTTAAGGAATCACCTTTTGAATTAG
AAGATTTTATTCCTATGGACAGTACAGTTAAAAACTATTTCATAACAGATGCGCAAACAGGTTCATCTAAGTGTGTGTGTT
CTGTTATTTTTTTATTACTTTTTTTTTTTGTTGAAATAATAAAATCCCAAGATTTATCTGTAGTTTCTAAGGTTGTCAAAGTGA
CTATTGACTATACAGAAATTTCATTTATGCTTTGGTGTAAAGATGGCCATGTAGAAACATTTTACCCAAAATTACAATCTAG
TCAAGCGTGGCAACCGGGTGTTGCTATGCCTAATCTTTACAAAATGCAAAGAATGCTATTAGAAAAGTGTGACCTTCAAA
ATTATGGTGATAGTGCAACATTACCTAAAGGCATAATGATGAATGTCGCAAAATATACTCAACTGTGTCAATATTTAAACA
CATTAACATTAGCTGTACCCTATAATATGAGAGTTATACATTTTGGTGCTGGTTCTGATAAAGGAGTTGCACCAGGTACAG
CTGTTTTAAGACAGTGGTTGCCTACGGGTACGCTGCTTGTCGATTCAGATCTTAATGACTTTGTCTCTGATGCAGATTCAAC
TTTGATTGGTGATTGTGCAACTGTACATACAGCTAATAAATGGGATCTCATTATTAGTGATATGTACGACCCTAAGACTAA
AAATGTTACAAAAGAAAATGACTCTAAAGAGGGTTTTTTCACTTACATTTGTGGGTTTATACAACAAAAGCTAGCTCTTGG
AGGTTCCGTGGCTATAAAGATAACAGAACATTCTT

**PS9-m5**
AACAAGCTAGTCTTAATGGAGTCACATTAATTGGAGAAGCCGTAAAAACACAGTTCAATTATTATAAGAAAGTTGATGGT
GTTGTCCAACAATTACCTGAAACTTACTTTACTCAGAGTAGAAATTTACAAGAATTTAAACCCAGGAGTCAAATGGAAATT
GATTTCTTAGAATTAGCTATGGATGAATTCATTGAACGGTATAAATTAGAAGGCTATGCCTTCGAACATATCGTTTATGGA
GATTTTAGTCATAGTCAGTTAGGTGGTTTACATCTACTGATTGGACTAGCTAAACGTTTTAAGGAATCACCTTTTGAATTAG
AAGATTTTATTCCTATGGACAGTACAGTTAAAAACTATTTCATAACAGATGCGCAAACAGGTTCATCTAAGTGTGTGTGTT
CTGTTATTGATTTATTACTTGATGATTTTGTTGAAATAATAAAATCCCAAGATTTATCTGTAGTTTCTAAGGTTGTCAAAGTG
ACTATTGACTATACAGAAATTTCATTTATGCTTTGGTGTAAAGATGGCCATGTAGAAACATTTTACCCAAAATTACAATCTA
GTCAAGCGTGGCAACCGGGTGTTGCTATGCCTAATCTTTACAAAATGCAAAGAATGCTATTAGAAAAGTGTGACCTTCAA
AATTATGGTGATAGTGCAACATTACCTAAAGGCATAATGATGAATGTCGCAAAATATACTCAACTGTGTCAATATTTAAAC
ACATTAACATTAGCTGTACCCTATAATATGAGAGTTATACATTTTTTTGCTGGTTCTTTTAAAGGAGTTGCACCAGGTACAG
CTGTTTTAAGACAGTGGTTGCCTACGGGTACGCTGCTTGTCGATTCAGATCTTAATGACTTTGTCTCTGATGCAGATTCAAC
TTTGATTGGTGATTGTGCAACTGTACATACAGCTAATAAATGGGATCTCATTATTAGTGATATGTACGACCCTAAGACTAA
AAATGTTACAAAAGAAAATGACTCTAAAGAGGGTTTTTTCACTTACATTTGTGGGTTTATACAACAAAAGCTAGCTCTTGG
AGGTTCCGTGGCTATAAAGATAACAGAACATTCTT

**PS9-m6**
AACAAGCTAGTCTTAATGGAGTCACATTAATTGGAGAAGCCGTAAAAACACAGTTCAATTATTATAAGAAAGTTGATGGT
GTTGTCCAACAATTACCTGAAACTTACTTTACTCAGAGTAGAAATTTACAAGAATTTAAACCCAGGAGTCAAATGGAAATT
GATTTCTTAGAATTAGCTATGGATGAATTCATTGAACGGTATAAATTAGAAGGCTATGCCTTCGAACATATCGTTTATGGA
GATTTTAGTCATTTTCAGTTAGGTTTTTTACATCTACTGATTGGACTAGCTAAACGTTTTAAGGAATCACCTTTTGAATTAGA
AGATTTTATTCCTATGGACAGTACAGTTAAAAACTATTTCATAACAGATGCGCAAACAGGTTCATCTAAGTGTGTGTGTTCT
GTTATTGATTTATTACTTGATGATTTTGTTGAAATAATAAAATCCCAAGATTTATCTGTAGTTTCTAAGGTTGTCAAAGTGA
CTATTGACTATACAGAAATTTCATTTATGCTTTGGTGTAAAGATGGCCATGTAGAAACATTTTACCCAAAATTACAATCTAG
TCAAGCGTGGCAACCGGGTGTTGCTATGCCTAATCTTTACAAAATGCAAAGAATGCTATTAGAAAAGTGTGACCTTCAAA
ATTATGGTGATAGTGCAACATTACCTAAAGGCATAATGATGAATGTCGCAAAATATACTCAACTGTGTCAATATTTAAACA
CATTAACATTAGCTGTACCCTATAATATGAGAGTTATACATTTTTTTGCTGGTTCTTTTAAAGGAGTTGCACCAGGTACAGC
TGTTTTAAGACAGTGGTTGCCTACGGGTACGCTGCTTGTCGATTCAGATCTTTTTGACTTTGTCTCTTTTGCAGATTCAACTT
TGATTGGTGATTGTGCAACTGTACATACAGCTAATAAATGGGATCTCATTATTAGTGATATGTACGACCCTAAGACTAAAA
ATGTTACAAAAGAAAATGACTCTAAAGAGGGTTTTTTCACTTACATTTGTGGGTTTATACAACAAAAGCTAGCTCTTGGAG
GTTCCGTGGCTATAAAGATAACAGAACATTCTT

**PS9-m7**
AACAAGCTAGTCTTAATGGAGTCACATTAATTGGAGAAGCCGTAAAAACACAGTTCAATTATTATAAGAAAGTTGATGGT
GTTGTCCAACAATTACCTGAAACTTACTTTACTCAGAGTAGAAATTTACAAGAATTTAAACCCAGGAGTCAAATGGAAATT
GATTTCTTAGAATTAGCTATGGATGAATTCATTGAACGGTATAAATTAGAAGGCTATGCCTTCGAACATATCGTTTATGGA
GATTTTAGTCATAGTCAGTTAGGTGGTTTACATCTACTGATTGGACTAGCTAAACGTTTTAAGGAATCACCTTTTGAATTAG
AAGATTTTATTCCTATGGACAGTACAGTTAAAAACTATTTCATAACAGATGCGCAAACAGGTTCATCTAAGTGTGTGTGTT
CTGTTATTTTTTTATTACTTTTTTTTTTTGTTGAAATAATAAAATCCCAAGATTTATCTGTAGTTTCTAAGGTTGTCAAAGTGA
CTATTGACTATACAGAAATTTCATTTATGCTTTGGTGTAAAGATGGCCATGTAGAAACATTTTACCCAAAATTACAATCTAG
TCAAGCGTGGCAACCGGGTGTTGCTATGCCTAATCTTTACAAAATGCAAAGAATGCTATTAGAAAAGTGTGACCTTCAAA

ATTATGGTGATAGTGCAACATTACCTAAAGGCATAATGATGAATGTCGCAAAATATACTCAACTGTGTCAATATTTAAACA
CATTAACATTAGCTGTACCCTATAATATGAGAGTTATACATTTTTTTGCTGGTTCTTTTAAAGGAGTTGCACCAGGTACAGC
TGTTTTAAGACAGTGGTTGCCTACGGGTACGCTGCTTGTCGATTCAGATCTTTTTGACTTTGTCTCTTTTGCAGATTCAACTT
TGATTGGTGATTGTGCAACTGTACATACAGCTAATAAATGGGATCTCATTATTAGTGATATGTACGACCCTAAGACTAAAA
ATGTTACAAAAGAAAATGACTCTAAAGAGGGTTTTTTCACTTACATTTGTGGGTTTATACAACAAAAGCTAGCTCTTGGAG
GTTCCGTGGCTATAAAGATAACAGAACATTCTT

**PS9-m8**
AACAAGCTAGTCTTAATGGAGTCACATTAATTGGAGAAGCCGTAAAAACACAGTTCAATTATTATAAGAAAGTTGATGGT
GTTGTCCAACAATTACCTGAAACTTACTTTACTCAGAGTAGAAATTTACAAGAATTTAAACCCAGGAGTCAAATGGAAATT
GATTTCTTAGAATTAGCTATGGATGAATTCATTGAACGGTATAAATTAGAAGGCTATGCCTTCGAACATATCGTTTATGGA
GATTTTAGTCATTTTCAGTTAGGTTTTTTACATCTACTGATTGGACTAGCTAAACGTTTTAAGGAATCACCTTTTGAATTAGA
AGATTTTATTCCTATGGACAGTACAGTTAAAAACTATTTCATAACAGATGCGCAAACAGGTTCATCTAAGTGTGTGTGTTCT
GTTATTTTTTTATTACTTTTTTTTTTTGTTGAAATAATAAAATCCCAAGATTTATCTGTAGTTTCTAAGGTTGTCAAAGTGACT
ATTGACTATACAGAAATTTCATTTATGCTTTGGTGTAAAGATGGCCATGTAGAAACATTTTACCCAAAATTACAATCTAGTC
AAGCGTGGCAACCGGGTGTTGCTATGCCTAATCTTTACAAAATGCAAAGAATGCTATTAGAAAAGTGTGACCTTCAAAATT
ATGGTGATAGTGCAACATTACCTAAAGGCATAATGATGAATGTCGCAAAATATACTCAACTGTGTCAATATTTAAACACAT
TAACATTAGCTGTACCCTATAATATGAGAGTTATACATTTTGGTGCTGGTTCTGATAAAGGAGTTGCACCAGGTACAGCTG
TTTTAAGACAGTGGTTGCCTACGGGTACGCTGCTTGTCGATTCAGATCTTAATGACTTTGTCTCTGATGCAGATTCAACTTT
GATTGGTGATTGTGCAACTGTACATACAGCTAATAAATGGGATCTCATTATTAGTGATATGTACGACCCTAAGACTAAAAA
TGTTACAAAAGAAAATGACTCTAAAGAGGGTTTTTTCACTTACATTTGTGGGTTTATACAACAAAAGCTAGCTCTTGGAGG
TTCCGTGGCTATAAAGATAACAGAACATTCTT

#### Mutations – Set 2

**Mut1**
AAGAAAGTTGATGGTGTTGTCCAACAATTACCTGAAACTTACTTTACTCAGAGTAGAAATTTACAAGAATTTAAACCCAGG
AGTCAAATGGAAATTGATTTCTTAGAATTAGCTATGGATGAATTCATTGAACGGTATAAATTAGAAGGCTATGCCTTCGAA
CATATCGTTTATGGAGATTTTAGTCATAGTCAGTTAGGTGGTTTACATCTACTGATTGGACTAGCTAAACGTTTTAAGGAAT
CACCTTTTGAATTAGAAGATTTTATTCCTATGGACAGTACAGTTAAAAACTATTTCATAACAGATGCGCAAACAGGTTCATC
TAAGTGTGTGTGTTCTGTTATTGATTTATTACTTGATGATTTTGTTGAAATATTTGTTTCTAAGGTTGTCAAAGTGACTATTG
ACTATACAGAAATTTCATTTATGCTTTGGTGTAAAGATGGCCATGTAGAAACATTTTACCCAAAATTACAATCTAGTCAAGC
GTGGCAACCGGGTGTTGCTATGCCTAATCTTTACAAAATGCAAAGAATGCTATTAGAAAAGTGTGACCTTCAAAATTATGG
TGATAGTGCAACATTACCTAAAGGCATAATGATGAATGTCGCAAAATATACTCAACTGTGTCAATATTTAAACA

**Mut2**
AAGAAAGTTGATGGTGTTGTCCAACAATTACCTGAAACTTACTTTACTCAGAGTAGAAATTTACAAGAATTTAAACCCAGG
AGTCAAATGGAAATTGATTTCTTAGAATTAGCTATGGATGAATTCATTGAACGGTATAAATTAGAAGGCTATGCCTTCGAA
CATATCGTTTATGGAGATTTTAGTCATAGTCAGTTAGGTGGTTTACATCTACTGATTGGACTAGCTAAACGTTTTAAGGAAT
CACCTTTTGAATTAGAAGATTTTATTCCTATGGACAGTACAGTTAAAAACTATTTCATAACAGATGCGCAAACAGGTTCATC
TAAGTGTGTGTGTTCTGTTATTGATTTATTACTTTTTAAAGTGACTATTGACTATACAGAAATTTCATTTATGCTTTGGTGTA
AAGATGGCCATGTAGAAACATTTTACCCAAAATTACAATCTAGTCAAGCGTGGCAACCGGGTGTTGCTATGCCTAATCTTT
ACAAAATGCAAAGAATGCTATTAGAAAAGTGTGACCTTCAAAATTATGGTGATAGTGCAACATTACCTAAAGGCATAATG
ATGAATGTCGCAAAATATACTCAACTGTGTCAATATTTAAACA

**Mut3**
AAGAAAGTTGATGGTGTTGTCCAACAATTACCTGAAACTTACTTTACTCAGAGTAGAAATTTACAAGAATTTAAACCCAGG
AGTCAAATGGAAATTGATTTCTTAGAATTAGCTATGGATGAATTCATTGAACGGTATAAATTAGAAGGCTATGCCTTCGAA
CATATCGTTTATGGAGATTTTAGTCATAGTCAGTTAGGTGGTTTACATCTACTGATTGGACTAGCTAAACGTTTTAAGGAAT
CACCTTTTGAATTAGAAGATTTTATTCCTATGGACAGTACAGTTAAAAACTATTTCATAACAGATGCGCAAACAGGTTCATC
TAAGTGTGTGTGTTCTGTTATTGATTTTATTGACTATACAGAAATTTCATTTATGCTTTGGTGTAAAGATGGCCATGTAGAA
ACATTTTACCCAAAATTACAATCTAGTCAAGCGTGGCAACCGGGTGTTGCTATGCCTAATCTTTACAAAATGCAAAGAATG
CTATTAGAAAAGTGTGACCTTCAAAATTATGGTGATAGTGCAACATTACCTAAAGGCATAATGATGAATGTCGCAAAATAT
ACTCAACTGTGTCAATATTTAAACA

**Mut4**
AAGAAAGTTGATGGTGTTGTCCAACAATTACCTGAAACTTACTTTACTCAGAGTAGAAATTTACAAGAATTTAAACCCAGG
AGTCAAATGGAAATTGATTTCTTAGAATTAGCTATGGATGAATTCATTGAACGGTATAAATTAGAAGGCTATGCCTTCGAA
CATATCGTTTATGGAGATTTTAGTCATAGTCAGTTAGGTGGTTTACATCTACTGATTGGACTAGCTAAACGTTTTAAGGAAT
CACCTTTTGAATTAGAAGATTTTATTCCTATGGACAGTACAGTTAAAAACTATTTCATAACAGATGCGCAAACAGGTTCATC
TAAGTGTGTGTGTTCTGTTTTTACAGAAATTTCATTTATGCTTTGGTGTAAAGATGGCCATGTAGAAACATTTTACCCAAAA
TTACAATCTAGTCAAGCGTGGCAACCGGGTGTTGCTATGCCTAATCTTTACAAAATGCAAAGAATGCTATTAGAAAAGTGT
GACCTTCAAAATTATGGTGATAGTGCAACATTACCTAAAGGCATAATGATGAATGTCGCAAAATATACTCAACTGTGTCAA
TATTTAAACA

**Mut5**
AAGAAAGTTGATGGTGTTGTCCAACAATTACCTGAAACTTACTTTACTCAGAGTAGAAATTTACAAGAATTTAAACCCAGG
AGTCAAATGGAAATTGATTTCTTAGAATTAGCTATGGATGAATTCATTGAACGGTATAAATTAGAAGGCTATGCCTTCGAA
CATATCGTTTATGGAGATTTTAGTCATAGTCAGTTAGGTGGTTTACATCTACTGATTGGACTAGCTAAACGTTTTAAGGAAT
CACCTTTTGAATTAGAAGATTTTATTCCTATGGACAGTACAGTTAAAAACTATTTCATAACAGATGCGCAAACAGGTTCATC
TAAGTGTTTTTCATTTATGCTTTGGTGTAAAGATGGCCATGTAGAAACATTTTACCCAAAATTACAATCTAGTCAAGCGTGG
CAACCGGGTGTTGCTATGCCTAATCTTTACAAAATGCAAAGAATGCTATTAGAAAAGTGTGACCTTCAAAATTATGGTGAT
AGTGCAACATTACCTAAAGGCATAATGATGAATGTCGCAAAATATACTCAACTGTGTCAATATTTAAACA

**Mut6**
AAGAAAGTTGATGGTGTTGTCCAACAATTACCTGAAACTTACTTTACTCAGAGTAGAAATTTACAAGAATTTAAACCCAGG
AGTCAAATGGAAATTGATTTCTTAGAATTAGCTATGGATGAATTCATTGAACGGTATAAATTAGAAGGCTATGCCTTCGAA
CATATCGTTTATGGAGATTTTAGTCATAGTCAGTTAGGTGGTTTACATCTACTGATTGGACTAGCTAAACGTTTTAAGGAAT
CACCTTTTGAATTAGAAGATTTTATTCCTATGGACAGTACAGTTAAAAACTATTTCATAACAGATGCGCAAACAGGTTCATC
TATGCTTTGGTGTAAAGATGGCCATGTAGAAACATTTTACCCAAAATTACAATCTAGTCAAGCGTGGCAACCGGGTGTTGC
TATGCCTAATCTTTACAAAATGCAAAGAATGCTATTAGAAAAGTGTGACCTTCAAAATTATGGTGATAGTGCAACATTACC
TAAAGGCATAATGATGAATGTCGCAAAATATACTCAACTGTGTCAATATTTAAACA

**Mut7**
AAGAAAGTTGATGGTGTTGTCCAACAATTACCTGAAACTTACTTTACTCAGAGTAGAAATTTACAAGAATTTAAACCCAGG
AGTCAAATGGAAATTGATTTCTTAGAATTAGCTATGGATGAATTCATTGAACGGTATAAATTAGAAGGCTATGCCTTCGAA
CATATCGTTTATGGAGATTTTAGTCATAGTCAGTTAGGTGGTTTACATCTACTGATTGGACTAGCTAAACGTTTTAAGGAAT
CACCTTTTGAATTAGAAGATTTTATTCCTATGGACAGTACAGTTAAAAACTATTTCATAACAGATGCGCAAACAGGTTCATC
TAAGTGTGTGTGTTCTGTTATTGATCTTCTCCTTGATGATTTTGTTGAAATCATTAAGAGCCAGGATCTTAGTGTTGTCAGC
AAGGTTGTCAAGGTGACGATTGACTATACAGAAATTTCATTTATGCTTTGGTGTAAAGATGGCCATGTAGAAACATTTTAC
CCAAAATTACAATCTAGTCAAGCGTGGCAACCGGGTGTTGCTATGCCTAATCTTTACAAAATGCAAAGAATGCTATTAGAA
AAGTGTGACCTTCAAAATTATGGTGATAGTGCAACATTACCTAAAGGCATAATGATGAATGTCGCAAAATATACTCAACTG
TGTCAATATTTAAACA

**Mut8: Stabilize symmetric fold – synonymous mutations as in Experiment 7, plus three non-synonymous mutations**
AAGAAAGTTGATGGTGTTGTCCAACAATTACCTGAAACTTACTTTACTCAGAGTAGAAATTTACAAGAATTTAAACCCAGG
AGTCAAATGGAAATTGATTTCTTAGAATTAGCTATGGATGAATTCATTGAACGGTATAAATTAGAAGGCTATGCCTTCGAA
CATATCGTTTATGGAGATTTTAGTCATAGTCAGTTAGGTGGTTTACATCTACTGATTGGACTAGCTAAACGTTTTAAGGAAT
CACCTTTTGAATTAGAAGATTTTATTCCTATGGACAGTACAGTTAAAAACTATTTCATAACAGATGCGCAAACAGGTTCATC
TAAGTGTGTGTGTTCTGTTATTGATCTTCTCCTTGATGATTTTGTTGACATCATTAAGAGCCAGGCTCTTAGTGATGTCAGC
AAGGTTGTCAAGGTGACGATTGACTATACAGAAATTTCATTTATGCTTTGGTGTAAAGATGGCCATGTAGAAACATTTTAC
CCAAAATTACAATCTAGTCAAGCGTGGCAACCGGGTGTTGCTATGCCTAATCTTTACAAAATGCAAAGAATGCTATTAGAA
AAGTGTGACCTTCAAAATTATGGTGATAGTGCAACATTACCTAAAGGCATAATGATGAATGTCGCAAAATATACTCAACTG
TGTCAATATTTAAACA

**Mut9: Mutations destabilizing the PS fold**
AAGAAAGTTGATGGTGTTGTCCAACAATTACCTGAAACTTACTTTACTCAGAGTAGAAATTTACAAGAATTTAAACCCAGG
AGTCAAATGGAAATTGATTTCTTAGAATTAGCTATGGATGAATTCATTGAACGGTATAAATTAGAAGGCTATGCCTTCGAA
CATATCGTTTATGGAGATTTTAGTCATAGTCAGTTAGGTGGTTTACATCTACTGATTGGACTAGCTAAACGTTTTAAGGAAT
CACCTTTTGAATTAGAAGATTTTATTCCTATGGACAGTACAGTTAAAAACTATTTCATAACAGATGCGCAAACAGGTTCATC
TAAGTGTGTGTGCAGCGTTATAGACTTACTGTTAGATGATTTTGTGGAAATCATCAAGTCTCAGGATCTTTCCGTCGTCAG

TAAGGTTGTTAAGGTGACTATAGACTATACCGAAATTTCATTTATGCTTTGGTGTAAAGATGGCCATGTAGAAACATTTTA
CCCAAAATTACAATCTAGTCAAGCGTGGCAACCGGGTGTTGCTATGCCTAATCTTTACAAAATGCAAAGAATGCTATTAGA
AAAGTGTGACCTTCAAAATTATGGTGATAGTGCAACATTACCTAAAGGCATAATGATGAATGTCGCAAAATATACTCAACT
GTGTCAATATTTAAACA

**Mut10: 23 nt extension of PS9 at the 5' end**
TAAAAACACAGTTCAATTATTATAAGAAAGTTGATGGTGTTGTCCAACAATTACCTGAAACTTACTTTACTCAGAGTAGAA
ATTTACAAGAATTTAAACCCAGGAGTCAAATGGAAATTGATTTCTTAGAATTAGCTATGGATGAATTCATTGAACGGTATA
AATTAGAAGGCTATGCCTTCGAACATATCGTTTATGGAGATTTTAGTCATAGTCAGTTAGGTGGTTTACATCTACTGATTG
GACTAGCTAAACGTTTTAAGGAATCACCTTTTGAATTAGAAGATTTTATTCCTATGGACAGTACAGTTAAAAACTATTTCAT
AACAGATGCGCAAACAGGTTCATCTAAGTGTGTGTGTTCTGTTATTGATTTATTACTTGATGATTTTGTTGAAATAATAAAA
TCCCAAGATTTATCTGTAGTTTCTAAGGTTGTCAAAGTGACTATTGACTATACAGAAATTTCATTTATGCTTTGGTGTAAAG
ATGGCCATGTAGAAACATTTTACCCAAAATTACAATCTAGTCAAGCGTGGCAACCGGGTGTTGCTATGCCTAATCTTTACA
AAATGCAAAGAATGCTATTAGAAAAGTGTGACCTTCAAAATTATGGTGATAGTGCAACATTACCTAAAGGCATAATGATG
AATGTCGCAAAATATACTCAACTGTGTCAATATTTAAACA

**Mut11: 23 nt extension of PS9 at the 5' end with deletion after the main PS**
TAAAAACACAGTTCAATTATTATAAGAAAGTTGATGGTGTTGTCCAACAATTACCTGAAACTTACTTTACTCAGAGTAGAA
ATTTACAAGAATTTAAACCCAGGAGTCAAATGGAAATTGATTTCTTAGAATTAGCTATGGATGAATTCATTGAACGGTATA
AATTAGAAGGCTATGCCTTCGAACATATCGTTTATGGAGATTTTAGTCATAGTCAGTTAGGTGGTTTACATCTACTGATTG
GACTAGCTAAACGTTTTAAGGAATCACCTTTTGAATTAGAAGATTTTATTCCTATGGACAGTACAGTTAAAAACTATTTCAT
AACAGATGCGCAAACAGGTTCATCTAAGTGTGTGTGTTCTGTTATTGATTTATTACTTGATGATTTTGTTGAAATAATAAAA
TCCCAAGATTTATCTGTAGTTTCTAAGGTTGTCAAAGTGACTATTGACTATACAGAAATTTCATTTATGCTTTGGTGTAAAG
ATGGC

**Mut12: in-frame deletion of the stable secondary structure element**
TAAAAACACAGTTCAATTATTATAAGAAAGTTGATGGTGTTGTCCAACAATTACCTGAAACTTACTTTACTCAGAGTAGAA
ATTTACAAGAATTTAAACCCAGGAGTCAAATGGAAATTGATTTCTTAGAATTAGCTATGGATGAATTCATTGAACGGTATA
AATTAGAAGGCTATGCCTTCGAACATGCGCAAACAGGTTCATCTAAGTGTGTGTGTTCTGTTATTGATTTATTACTTGATGA
TTTTGTTGAAATAATAAAATCCCAAGATTTATCTGTAGTTTCTAAGGTTGTCAAAGTGACTATTGACTATACAGAAATTTCA
TTTATGCTTTGGTGTAAAGATGGC

#### Mutations – Set3

**X1: Long control**
TTATTTAGAAATGCCCGTAATGGTGTTCTTATTACAGAAGGTAGTGTTAAAGGTTTACAACCATCTGTAGGTCCCAAACAA
GCTAGTCTTAATGGAGTCACATTAATTGGAGAAGCCGTAAAAACACAGTTCAATTATTATAAGAAAGTTGATGGTGTTGTC
CAACAATTACCTGAAACTTACTTTACTCAGAGTAGAAATTTACAAGAATTTAAACCCAGGAGTCAAATGGAAATTGATTTC
TTAGAATTAGCTATGGATGAATTCATTGAACGGTATAAATTAGAAGGCTATGCCTTCGAACATATCGTTTATGGAGATTTT
AGTCATAGTCAGTTAGGTGGTTTACATCTACTGATTGGACTAGCTAAACGTTTTAAGGAATCACCTTTTGAATTAGAAGAT
TTTATTCCTATGGACAGTACAGTTAAAAACTATTTCATAACAGATGCGCAAACAGGTTCATCTAAGTGTGTGTGTTCTGTTA
TTGATTTATTACTTGATGATTTTGTTGAAATAATAAAATCCCAAGATTTATCTGTAGTTTCTAAGGTTGTCAAAGTGACTATT
GACTATACAGAAATTTCATTTATGCTTTGGTGTAAAGATGGCCATGTAGAAACATTTTACCCAAAATTACAATCTAGTCAA
GCGTGGCAACCGGGTGTTGCTATGCCTAATCTTTACAAAATGCAAAGAATGCTATTAGAAAAGTGTGACCTTCAAAATTAT
GGTGATAGTGCAACATTACCTAAAGGCATAATGATGAATGTCGCAAAATATACTCAACTGTGTCAATATTTAAACACATTA
ACATTAGCTGTACCCTATAATATGAGAGTTATACATTTTGGTGCTGGTTCTGATAAAGGAGTTGCACCAGGTACAGCTGTT
TTAAGACAGTGGTTGCCTACGGGTACGCTGCTTGTCGATTCAGATCTTAATGACTTTGTCTCTGATGCAGATTCAACTTTGA
TTGGTGATTGTGCAACTGTACATACAGCTAATAAATGGGATCTCATTATTAGTGATATGTACGACCCTAAGACTAAAAATG
TTACAAAAGAAAATGACTCTAAAGAGGGTTTTTTCACTTACATTTGTGGGTTTATACAACAAAAGCTAGCTCTTGGAGGTT
CCGTGGCTATAAAGATA

**X2: Short control**
GTAAAAACACAGTTCAATTATTATAAGAAAGTTGATGGTGTTGTCCAACAATTACCTGAAACTTACTTTACTCAGAGTAGA
AATTTACAAGAATTTAAACCCAGGAGTCAAATGGAAATTGATTTCTTAGAATTAGCTATGGATGAATTCATTGAACGGTAT
AAATTAGAAGGCTATGCCTTCGAACATATCGTTTATGGAGATTTTAGTCATAGTCAGTTAGGTGGTTTACATCTACTGATT

GGACTAGCTAAACGTTTTAAGGAATCACCTTTTGAATTAGAAGATTTTATTCCTATGGACAGTACAGTTAAAAACTATTTCA
TAACAGATGCGCAAACAGGTTCATCTAAGTGTGTGTGTTCTGTTATTGATTTATTACTTGATGATTTTGTTGAAATAATAAA
ATCCCAAGATTTATCTGTAGTTTCTAAGGTTGTCAAAGTGACTATTGACTATACAGAAATTTCATTTATGCTTTGGTGTAAA
GATGGCCATGTAGAAACATTTTACCCAAAATTACAATCTAGTCAAGCGTGGCAACCGGGTGTTGCTATGCCTAATCTTTAC
AAAATGCAAAGAATGCTATTAGAAAAGTGTGACCTTCAAAATTATGGTG

**X3: All PSs/Stops stabilised preventing dynamics**
TTATTTAGAAATGCCCGTAATGGTGTTCTTATTACAGAAGGTAGTGTCAAGGGCCTCCAACCGTCGGTAGGCCCCAAGCA
AGCTAGGCTGAATGGGGTCACACTGATCGGGGAGGCCGTCAAGACACAGTTCAATTATTATAAGAAAGTTGATGGTGTT
GTCCAACAATTACCTGAAACTTACTTTACTCAGAGTAGAAATTTACAAGAATTTAAACCCAGGAGTCAAATGGAGATAGAC
TTCCTAGAATTGGCTATGGATGAATTCATCGAGCGGTATAAACTGGAAGGCTATGCCTTCGAGCATATCGTCTACGGTGA
CTTCAGCCATAGCCAGCTAGGAGGTCTTCATCTACTGATTGGACTAGCTAAACGTTTTAAGGAATCACCTTTTGAATTAGA
AGATTTTATTCCTATGGACAGTACAGTTAAAAACTATTTCATAACAGATGCGCAAACAGGTTCATCTAAGTGTGTATGCTCC
GTGATCGATCTGTTACTTGACGACTTTGTTGAGATCATCAAGTCCCAGGACTTATCGGTGGTCTCCAAAGTCGTCAAAGTA
ACGATCGACTACACGGAGATCTCATTTATGCTTTGGTGTAAAGATGGCCATGTAGAAACATTTTACCCAAAATTACAATCT
AGTCAAGCGTGGCAACCGGGTGTTGCTATGCCTAATCTTTACAAAATGCAAAGAATGCTATTAGAAAAGTGTGACCTTCA
AAATTATGGTGATTCGGCGACGCTGCCGAAGGGCATCATGATGAACGTGGCCAAGTACACACAACTGTGTCAGTACTTGA
ACACGCTCACATTGGCTGTGCCCTACAACATGCGAGTCATCCATTTTGGTGCTGGTTCTGATAAAGGAGTTGCACCAGGTA
CAGCTGTTTTAAGACAGTGGTTGCCTACGGGTACGCTGCTTGTCGATTCAGATCTGAACGACTTTGTCAGCGATGCAGATT
CCACCCTGATCGGCGACTGCGCAACTGTGCACACGGCGAACAAGTGGGATTTGATCATCTCTGACATGTACGACCCCAAG
ACTAAAAATGTTACAAAAGAAAATGACTCTAAAGAGGGTTTTTCACTTACATTTGTGGGTTTATACAACAAAAGCTAGCT
CTTGGAGGTTCCGTGGCTATAAAGATA

**X4: Short – PS0 & PS-1 stabilised**
GTAAAAACACAGTTCAATTATTATAAGAAAGTTGATGGTGTTGTCCAACAATTACCTGAAACTTACTTTACTCAGAGTAGA
AATTTACAAGAATTTAAACCCAGGAGTCAAATGGAGATAGACTTCCTAGAATTGGCTATGGATGAATTCATCGAGCGGTA
TAAACTGGAAGGCTATGCCTTCGAGCATATCGTCTACGGTGACTTCAGCCATAGCCAGCTAGGAGGTCTTCATCTACTGAT
TGGACTAGCTAAACGTTTTAAGGAATCACCTTTTGAATTAGAAGATTTTATTCCTATGGACAGTACAGTTAAAAACTATTTC
ATAACAGATGCGCAAACAGGTTCATCTAAGTGTGTATGCTCCGTGATCGATCTGTTACTTGACGACTTTGTTGAGATCATC
AAGTCCCAGGACTTATCGGTGGTCTCCAAAGTCGTCAAAGTAACGATCGACTACACGGAGATCTCATTTATGCTTTGGTGT
AAAGATGGCCATGTAGAAACATTTTACCCAAAATTACAATCTAGTCAAGCGTGGCAACCGGGTGTTGCTATGCCTAATCTT
TACAAAATGCAAAGAATGCTATTAGAAAAGTGTGACCTTCAAAATTATGGTG

**X5: Level 1 dynamics mutated (PS0 stage 1 only)**
GTAAAAACACAGTTCAATTATTATAAGAAAGTTGATGGTGTTGTCCAACAATTACCTGAAACTTACTTTACTCAGAGTAGA
AATTTACAAGAATTTAAACCCAGGAGTCAAATGGAAATTGATTTCTTAGAATTAGCTATGGATGAATTCATTGAACGGTAT
AAATTAGAAGGCTATGCCTTCGAACATATCGTTTATGGAGATTTTAGTCATAGTCAGTTAGGTGGTTTACATCTACTGATT
GGACTAGCTAAACGTTTTAAGGAGTCGCCGTTCGAGCTGGAGGACTTCATCCCGATGGACAGCACGGTCAAGAACTACTT
CATCACGGACGCGCAAACCGGGTCCTCCAAGTGCGTGTGCTCGGTCATCGACTTATTACTTGATGATTTTGTTGAAATAAT
AAAATCCCAAGATTTATCTGTAGTTTCTAAGGTTGTCAAAGTGACTATTGACTATACAGAAATTTCATTTATGCTTTGGTGT
AAAGATGGCCATGTAGAAACATTTTACCCAAAATTACAATCTAGTCAAGCGTGGCAACCGGGTGTTGCTATGCCTAATCTT
TACAAAATGCAAAGAATGCTATTAGAAAAGTGTGACCTTCAAAATTATGGTG

**X6: Level 2 dynamics mutated (PS-1 stage 1 & PS0 stage 2)**
GTAAAAACACAGTTCAATTATTATAAGAAAGTTGATGGTGTTGTCCAACAATTACCTGAAACTTACTTTACTCAGAGTAGA
AATTTACAAGAATTTAAACCCAGGAGTCAAATGGAAATTGATTTCTTAGAATTAGCTATGGATGAATTCATTGAACGGTAT
AAATTAGAAGGCTATGCCTTCGAACATATCGTTTATGGAGACTTCAGCCATTCGCAGCTGGGCGGCCTGCACCTGCTGATC
GGGCTGGCCAAGCGCTTCAAGGAGAGCCCCTTCGAGCTGGAAGCTTTATTCCTATGGACAGTACCGTCAAGAACTACTTC
ATCACGGACGCGCAGACGGGCTCGTCCAAGTGCGTGTGCTCCGTCATCGACCTGCTGCTGGACGACTTCGTGGAGATCAT
CAAGTCCCAGGACCTGTCGGTGGTCTCCAAGGTCGTCAAGGTGACCATCGACTACACAGAAATTTCATTTATGCTTTGGTG
TAAAGATGGCCATGTAGAAACATTTTACCCAAAATTACAATCTAGTCAAGCGTGGCAACCGGGTGTTGCTATGCCTAATCT
TTACAAAATGCAAAGAATGCTATTAGAAAAGTGTGACCTTCAAAATTATGGTG

**X7: Level 3 dynamics mutated (PS-1 stage 2 & PS0 stage 2)**
GTAAAAACACAGTTCAATTATTATAAGAAAGTTGATGGTGTTGTCCAACAATTACCTGAAACTTACTTTACTCAGAGTAGA
AATTTACAAGAATTTAAACCCAGGAGTCAGATGGAGATCGACCTCCTGGAGCTGGCCATGGACGAGTTCATCGAGCGGTA

CAAGCTGGAGGGCTACGCCTTCGAGCACATCGTCTACGGGGACTTCAGCCACAGCCAGAGCGGCGGCCTGCACCTGCTG
ATCGGGCTGGCCAAGCGCTTCAAGGAGTCCCCCTTCGAATTAGAAGATTTTATTCCTATGGACAGTACCGTCAAGAACTAC
TTCATCACGGACGCGCAGACGGGCTCGTCCAAGTGCGTGTGCTCCGTCATCGACCTGCTGCTGGACGACTTCGTGGAGAT
CATCAAGTCCCAGGACCTGTCGGTGGTCTCCAAGGTCGTCAAGGTGACCATCGACTACACAGAAATTTCATTTATGCTTTG
GTGTAAAGATGGCCATGTAGAAACATTTTACCCAAAATTACAATCTAGTCAAGCGTGGCAACCGGGTGTTGCTATGCCTA
ATCTTTACAAAATGCAAAGAATGCTATTAGAAAAGTGTGACCTTCAAAATTATGGTG

**X8: N protein binding sites surrounding PSs mutated**
GTCAAGACCCAGTTCAACTACTACAAGAAGGTCGACGGGGTCGTCCAGCAGCTGCCCGAGACCTACTTCACCCAGAGCCG
GAACCTGCAGGAGTTCAAGCCCCGGAGCCAGATGGAAATCGATTTCTTAGAATTAGCTATGGATGAATTCATTGAACGGT
ATAAATTAGAAGGCTATGCCTTCGAACATATCGTTTATGGAGATTTTAGTCATAGTCAGTTAGGTGGTTTACATCTACTGAT
TGGACTAGCTAAACGTTTTAAGGAATCACCTTTTGAATTAGAAGATTTTATTCCTATGGACAGTACAGTTAAAAACTATTTC
ATAACAGATGCGCAAACAGGTTCATCTAAGTGTGTGTGTTCTGTTATTGATTTATTACTTGATGATTTTGTTGAAATAATAA
AATCCCAAGATTTATCTGTAGTTTCTAAGGTTGTCAAAGTGACTATTGACTATACAGAAATTTCATTTATGCTTTGGTGTAA
AGATGGCCATGTAGAAACATTTTACCCGAAGCTGCAGTCCTCGCAGGCGTGGCAGCCGGGTGTTGCTATGCCGAACCTGT
ACAACATGCAGCGGATGCTCCTGGAGAAGTGCGACCTCCAGAACTACGGCG

**X9: All N-binding motifs mutated**
GT**G**AA**G**AC**C**CAGTTCAA**C**TA**C**TA**C**AAGAA**G**GTTGATGGTGTTGTCCA**G**CA**GC**T**C**CC**G**GA**G**AC**C**TACTTTACTCAGAGT**C**G
**C**AA**CC**T**C**CA**G**GA**G**TT**C**AA**G**CCCAGGAGTCAAATGGA**G**AT**C**GA**C**TTC**C**T**C**GA**GC**T**C**GCTATGGATGA**G**TTCAT**C**GA**G**CGGT
A**C**AA**CC**T**C**GAAGGCTATGCCTTCGAACA**C**ATCGT**G**TA**C**GG**C**GA**C**TT**C**AGTCATAGTCAGTTAGGTGGTTTACATCTACTGA
TTGGACTAGCTAAACGTTTTAAGGAATCACC**G**TT**C**GA**GC**T**C**GA**G**GA**C**TT**C**AT**C**CCTATGGACAGTACAGT**G**AA**G**AACTA**C**T
TCAT**C**AC**C**GATGCGCAAACAGGTTCATCTAAGTGTGTGTGTTCTGT**G**AT**C**GA**CC**T**CC**T**C**CT**G**GA**C**GA**C**TT**C**GT**G**GA**G**AT**C**A
T**C**AA**G**TCCCAAGATTTATCTGTAGTTTCTAAGGTTGTCAAAGTGACTATTGACTA**C**AC**C**GA**G**AT**C**TC**C**TT**C**ATGCTTTGGTG
TAAAGATGGCCATGTAGA**G**AC**C**TT**C**TACCC**C**AA**GC**T**C**CAATCTAGTCAAGCGTGGCAACCGGGTGTTGCTATGCCTAATCT
TTACAA**G**ATGCA**GC**G**C**ATGCT**CC**T**C**GA**G**AAGTG**C**GACCT**G**CA**G**AA**C**TA**C**GGTG

**X10: In-frame removal of starred SL in PS0 stage 2**
GTAAAAACACAGTTCAATTATTATAAGAAAGTTGATGGTGTTGTCCAACAATTACCTGAAACTTACTTTACTCAGAGTAGA
AATTTACAAGAATTTAAACCCAGGAGTCAAATGGAAATTGATTTCTTAGAATTAGCTATGGATGAATTCATTGAACGGTAT
AAATTAGAAGGCTATGCCTTCGAACATATCGTTTATGGAGATTTTAGTCATAGTCAGTTAGGTGGTTTACATCTACTGATT
GGACTAGCTAAACGTTTTAAGGAATCACCTTTTGAATTAGAAGATTTTATTCCTATGGACAGTACAGTTAAAAACTATTTAT
TACTTGATGATTTTGTTGAAATAATAAAATCCCAAGATTTATCTGTAGTTTCTAAGGTTGTCAAAGTGACTATTGACTATAC
AGAAATTTCATTTATGCTTTGGTGTAAAGATGGCCATGTAGAAACATTTTACCCAAAATTACAATCTAGTCAAGCGTGGCA
ACCGGGTGTTGCTATGCCTAATCTTTACAAAATGCAAAGAATGCTATTAGAAAAGTGTGACCTTCAAAATTATGGTG

**X11: In-frame removal of starred SL in PS-1 stage 2**
GTAAAAACACAGTTCAATTATTATAAGAAAGTTGATGGTGTTGTCCAACAATTACCTGAAACTTACTTTACTCAGAGTAGA
AATTTACAAGAATTTAAACCCAGGAGTCAAATGGAAATTGATTTCTTAGAATTAGCTATGGATGAATTCATTGAACGGTAT
AAATTAGAAGGCTATGCCTTCGAACATATCGTTTATGGAGATTTTGCTAAACGTTTTAAGGAATCACCTTTTGAATTAGAA
GATTTTATTCCTATGGACAGTACAGTTAAAAACTATTTCATAACAGATGCGCAAACAGGTTCATCTAAGTGTGTGTGTTCT
GTTATTGATTTATTACTTGATGATTTTGTTGAAATAATAAAATCCCAAGATTTATCTGTAGTTTCTAAGGTTGTCAAAGTGA
CTATTGACTATACAGAAATTTCATTTATGCTTTGGTGTAAAGATGGCCATGTAGAAACATTTTACCCAAAATTACAATCTAG
TCAAGCGTGGCAACCGGGTGTTGCTATGCCTAATCTTTACAAAATGCAAAGAATGCTATTAGAAAAGTGTGACCTTCAAA
ATTATGGTG

**X12: In-frame removal of starred SLs in both PS0 and PS-1 stage 2**
GTAAAAACACAGTTCAATTATTATAAGAAAGTTGATGGTGTTGTCCAACAATTACCTGAAACTTACTTTACTCAGAGTAGA
AATTTACAAGAATTTAAACCCAGGAGTCAAATGGAAATTGATTTCTTAGAATTAGCTATGGATGAATTCATTGAACGGTAT
AAATTAGAAGGCTATGCCTTCGAACATATCGTTTATGGAGATTTTGCTAAACGTTTTAAGGAATCACCTTTTGAATTAGAA
GATTTTATTCCTATGGACAGTACAGTTAAAAACTATTTATTACTTGATGATTTTGTTGAAATAATAAAATCCCAAGATTTATC
TGTAGTTTCTAAGGTTGTCAAAGTGACTATTGACTATACAGAAATTTCATTTATGCTTTGGTGTAAAGATGGCCATGTAGA
AACATTTTACCCAAAATTACAATCTAGTCAAGCGTGGCAACCGGGTGTTGCTATGCCTAATCTTTACAAAATGCAAAGAAT
GCTATTAGAAAAGTGTGACCTTCAAAATTATGGTG

**X13: PS9 3′ extended to include all 6 N-binding motifs**
TCTGTAGGTCCCAAACAAGCTAGTCTTAATGGAGTCACATTAATTGGAGAAGCCGTAAAAACACAGTTCAATTATTATAAG
AAAGTTGATGGTGTTGTCCAACAATTACCTGAAACTTACTTTACTCAGAGTAGAAATTTACAAGAATTTAAACCCAGGAGT
CAAATGGAAATTGATTTCTTAGAATTAGCTATGGATGAATTCATTGAACGGTATAAATTAGAAGGCTATGCCTTCGAACAT
ATCGTTTATGGAGATTTTAGTCATAGTCAGTTAGGTGGTTTACATCTACTGATTGGACTAGCTAAACGTTTTAAGGAATCA
CCTTTTGAATTAGAAGATTTTATTCCTATGGACAGTACAGTTAAAAACTATTTCATAACAGATGCGCAAACAGGTTCATCTA
AGTGTGTGTGTTCTGTTATTGATTTATTACTTGATGATTTTGTTGAAATAATAAAATCCCAAGATTTATCTGTAGTTTCTAAG
GTTGTCAAAGTGACTATTGACTATACAGAAATTTCATTTATGCTTTGGTGTAAAGATGGCCATGTAGAAACATTTTACCCAA
AATTACAATCTAGTCAAGCGTGGCAACCGGGTGTTGCTATGCCTAATCTTTACAAAATGCAAAGAATGCTATTAGAAAAGT
GTGACCTTCAAAATTATGGTGATAGTGCAACATTACCTAAAGGCATAATGATGAATGTCGCAAAATATACTCAACTGTGTC
AATATTTAAACACATTAACATTAGCTGTACCCTATAATATGAGAGTTATACATTTTGGTGCTGGTTCTGATAAAGGAGTTGC
ACCAGGTACAGCTGTTTTAAGACAGTGGTTGCCTACGGGTACGCTGCTTGTCGATTCAGATCTTAATGACTTTGTCTCTGA
TGCAGATTCAACTTTGATTGGTGATTGTGCAACTGTACATACAGCTAATAAATGGGATCTCATTATTAGTGATATGTACGA
CCCTAAGACTAAAAATGTTACAAAAGAAAATGACTCTAAAGAGGGTTTTTTCACTTACATTTGTGGGTTTATACAACAAAA
GCTAGCTCTTGGAGGTTCCGTGGCTATAAAGATAACAGAACATTCTTGGAATGCTGATCTTTATAAGCTCATGGGACACTT
CGCATGGTGGACAGCCTTTGTTACTAATGTGAATGCGTCATCATCTGAAGCATTTTTAATTGGATGTAATTATCTTGGCAA
A

**X14: PS9 3′ extended to include all 6 N-binding motifs and 5′ reduced to avoid spurious contacts**
GTAAAAACACAGTTCAATTATTATAAGAAAGTTGATGGTGTTGTCCAACAATTACCTGAAACTTACTTTACTCAGAGTAGA
AATTTACAAGAATTTAAACCCAGGAGTCAAATGGAAATTGATTTCTTAGAATTAGCTATGGATGAATTCATTGAACGGTAT
AAATTAGAAGGCTATGCCTTCGAACATATCGTTTATGGAGATTTTAGTCATAGTCAGTTAGGTGGTTTACATCTACTGATT
GGACTAGCTAAACGTTTTAAGGAATCACCTTTTGAATTAGAAGATTTTATTCCTATGGACAGTACAGTTAAAAACTATTTCA
TAACAGATGCGCAAACAGGTTCATCTAAGTGTGTGTGTTCTGTTATTGATTTATTACTTGATGATTTTGTTGAAATAATAAA
ATCCCAAGATTTATCTGTAGTTTCTAAGGTTGTCAAAGTGACTATTGACTATACAGAAATTTCATTTATGCTTTGGTGTAAA
GATGGCCATGTAGAAACATTTTACCCAAAATTACAATCTAGTCAAGCGTGGCAACCGGGTGTTGCTATGCCTAATCTTTAC
AAAATGCAAAGAATGCTATTAGAAAAGTGTGACCTTCAAAATTATGGTGATAGTGCAACATTACCTAAAGGCATAATGAT
GAATGTCGCAAAATATACTCAACTGTGTCAATATTTAAACACATTAACATTAGCTGTACCCTATAATATGAGAGTTATACAT
TTTGGTGCTGGTTCTGATAAAGGAGTTGCACCAGGTACAGCTGTTTTAAGACAGTGGTTGCCTACGGGTACGCTGCTTGTC
GATTCAGATCTTAATGACTTTGTCTCTGATGCAGATTCAACTTTGATTGGTGATTGTGCAACTGTACATACAGCTAATAAAT
GGGATCTCATTATTAGTGATATGTACGACCCTAAGACTAAAAATGTTACAAAAGAAAATGACTCTAAAGAGGGTTTTTTCA
CTTACATTTGTGGGTTTATACAACAAAAGCTAGCTCTTGGAGGTTCCGTGGCTATAAAGATAACAGAACATTCTTGGAATG
CTGATCTTTATAAGCTCATGGGACACTTCGCATGGTGGACAGCCTTTGTTACTAATGTGAATGCGTCATCATCTGAAGCAT
TTTTAATTGGATGTAATTATCTTGGCAAA

#### Mutations – Set4

**F3 dyscoded**
GAGAAACAATGAGTTACTTGTTTCAACATGCCAATTTAGATTCTTGCAAAAGAGTCTTGAACGTGGTGTGTAAAACTTGTG
GACAACAGCAGACAACCCTTAAGGGTGTAGAAGCTGTTATGTACATGGGCACACTTTCTTATGAACAATTTAAGAAAGGT
GTTCAGATACCTTGTACGTGTGGTAAACAAGCTACAAAATATCTAGTACAACAGGAGTCACCTTTTGTTATGATGTCAGCA
CCACCTGCTCAGTATGAACTTAAGCATGGTACATTTACTTGTGCTAGTGAGTACACTGGTAATTACCAGTGTGGTCACTAT
AAACATATAACTTCTAAAGAAACTTTGTATTGCATAGACGGTGCTTTACTTACAAAGTCCTCAGAATACAAAGGTCCTATTA
CGGATGTTTTCTACAAAGAAAACAGTTACACAACAACCATAAAACCAGTTACTTATAAATTGGATGGTGTTGTTTGTACAG
AAATTGACCCTAAGTTGGACAATTATTATAAGAAAGACAATTCTTATTTCACAGAGCAACCAATTGATCTTGTACCAAACC
AACCATATCCAAACGCAAGCTTCGATAATTTTAAGTTTGTATGTGATAATATCAAATTTGCGGATGATCTGAACCAGCTGA
CGGGGTATAAGAAACCGGCGAGCAGAGAGCTGAAAGTGACATTTTTCCCGGACCTGAATGGGGATGTGGTGGCGATTG
ATTATAAACACTACACACCCAGCTTTAAGAAAGGAGCGAAACTGCTGCATAAACCGATTGTGTGGCATGTGAACAATGCA
ACGAATAAAGCCACGTATAAACCAAATACCTGGTGTATACGGTGTCTGTGGAGCACAAAACCAGTGGAAACAAGCAATA
GCTTTGATGTACTGAAGAGCGAGGACGCGCAGGGAATGGATAATCTGGCCTGCGAAGATCTGAAACCAGTCAGCGAAGA
AGTAGTGGAAAATCCGACCATACAGAAAGACGTGCTGGAGTGTAATGTGAAAACGACCGAAGTGGTAGGAGACATTATA
CTGAAACCAGCAAATAATAGCCTGAAAATTACAGAAGAGGTGGGCCACACAGATCTGATGGCGGCGTATGTAGACAATA
GCAGCCTGACGATTAAGAAACCGAATGAACTGAGCAGAGTACTGGGGCTGAAAACCCTGGCGACGCATGGGCTGGCGG
CGGTGAATAGCGTCCCGTGGGATACGATAGCGAATTATGCGAAGCCGTTTCTGAACAAAGTGGTGAGCACAACGACGAA
CATAGTGACACGGTGTCTGAACCGGGTGTGTACGAATTATATGCCGTATTTCTTTACGCTGCTGCTGCAACTGTGTACGTT
TACGAGAAGCACAAATAGCAGAATTAAAGCAAGCATGCCGACGACGATAGCAAAGAATACGGTGAAGAGCGTCGGGAA

ATTTTGTCTGGAGGCGAGCTTTAATTATCTGAAGAGCCCGAATTTTAGCAAACTGATAAATATTATAATTTGGTTTCTGCTG
CTGAGCGTGTGCCTGGGGAGCCTGATCTACAGCACCGCGGCGCTGGGGGTGCTGATGAGCAATCTGGGCATGCCGAGCT
ACTGTACGGGGTACAGAGAAGGCTATCTGAACAGCACGAATGTCACGATTGCAACCTACTGTACGGGGAGCATACCGTG
TAGCGTGTGTCTGAGCGGGCTGGATAGCCTGGACACCTATCCGAGCCTGGAAACGATACAAATTACCATTAGCAGCTTTA
AATGGGATCTGACGGCGTTTGGCCTGGTGGCAGAGTGGTTTCTGGCATATATTCTGTTCACGAGGTTTTTCTATGTACTGG
GACTGGCGGCAATCATGCAACTGTTTTTCAGCTATTTTGCAGTACATTTTATTAGCAATAGCTGGCTGATGTGGCTGATAA
TTAATCTGGTACAAATGGCCCCGATTAGCGCGATGGTGAGAATGTACATCTTCTTTGCAAGCTTTTATTATGTATGGAAAA
GCTATGTGCATGTGGTAGACGGGTGTAATAGCAGCACGTGTATGATGTGTTACAAACGGAATAGAGCAACAAGAGTCGA
ATGTACAACGATTGTGAATGGGGTGAGAAGGAGCTTTTATGTCTATGCGAATGGAGGGAAAGGCTTTTGCAAACTGCAC
AATTGGAATTGTGTGAATTGTGATACATTCTGTGCGGGGAGCACATTTATTAGCGATGAAGTGGCGAGAGACgTGAGCCT
GCAGTTTAAAAGACCAATAAATCCGACGGACCAGAGCAGCTACATCGTGGATAGCGTGACAGTGAAGAATGGGAGCATC
CATCTGTACTTTGATAAAGCGGGGCAAAAGACGTATGAAAGACATAGCCTGAGCCATTTTGTGAACCTGGACAACCTGAG
AGCGAATAACACGAAAGGGAGCCTGCCGATTAATGTGATAGTGTTTGATGGGAAAAGCAAATGTGAAGAAAGCAGCGC
AAAAAGCGCGAGCGTGTACTACAGCCAGCTGATGTGTCAACCGATACTGCTGCTGGATCAGGCACTGGTGAGCGATGTG
GGGGATAGCGCGGAAGTGGCAGTGAAAATGTTTGATGCGTACGTGAATACGTTTAGCAGCACGTTTAACGTACCAATGG
AAAAACTGAAAACACTGGTGGCAACGGCAGAAGCGGAACTGGCAAAGAATGTGAGCCTGGACAATGTCCTGAGCACGTT
TATTAGCGCAGCGCGGCAAGGGTTTGTGGATAGCGATGTAGAAACGAAAGATGTGGTGGAATGTCTGAAACTGAGCCAT
CAAAGCGACATAGAAGTGACGGGCGATAGCTGTAATAACTATATGCTGACCTATAACAAAGTGGAAAACATGACACCCC
GGGACCTGGGGGCGTGTATTGACTGTAGCGCGCGGCATATTAATGCGCAGGTAGCAAAAAGCCACAACATTGCGCTGAT
ATGGAACGTGAAAGATTTCATGAGCCTGAGCGAACAACTGCGAAAACAAATACGGAGCGCGGCGAAAAAGAATAACCTG
CCGTTTAAGCTGACATGTGCAACGACGAGACAAGTGGTGAATGTGGTAACAACAAAGATAGCACTGAAGGGGGGTAA

**F3 encoded**

GAGAAACAATGAGTTACTTGTTTCAACATGCCAATTTAGATTCTTGCAAAAGAGTCTTGAACGTGGTGTGTAAAACTTGTG
GACAACAGCAGACAACCCTTAAGGGTGTAGAAGCTGTTATGTACATGGGCACACTTTCTTATGAACAATTTAAGAAAGGT
GTTCAGATACCTTGTACGTGTGGTAAACAAGCTACAAAATATCTAGTACAACAGGAGTCACCTTTTGTTATGATGTCAGCA
CCACCTGCTCAGTATGAACTTAAGCATGGTACATTTACTTGTGCTAGTGAGTACACTGGTAATTACCAGTGTGGTCACTAT
AAACATATAACTTCTAAAGAAACTTTGTATTGCATAGACGGTGCTTTACTTACAAAGTCCTCAGAATACAAAGGTCCTATTA
CGGATGTTTTCTACAAAGAAAACAGTTACACAACAACCATAAAACCAGTTACTTATAAATTGGATGGTGTTGTTTGTACAG
AAATTGACCCTAAGTTGGACAATTATTATAAGAAAGACAATTCTTATTTCACAGAGCAACCAATTGATCTTGTACCAAACC
AACCTTATCCTAATGCTAGTTTTGATAATTTTAAGTTTGTTTGTGATAATATTAAATTTGCTGATGATTTAAATCAGTTAACT
GGTTATAAGAAACCTGCTAGTCGTGAGTTAAAAGTTACTTTTTTTCCTGATTTAAATGGTGATGTTGTTGCTATTGATTATA
AACATTATACTCCTAGTTTTAAGAAAGGTGCTAAATTATTACATAAACCTATTGTTTGGCATGTTAATAATGCTACTAATAA
AGCTACTTATAAACCTAATACTTGGTGTATTCGTTGTTTATGGAGTACTAAACCTGTTGAAACTAGTAATAGTTTTGATGTT
TTAAAGAGTGAGGATGCTCAGGGTATGGATAATTTAGCTTGTGAAGATTTAAAACCTGTTAGTGAAGAAGTTGTTGAAAA
TCCTACTATTCAGAAAGATGTTTTAGAGTGTAATGTTAAAACTACTGAAGTTGTTGGTGATATTATTTTAAAACCTGCTAAT
AATAGTTTAAAAATTACTGAAGAGGTTGGTCATACTGATTTAATGGCTGCTTATGTTGATAATAGTAGTTTAACTATTAAG
AAACCTAATGAATTAAGTCGTGTTTTAGGTTTAAAAACTTTAGCTACTCATGGTTTAGCTGCTGTTAATAGTGTTCCTTGGG
ATACTATTGCTAATTATGCTAAGCCTTTTTTAAATAAAGTTGTTAGTACTACTACTAATATTGTTACTCGTTGTTTAAATCGT
GTTTGTACTAATTATATGCCTTATTTTTTTACTTTATTATTACAATTATGTACTTTTACTCGTAGTACTAATAGTCGTATTAAA
GCTAGTATGCCTACTACTATTGCTAAGAATACTGTTAAGAGTGTTGGTAAATTTTGTTTAGAGGCTAGTTTTAATTATTTAA
AGAGTCCTAATTTTAGTAAATTAATTAATATTATTATTTGGTTTTTATTATTAAGTGTTTGTTTAGGTAGTTTAATTTATAGTA
CTGCTGCTTTAGGTGTTTTAATGAGTAATTTAGGTATGCCTAGTTATTGTACTGGTTATCGTGAAGGTTATTTAAATAGTAC
TAATGTTACTATTGCTACTTATTGTACTGGTAGTATTCCTTGTAGTGTTTGTTTAAGTGGTTTAGATAGTTTAGATACTTATC
CTAGTTTAGAAACTATTCAAATTACTATTAGTAGTTTTAAATGGGATTTAACTGCTTTTGGTTTAGTTGCTGAGTGGTTTTTA
GCTTATATTTTATTTACTCGTTTTTTTTATGTTTTAGGTTTAGCTGCTATTATGCAATTATTTTTTAGTTATTTTGCTGTTCATT
TTATTAGTAATAGTTGGTTAATGTGGTTAATTATTAATTTAGTTCAAATGGCTCCTATTAGTGCTATGGTTCGTATGTATATT
TTTTTTGCTAGTTTTTATTATGTTTGGAAAAGTTATGTTCATGTTGTTGATGGTTGTAATAGTAGTACTTGTATGATGTGTTA
TAAACGTAATCGTGCTACTCGTGTTGAATGTACTACTATTGTTAATGGTGTTCGTCGTAGTTTTTATGTTTATGCTAATGGT
GGTAAAGGTTTTTGTAAATTACATAATTGGAATTGTGTTAATTGTGATACTTTTTGTGCTGGTAGTACTTTTATTAGTGATG
AAGTTGCTCGTGATTTAAGTTTACAGTTTAAACGTCCTATTAATCCTACTGATCAGAGTAGTTATATTGTTGATAGTGTTAC
TGTTAAGAATGGTAGTATTCATTTATATTTTGATAAAGCTGGTCAAAAGACTTATGAACGTCATAGTTTAAGTCATTTTGTT
AATTTAGATAATTTACGTGCTAATAATACTAAAGGTAGTTTACCTATTAATGTTATTGTTTTTGATGGTAAAAGTAAATGTG
AAGAAAGTAGTGCTAAAAGTGCTAGTGTTTATTATAGTCAGTTAATGTGTCAACCTATTTTATTATTAGATCAGGCTTTAGT
TAGTGATGTTGGTGATAGTGCTGAAGTTGCTGTTAAAATGTTTGATGCTTATGTTAATACTTTTAGTAGTACTTTTAATGTT
CCTATGGAAAAATTAAAAACTTTAGTTGCTACTGCTGAAGCTGAATTAGCTAAGAATGTTAGTTTAGATAATGTTTTAAGT
ACTTTTATTAGTGCTGCTCGTCAAGGTTTTGTTGATAGTGATGTTGAAACTAAAGATGTTGTTGAATGTTTAAAATTAAGTC

ATCAAAGTGATATTGAAGTTACTGGTGATAGTTGTAATAATTATATGTTAACTTATAATAAAGTTGAAAATATGACTCCTCG
TGATTTAGGTGCTTGTATTGATTGTAGTGCTCGTCATATTAATGCTCAGGTTGCTAAAAGTCATAATATTGCTTTAATTTGG
AATGTTAAAGATTTTATGAGTTTAAGTGAACAATTACGTAAACAAATTCGTAGTGCTGCTAAAAAGAATAATTTACCTTTTA
AGTTAACTTGTGCTACTACTCGTCAAGTTGTTAATGTTGTTACTACTAAGATTGCTTTAAAGGGTGGTAA

**F4 dyscoded**
GTAAAATTGTGAATAATTGGCTGAAGCAGCTGATTAAAGTGACACTGGTGTTCCTGTTTGTGGCGGCGATTTTCTATCTGA
TAACACCGGTGCATGTCATGAGCAAACATACGGACTTTAGCAGCGAAATCATAGGATACAAGGCGATTGATGGGGGGGT
CACGCGGGACATAGCAAGCACAGATACGTGTTTTGCGAACAAACATGCGGATTTTGACACATGGTTTAGCCAGCGGGGG
GGGAGCTATACGAATGACAAAGCGTGCCCACTGATTGCGGCAGTCATAACAAGAGAAGTGGGGTTTGTCGTGCCGGGGC
TGCCGGGCACGATACTGCGCACAACGAATGGGGACTTTCTGCATTTCCTGCCGAGAGTGTTTAGCGCAGTGGGGAACATC
TGTTACACACCAAGCAAACTGATAGAGTACACGGACTTTGCAACAAGCGCGTGTGTGCTGGCGGCGGAATGTACAATTTT
TAAAGATGCGAGCGGGAAGCCAGTACCATATTGTTATGATACCAATGTACTGGAAGGGAGCGTGGCGTATGAAAGCCTG
CGCCCGGACACACGGTATGTGCTGATGGATGGCAGCATTATTCAATTTCCGAACACCTACCTGGAAGGGAGCGTGAGAGT
GGTAACAACGTTTGATAGCGAGTACTGTAGGCACGGCACGTGTGAAAGAAGCGAAGCGGGGGTGTGTGTAAGCACGAG
CGGGAGATGGGTACTGAACAATGATTATTACAGAAGCCTGCCAGGAGTGTTCTGTGGGGTAGATGCGGTAAATCTGCTG
ACGAATATGTTTACACCACTGATTCAACCGATTGGGGCGCTGGACATAAGCGCAAGCATAGTAGCGGGGGGGATTGTAG
CGATCGTAGTAACATGCCTGGCCTACTATTTTATGAGGTTTAGAAGAGCGTTTGGGGAATACAGCCATGTAGTGGCCTTTA
ATACGCTGCTGTTCCTGATGAGCTTCACGGTACTGTGTCTGACACCAGTGTACAGCTTCCTGCCGGGGGTGTATAGCGTGA
TTTACCTGTACCTGACATTTTATCTGACGAATGATGTGAGCTTTCTGGCACATATTCAGTGGATGGTGATGTTCACACCGCT
GGTACCGTTCTGGATAACAATTGCGTATATCATTTGTATTAGCACAAAGCATTTCTATTGGTTCTTTAGCAATTACCTGAAG
AGACGGGTAGTCTTTAATGGGGTGAGCTTTAGCACGTTTGAAGAAGCGGCGCTGTGCACCTTTCTGCTGAATAAAGAAAT
GTATCTGAAGCTGCGGAGCGATGTGCTGCTGCCGCTGACGCAATATAATAGATACCTGGCGCTGTATAATAAGTACAAGT
ATTTTAGCGGAGCAATGGATACAACGAGCTACAGAGAAGCGGCGTGTTGTCATCTGGCAAAGGCGCTGAATGACTTCAG
CAACAGCGGGAGCGATGTGCTGTACCAACCACCACAAACCAGCATCACCAGCGCTGTTTTGCAGAGTGGTTTTAGAAAAA
TGGCATTCCCATCTGGTAAAGTTGAGGGTTGTATGGTACAAGTAACTTGTGGTACAACTACACTTAACGGTCTTTGGCTTG
ATGACGTAGTTTACTGTCCAAGACATGTGATCTGCACCTCTGAAGACATGCTTAACCCTAATTATGAAGATTTACTCATTCG
TAAGTCTAATCATAATTTCTTGGTACAGGCTGGTAATGTTCAACTCAGGGTTATTGGACATTCTATGCAAAATTGTGTACTT
AAGCTTAAGGTTGATACAGCCAATCCTAAGACACCTAAGTATAAGTTTGTTCGCATTCAACCAGGACAGACTTTTTCAGTG
TTAGCTTGTTACAATGGTTCACCATCTGGTGTTTACCAATGTGCTATGAGGCCCAATTTCACTATTAAGGGTTCATTCCTTA
ATGGTTCATGTGGTAGTGTTGGTTTTAACATAGATTATGACTGTGTCTCTTTTTGTTACATGCACCATATGGAATTACCAAC
TGGAGTTCATGCTGGCACAGACTTAGAAGGTAACTTTTATGGACCTTTTGTTGACAGGCAAACAGCACAAGCAGCTGGTA
CGGACACAACTATTACAGTTAATGTTTTAGCTTGGTTGTACGCTGCTGTTATAAATGGAGACAGGTGGTTTCTCAATCGAT
TTACCACAACTCTTAATGACTTTAACCTTGTGGCTATGAAGTACAATTATGAACCTCTAACACAAGACCATGTTGACATACT
AGGACCTCTTTCTGCTCAAACTGGAATTGCCGTTTTAGATATGTGTGCTTCATTAAAAGAATTACTGCAAAATGGTATGAA
TGGACGTACCATATTGGGTAGTGCTTTATTAGAAGATGAATTTACACCTTTTGATGTTGTTAGACAATGCTCAGGTGTTACT
TTCCAAAGTGCAGTGAAAAGAACAATCAAGGGTACACACCACTGGTTGTTACTCACAATTTTGACTTCACTTTTAGTTTTAG
TCCAGAGTACTCAATGGTCTTTGTTCTTTTTTTTGTATGAAAATGCCTTTTTACCTTTTGCTATGGGTATTATTGCTATGTCTG
CTTTTGCAATGATGTTTGTCAAACATAAGCATGCATTTCTCTGTTTGTTTTTGTTACCTTCTCTTGCCACTGTAGCTTATTTTA
ATATGGTCTATATGCCTGCTAGTTGGGTGATGCGTATTATGACATGGTTGGATATGGTTGATACTAGTTTGTCTGGTTTTA
AGCTAAAAGACTGTGTTATGTATGCATCAGCTGTAGTGTTACTAATCCTTATGACAGCAAGAACTGTGTATGATGATGGTG
CTAGGAGAGTGTGGACACTTATGAATGTCTTGACACTCGTTTATAAAGTTTATTATGGTAATGCTTTAGATCAAGCCATTTC
CATGTGGGCTCTTATAATCTCTGTTACTTCTAACTACTCAGGTGTAGTTACAACTGTCATGTTTTTGGCCAGAGGTATTGTTT
TTATGTGTGTTGAGTATTGCCCTATTTTCTTCATAACTGGTAATACACTTCAGTGTATAATGCTAGTTTATTGTTTCTTAGGC
TATTTTTGTACTTGTTACTTTGGCCTCTTTTGTTTACTCAACCGCTACTTTAGACTGACTCTTGGTGTTTATGATTACTTAGTT
TCTACACAGGAGTTTAGATATATGAATTCACAGGGACTACTCCCACCCAAGAATAGCATAGATGCCTTCAAACTCAACATT
AAATTGTTGGGTGTTGGTGGCAAACCTTGTATCAAAGTAGCCACTGTACAGTCTA

**F4 encoded**
GTAAAATTGTTAATAATTGGTTAAAGCAGTTAATTAAAGTTACTTTAGTTTTTTTATTTGTTGCTGCTATTTTTTATTTAATTA
CTCCTGTTCATGTTATGAGTAAACATACTGATTTTAGTAGTGAAATTATTGGTTATAAGGCTATTGATGGTGGTGTTACTCG
TGATATTGCTAGTACTGATACTTGTTTTGCTAATAAACATGCTGATTTTGATACTTGGTTTAGTCAGCGTGGTGGTAGTTAT
ACTAATGATAAAGCTTGTCCTTTAATTGCTGCTGTTATTACTCGTGAAGTTGGTTTTGTTGTTCCTGGTTTACCTGGTACTAT
TTTACGTACTACTAATGGTGATTTTTTACATTTTTTACCTCGTGTTTTTAGTGCTGTTGGTAATATTTGTTATACTCCTAGTAA
ATTAATTGAGTATACTGATTTTGCTACTAGTGCTTGTGTTTTAGCTGCTGAATGTACTATTTTTAAAGATGCTAGTGGTAAG
CCTGTTCCTTATTGTTATGATACTAATGTTTTAGAAGGTAGTGTTGCTTATGAAAGTTTACGTCCTGATACTCGTTATGTTTT
AATGGATGGTAGTATTATTCAATTTCCTAATACTTATTTAGAAGGTAGTGTTCGTGTTGTTACTACTTTTGATAGTGAGTAT

TGTCGTCATGGTACTTGTGAACGTAGTGAAGCTGGTGTTTGTGTTAGTACTAGTGGTCGTTGGGTTTTAAATAATGATTAT
TATCGTAGTTTACCTGGTGTTTTTTGTGGTGTTGATGCTGTTAATTTATTAACTAATATGTTTACTCCTTTAATTCAACCTATT
GGTGCTTTAGATATTAGTGCTAGTATTGTTGCTGGTGGTATTGTTGCTATTGTTGTTACTTGTTTAGCTTATTATTTTATGCG
TTTTCGTCGTGCTTTTGGTGAATATAGTCATGTTGTTGCTTTTAATACTTTATTATTTTTAATGAGTTTTACTGTTTTATGTTT
AACTCCTGTTTATAGTTTTTTACCTGGTGTTTATAGTGTTATTTATTTATATTTAACTTTTTATTTAACTAATGATGTTAGTTTT
TTAGCTCATATTCAGTGGATGGTTATGTTTACTCCTTTAGTTCCTTTTTGGATTACTATTGCTTATATTATTTGTATTAGTACT
AAGCATTTTTATTGGTTTTTTAGTAATTATTTAAAGCGTCGTGTTGTTTTTAATGGTGTTAGTTTTAGTACTTTTGAAGAAGC
TGCTTTATGTACTTTTTTATTAAATAAAGAAATGTATTTAAAGTTACGTAGTGATGTTTTATTACCTTTAACTCAATATAATC
GTTATTTAGCTTTATATAATAAGTATAAGTATTTTAGTGGTGCTATGGATACTACTAGTTATCGTGAAGCTGCTTGTTGTCA
TTTAGCTAAGGCTTTAAATGATTTTAGTAATAGTGGTAGTGATGTTTTATATCAACCTCCTCAAACTAGTATTACTAGTGCT
GTTTTGCAGAGTGGTTTTAGAAAAATGGCATTCCCATCTGGTAAAGTTGAGGGTTGTATGGTACAAGTAACTTGTGGTAC
AACTACACTTAACGGTCTTTGGCTTGATGACGTAGTTTACTGTCCAAGACATGTGATCTGCACCTCTGAAGACATGCTTAA
CCCTAATTATGAAGATTTACTCATTCGTAAGTCTAATCATAATTTCTTGGTACAGGCTGGTAATGTTCAACTCAGGGTTATT
GGACATTCTATGCAAAATTGTGTACTTAAGCTTAAGGTTGATACAGCCAATCCTAAGACACCTAAGTATAAGTTTGTTCGC
ATTCAACCAGGACAGACTTTTTCAGTGTTAGCTTGTTACAATGGTTCACCATCTGGTGTTTACCAATGTGCTATGAGGCCCA
ATTTCACTATTAAGGGTTCATTCCTTAATGGTTCATGTGGTAGTGTTGGTTTTAACATAGATTATGACTGTGTCTCTTTTTGT
TACATGCACCATATGGAATTACCAACTGGAGTTCATGCTGGCACAGACTTAGAAGGTAACTTTTATGGACCTTTTGTTGAC
AGGCAAACAGCACAAGCAGCTGGTACGGACACAACTATTACAGTTAATGTTTTAGCTTGGTTGTACGCTGCTGTTATAAAT
GGAGACAGGTGGTTTCTCAATCGATTTACCACAACTCTTAATGACTTTAACCTTGTGGCTATGAAGTACAATTATGAACCTC
TAACACAAGACCATGTTGACATACTAGGACCTCTTTCTGCTCAAACTGGAATTGCCGTTTTAGATATGTGTGCTTCATTAAA
AGAATTACTGCAAAATGGTATGAATGGACGTACCATATTGGGTAGTGCTTTATTAGAAGATGAATTTACACCTTTTGATGT
TGTTAGACAATGCTCAGGTGTTACTTTCCAAAGTGCAGTGAAAAGAACAATCAAGGGTACACACCACTGGTTGTTACTCAC
AATTTTGACTTCACTTTTAGTTTTAGTCCAGAGTACTCAATGGTCTTTGTTCTTTTTTTTGTATGAAAATGCCTTTTTACCTTT
TGCTATGGGTATTATTGCTATGTCTGCTTTTGCAATGATGTTTGTCAAACATAAGCATGCATTTCTCTGTTTGTTTTTGTTAC
CTTCTCTTGCCACTGTAGCTTATTTTAATATGGTCTATATGCCTGCTAGTTGGGTGATGCGTATTATGACATGGTTGGATAT
GGTTGATACTAGTTTGTCTGGTTTTAAGCTAAAAGACTGTGTTATGTATGCATCAGCTGTAGTGTTACTAATCCTTATGACA
GCAAGAACTGTGTATGATGATGGTGCTAGGAGAGTGTGGACACTTATGAATGTCTTGACACTCGTTTATAAAGTTTATTAT
GGTAATGCTTTAGATCAAGCCATTTCCATGTGGGCTCTTATAATCTCTGTTACTTCTAACTACTCAGGTGTAGTTACAACTG
TCATGTTTTTGGCCAGAGGTATTGTTTTTATGTGTGTTGAGTATTGCCCTATTTTCTTCATAACTGGTAATACACTTCAGTGT
ATAATGCTAGTTTATTGTTTCTTAGGCTATTTTTGTACTTGTTACTTTGGCCTCTTTTGTTTACTCAACCGCTACTTTAGACTG
ACTCTTGGTGTTTATGATTACTTAGTTTCTACACAGGAGTTTAGATATATGAATTCACAGGGACTACTCCCACCCAAGAATA
GCATAGATGCCTTCAAACTCAACATTAAATTGTTGGGTGTTGGTGGCAAACCTTGTATCAAAGTAGCCACTGTACAGTCTA

**F7X3**

GCTGAAAATGTAACAGGACTCTTTAAAGATTGTAGTAAGGTAATCACTGGGTTACATCCTACACAGGCACCTACACACCTC
AGTGTTGACACTAAATTCAAAACTGAAGGTTTATGTGTTGACATACCTGGCATACCTAAGGACATGACCTATAGAAGACTC
ATCTCTATGATGGGTTTTAAAATGAATTATCAAGTTAATGGTTACCCTAACATGTTTATCACCCGCGAAGAAGCTATAAGA
CATGTACGTGCATGGATTGGCTTCGATGTCGAGGGGTGTCATGCTACTAGAGAAGCTGTTGGTACCAATTTACCTTTACAG
CTAGGTTTTTCTACAGGTGTTAACCTAGTTGCTGTACCTACAGGTTATGTTGATACACCTAATAATACAGATTTTTCCAGAG
TTAGTGCTAAACCACCGCCTGGAGATCAATTTAAACACCTCATACCACTTATGTACAAAGGACTTCCTTGGAATGTAGTGC
GTATAAAGATTGTACAAATGTTAAGTGACACACTTAAAAATCTCTCTGACAGAGTCGTATTTGTCTTATGGGCACATGGCT
TTGAGTTGACATCTATGAAGTATTTTGTGAAAATAGGACCTGAGCGCACCTGTTGTCTATGTGATAGACGTGCCACATGCT
TTTCCACTGCTTCAGACACTTATGCCTGTTGGCATCATTCTATTGGATTTGATTACGTCTATAATCCGTTTATGATTGATGTT
CAACAATGGGGTTTTACAGGTAACCTACAAAGCAACCATGATCTGTATTGTCAAGTCCATGGTAATGCACATGTAGCTAGT
TGTGATGCAATCATGACTAGGTGTCTAGCTGTCCACGAGTGCTTTGTTAAGCGTGTTGACTGGACTATTGAATATCCTATA
ATTGGTGATGAACTGAAGATTAATGCGGCTTGTAGAAAGGTTCAACACATGGTTGTTAAAGCTGCATTATTAGCAGACAA
ATTCCCAGTTCTTCACGACATTGGTAACCCTAAAGCTATTAAGTGTGTACCTCAAGCTGATGTAGAATGGAAGTTCTATGA
TGCACAGCCTTGTAGTGACAAAGCTTATAAAATAGAAGAATTATTCTATTCTTATGCCACACATTCTGACAAATTCACAGAT
GGTGTATGCCTATTTTGGAATTGCAATGTCGATAGATATCCTGCTAATTCCATTGTTTGTAGATTTGACACTAGAGTGCTAT
CTAACCTTAACTTGCCTGGTTGTGATGGTGGCAGTTTGTATGTAAATAAACATGCATTCCACACACCAGCTTTTGATAAAA
GTGCTTTTGTTAATTTAAAACAATTACCATTTTTCTATTACTCTGACAGTCCATGTGAGTCTCATGGAAAACAAGTAGTGTC
AGATATAGATTATGTACCACTAAAGTCTGCTACGTGTATAACACGTTGCAATTTAGGTGGTGCTGTCTGTAGACATCATGC
TAATGAGTACAGATTGTATCTCGATGCTTATAACATGATGATCTCAGCTGGCTTTAGCTTGTGGGTTTACAAACAATTTGAT
ACTTATAACCTCTGGAACACTTTTACAAGACTTCAGAGTTTAGAAAATGTGGCTTTTAATGTTGTAAATAAGGGACACTTTG
ATGGACAACAGGGTGAAGTACCAGTTTCTATCATTAATAACACTGTTTACACAAAAGTTGATGGTGTTGATGTAGAATTGT
TTGAAAATAAAACAACATTACCTGTTAATGTAGCATTTGAGCTTTGGGCTAAGCGCAACATTAAACCAGTACCAGAGGTG
AAAATACTCAATAATTTGGGTGTGGACATTGCTGCTAATACTGTGATCTGGGACTACAAAAGAGATGCTCCAGCACATATA

TCTACTATTGGTGTTTGTTCTATGACTGACATAGCCAAGAAACCAACTGAAACGATTTGTGCACCACTCACTGTCTTTTTTG
ATGGTAGAGTTGATGGTCAAGTAGACTTATTTAGAAATGCCCGTAATGGTGTTCTTATTACAGAAGGTAGTGTCAAGGGC
CTCCAACCGTCGGTAGGCCCCAAGCAAGCTAGGCTGAATGGGGTCACACTGATCGGGGAGGCCGTCAAGACACAGTTCA
ATTATTATAAGAAAGTTGATGGTGTTGTCCAACAATTACCTGAAACTTACTTTACTCAGAGTAGAAATTTACAAGAATTTAA
ACCCAGGAGTCAAATGGAGATAGACTTCCTAGAATTGGCTATGGATGAATTCATCGAGCGGTATAAACTGGAAGGCTATG
CCTTCGAGCATATCGTCTACGGTGACTTCAGCCATAGCCAGCTAGGAGGTCTTCATCTACTGATTGGACTAGCTAAACGTT
TTAAGGAATCACCTTTTGAATTAGAAGATTTTATTCCTATGGACAGTACAGTTAAAAACTATTTCATAACAGATGCGCAAAC
AGGTTCATCTAAGTGTGTATGCTCCGTGATCGATCTGTTACTTGACGACTTTGTTGAGATCATCAAGTCCCAGGACTTATCG
GTGGTtTCCAAAGTCGTCAAAGTAACGATCGACTACACGGAGATCTCATTTATGCTTTGGTGTAAAGATGGCCATGTAGAA
ACATTTTACCCAAAATTACAATCTAGTCAAGCGTGGCAACCGGGTGTTGCTATGCCTAATCTTTACAAAATGCAAAGAATG
CTATTAGAAAAGTGTGACCTTCAAAATTATGGTGATTCGGCGACGCTGCCGAAGGGCATCATGATGAACGTGGCCAAGTA
CACACAACTGTGTCAGTACTTGAACACGCTCACATTGGCTGTGCCCTACAACATGCGAGTCATCCATTTTGGTGCTGGTTCT
GATAAAGGAGTTGCACCAGGTACAGCTGTTTTAAGACAGTGGTTGCCTACGGGTACGCTGCTTGTCGATTCAGATCTGAA
CGACTTTGTCAGCGATGCAGATTCCACCCTGATCGGCGACTGCGCAACTGTGCACACGGCGAACAAGTGGGATTTGATCA
TCTCTGACATGTACGACCCCAAGACTAAAAATGTTACAAAAGAAAATGACTCTAAAGAGGGTTTTTTCACTTACATTTGTG
GGTTTATACAACAAAAGCTAGCTCTTGGAGGTTCCGTGGCTATAAAGATAACAGAACATTCTTGGAATGCTGATCTTTATA
AGCTCATGGGACACTTCGCATGGTGGACAGCCTTTGTTACTAATGTGAATGCGTCATCATCTGAAGCATTTTTAATTGGAT
GTAATTATCTTGGCAAACCACGCGAACAAATAGATGGTTATGTCATGCATGCAAATTACATATTTTGGAGGAATACAAATC
CAATTCAGTTGTCTTCCTATTCTTTATTTGACATGAGTAAATTTCCCCTTAAATTAAGGGGTACTGCTGTTATGTCTTTAAAA
GAAGGTCAAATCAATGATATGATTTTATCTCTTCTTAGTAAAGGTAGACTTATAATTAGAGAAAACAACAGAGTTGTTATT
TCTAGTGATGTTCTTGTTAACAACTAAACGAACAAT

**F7X3X8**
GCTGAAAATGTAACAGGACTCTTTAAAGATTGTAGTAAGGTAATCACTGGGTTACATCCTACACAGGCACCTACACACCTC
AGTGTTGACACTAAATTCAAAACTGAAGGTTTATGTGTTGACATACCTGGCATACCTAAGGACATGACCTATAGAAGACTC
ATCTCTATGATGGGTTTTAAAATGAATTATCAAGTTAATGGTTACCCTAACATGTTTATCACCCGCGAAGAAGCTATAAGA
CATGTACGTGCATGGATTGGCTTCGATGTCGAGGGGTGTCATGCTACTAGAGAAGCTGTTGGTACCAATTTACCTTTACAG
CTAGGTTTTTCTACAGGTGTTAACCTAGTTGCTGTACCTACAGGTTATGTTGATACACCTAATAATACAGATTTTTCCAGAG
TTAGTGCTAAACCACCGCCTGGAGATCAATTTAAACACCTCATACCACTTATGTACAAAGGACTTCCTTGGAATGTAGTGC
GTATAAAGATTGTACAAATGTTAAGTGACACACTTAAAAATCTCTCTGACAGAGTCGTATTTGTCTTATGGGCACATGGCT
TTGAGTTGACATCTATGAAGTATTTTGTGAAAATAGGACCTGAGCGCACCTGTTGTCTATGTGATAGACGTGCCACATGCT
TTTCCACTGCTTCAGACACTTATGCCTGTTGGCATCATTCTATTGGATTTGATTACGTCTATAATCCGTTTATGATTGATGTT
CAACAATGGGGTTTTACAGGTAACCTACAAAGCAACCATGATCTGTATTGTCAAGTCCATGGTAATGCACATGTAGCTAGT
TGTGATGCAATCATGACTAGGTGTCTAGCTGTCCACGAGTGCTTTGTTAAGCGTGTTGACTGGACTATTGAATATCCTATA
ATTGGTGATGAACTGAAGATTAATGCGGCTTGTAGAAAGGTTCAACACATGGTTGTTAAAGCTGCATTATTAGCAGACAA
ATTCCCAGTTCTTCACGACATTGGTAACCCTAAAGCTATTAAGTGTGTACCTCAAGCTGATGTAGAATGGAAGTTCTATGA
TGCACAGCCTTGTAGTGACAAAGCTTATAAAATAGAAGAATTATTCTATTCTTATGCCACACATTCTGACAAATTCACAGAT
GGTGTATGCCTATTTTGGAATTGCAATGTCGATAGATATCCTGCTAATTCCATTGTTTGTAGATTTGACACTAGAGTGCTAT
CTAACCTTAACTTGCCTGGTTGTGATGGTGGCAGTTTGTATGTAAATAAACATGCATTCCACACACCAGCTTTTGATAAAA
GTGCTTTTGTTAATTTAAAACAATTACCATTTTTCTATTACTCTGACAGTCCATGTGAGTCTCATGGAAAACAAGTAGTGTC
AGATATAGATTATGTACCACTAAAGTCTGCTACGTGTATAACACGTTGCAATTTAGGTGGTGCTGTCTGTAGACATCATGC
TAATGAGTACAGATTGTATCTCGATGCTTATAACATGATGATCTCAGCTGGCTTTAGCTTGTGGGTTTACAAACAATTTGAT
ACTTATAACCTCTGGAACACTTTTACAAGACTTCAGAGTTTAGAAAATGTGGCTTTTAATGTTGTAAATAAGGGACACTTTG
ATGGACAACAGGGTGAAGTACCAGTTTCTATCATTAATAACACTGTTTACACAAAAGTTGATGGTGTTGATGTAGAATTGT
TTGAAAATAAAACAACATTACCTGTTAATGTAGCATTTGAGCTTTGGGCTAAGCGCAACATTAAACCAGTACCAGAGGTG
AAAATACTCAATAATTTGGGTGTGGACATTGCTGCTAATACTGTGATCTGGGACTACAAAAGAGATGCTCCAGCACATATA
TCTACTATTGGTGTTTGTTCTATGACTGACATAGCCAAGAAACCAACTGAAACGATTTGTGCACCACTCACTGTCTTTTTTG
ATGGTAGAGTTGATGGTCAAGTAGACTTATTTAGAAATGCCCGTAATGGTGTTCTTATTACAGAAGGTAGTGTCAAGGGC
CTCCAACCGTCGGTAGGCCCCAAGCAAGCTAGGCTGAATGGGGTCACACTGATCGGGGAGGCCGTCAAGACACAGTTCA
ACTACTACAAGAAGGTCGACGGGGTCGTCCAGCAGCTGCCCGAGACtTACTTCACCCAGAGCCGGAACCTGCAGGAGTTC
AAGCCCCGGAGCCAGATGGAGATAGACTTCCTAGAATTGGCTATGGATGAATTCATCGAGCGGTATAAACTGGAAGGCT
ATGCCTTCGAGCATATCGTCTACGGTGACTTCAGCCATAGCCAGCTAGGAGGTCTTCATCTACTGATTGGACTAGCTAAAC
GTTTTAAGGAATCACCTTTTGAATTAGAAGATTTTATTCCTATGGACAGTACAGTTAAAAACTATTTCATAACAGATGCGCA
AACAGGTTCATCTAAGTGTGTATGCTCCGTGATCGATCTGTTACTTGACGACTTTGTTGAGATCATCAAGTCCCAGGACTT
ATCGGTGGTtTCCAAAGTCGTCAAAGTAACGATCGACTACACGGAGATCTCATTTATGCTTTGGTGTAAAGATGGCCATGT
AGAAACATTTTACCCGAAGCTGCAGTCCTCGCAGGCGTGGCAGCCGGGTGTTGCTATGCCGAACCTGTACAACATGCAGC
GGATGCTCCTGGAGAAGTGCGACCTCCAGAACTACGGCGATTCGGCGACGCTGCCGAAGGGCATCATGATGAACGTGGC

CAAGTACACACAACTGTGTCAGTACTTGAACACGCTCACATTGGCTGTGCCCTACAACATGCGAGTCATCCATTTTGGTGC
TGGTTCTGATAAAGGAGTTGCACCAGGTACAGCTGTTTTAAGACAGTGGTTGCCTACGGGTACGCTGCTTGTCGATTCAG
ATCTGAACGACTTTGTCAGCGATGCAGATTCCACCCTGATCGGCGACTGCGCAACTGTGCACACGGCGAACAAGTGGGAT
TTGATCATCTCTGACATGTACGACCCCAAGACTAAAAATGTTACAAAAGAAAATGACTCTAAAGAGGGTTTTTCACTTAC
ATTTGTGGGTTTATACAACAAAAGCTAGCTCTTGGAGGTTCCGTGGCTATAAAGATAACAGAACATTCTTGGAATGCTGAT
CTTTATAAGCTCATGGGACACTTCGCATGGTGGACAGCCTTTGTTACTAATGTGAATGCGTCATCATCTGAAGCATTTTTAA
TTGGATGTAATTATCTTGGCAAACCACGCGAACAAATAGATGGTTATGTCATGCATGCAAATTACATATTTTGGAGGAATA
CAAATCCAATTCAGTTGTCTTCCTATTCTTTATTTGACATGAGTAAATTTCCCCTTAAATTAAGGGGTACTGCTGTTATGTCT
TTAAAAGAAGGTCAAATCAATGATATGATTTTATCTCTTCTTAGTAAAGGTAGACTTATAATTAGAGAAAACAACAGAGTT
GTTATTTCTAGTGATGTTCTTGTTAACAACTAAACGAACAAT

**F7X7**
GCTGAAAATGTAACAGGACTCTTTAAAGATTGTAGTAAGGTAATCACTGGGTTACATCCTACACAGGCACCTACACACCTC
AGTGTTGACACTAAATTCAAAACTGAAGGTTTATGTGTTGACATACCTGGCATACCTAAGGACATGACCTATAGAAGACTC
ATCTCTATGATGGGTTTTAAAATGAATTATCAAGTTAATGGTTACCCTAACATGTTTATCACCCGCGAAGAAGCTATAAGA
CATGTACGTGCATGGATTGGCTTCGATGTCGAGGGGTGTCATGCTACTAGAGAAGCTGTTGGTACCAATTTACCTTTACAG
CTAGGTTTTTCTACAGGTGTTAACCTAGTTGCTGTACCTACAGGTTATGTTGATACACCTAATAATACAGATTTTTCCAGAG
TTAGTGCTAAACCACCGCCTGGAGATCAATTTAAACACCTCATACCACTTATGTACAAAGGACTTCCTTGGAATGTAGTGC
GTATAAAGATTGTACAAATGTTAAGTGACACACTTAAAAATCTCTCTGACAGAGTCGTATTTGTCTTATGGGCACATGGCT
TTGAGTTGACATCTATGAAGTATTTTGTGAAAATAGGACCTGAGCGCACCTGTTGTCTATGTGATAGACGTGCCACATGCT
TTTCCACTGCTTCAGACACTTATGCCTGTTGGCATCATTCTATTGGATTTGATTACGTCTATAATCCGTTTATGATTGATGTT
CAACAATGGGGTTTTACAGGTAACCTACAAAGCAACCATGATCTGTATTGTCAAGTCCATGGTAATGCACATGTAGCTAGT
TGTGATGCAATCATGACTAGGTGTCTAGCTGTCCACGAGTGCTTTGTTAAGCGTGTTGACTGGACTATTGAATATCCTATA
ATTGGTGATGAACTGAAGATTAATGCGGCTTGTAGAAAGGTTCAACACATGGTTGTTAAAGCTGCATTATTAGCAGACAA
ATTCCCAGTTCTTCACGACATTGGTAACCCTAAAGCTATTAAGTGTGTACCTCAAGCTGATGTAGAATGGAAGTTCTATGA
TGCACAGCCTTGTAGTGACAAAGCTTATAAAATAGAAGAATTATTCTATTCTTATGCCACACATTCTGACAAATTCACAGAT
GGTGTATGCCTATTTTGGAATTGCAATGTCGATAGATATCCTGCTAATTCCATTGTTTGTAGATTTGACACTAGAGTGCTAT
CTAACCTTAACTTGCCTGGTTGTGATGGTGGCAGTTTGTATGTAAATAAACATGCATTCCACACACCAGCTTTTGATAAAA
GTGCTTTTGTTAATTTAAAACAATTACCATTTTTCTATTACTCTGACAGTCCATGTGAGTCTCATGGAAAACAAGTAGTGTC
AGATATAGATTATGTACCACTAAAGTCTGCTACGTGTATAACACGTTGCAATTTAGGTGGTGCTGTCTGTAGACATCATGC
TAATGAGTACAGATTGTATCTCGATGCTTATAACATGATGATCTCAGCTGGCTTTAGCTTGTGGGTTTACAAACAATTTGAT
ACTTATAACCTCTGGAACACTTTTACAAGACTTCAGAGTTTAGAAAATGTGGCTTTTAATGTTGTAAATAAGGGACACTTTG
ATGGACAACAGGGTGAAGTACCAGTTTCTATCATTAATAACACTGTTTACACAAAAGTTGATGGTGTTGATGTAGAATTGT
TTGAAAATAAAACAACATTACCTGTTAATGTAGCATTTGAGCTTTGGGCTAAGCGCAACATTAAACCAGTACCAGAGGTG
AAAATACTCAATAATTTGGGTGTGGACATTGCTGCTAATACTGTGATCTGGGACTACAAAAGAGATGCTCCAGCACATATA
TCTACTATTGGTGTTTGTTCTATGACTGACATAGCCAAGAAACCAACTGAAACGATTTGTGCACCACTCACTGTCTTTTTTG
ATGGTAGAGTTGATGGTCAAGTAGACTTATTTAGAAATGCCCGTAATGGTGTTCTTATTACAGAAGGTAGTGTTAAAGGT
TTACAACCATCTGTAGGTCCCAAACAAGCTAGTCTTAATGGAGTCACATTAATTGGAGAAGCCGTAAAAACACAGTTCAAT
TATTATAAGAAAGTTGATGGTGTTGTCCAACAATTACCTGAAACTTACTTTACTCAGAGTAGAAATTTACAAGAATTTAAAC
CCAGGAGTCAAATGGAAATTGATTTCTTAGAATTAGCTATGGATGAATTCATTGAACGGTATAAATTAGAAGGCTATGCCT
TCGAACATATCGTTTATGGAGATTTTAGTCATAGTCAGTTAGGTGGTTTACATCTACTGATTGGACTAGCTAAACGTTTTAA
GGAATCACCTTTTGAATTAGAAGATTTTATTCCTATGGACAGTACAGTTAAAAACTATTTCATAACAGATGCGCAAACAGG
TTCATCTAAGTGTGTGTGCAGCGTTATAGACTTACTGTTAGATGATTTTGTGGAAATCATCAAGTCTCAGGATCTTTCCGTC
GTCAGTAAGGTTGTTAAGGTGACTATAGACTATACCGAAATTTCATTTATGCTTTGGTGTAAAGATGGCCATGTAGAAACA
TTTTACCCAAAATTACAATCTAGTCAAGCGTGGCAACCGGGTGTTGCTATGCCTAATCTTTACAAAATGCAAAGAATGCTA
TTAGAAAAGTGTGACCTTCAAAATTATGGTGATAGTGCAACATTACCTAAAGGCATAATGATGAATGTCGCAAAATATACT
CAACTGTGTCAATATTTAAACACATTAACATTAGCTGTACCCTATAATATGAGAGTTATACATTTTGGTGCTGGTTCTGATA
AAGGAGTTGCACCAGGTACAGCTGTTTTAAGACAGTGGTTGCCTACGGGTACGCTGCTTGTCGATTCAGATCTTAATGACT
TTGTCTCTGATGCAGATTCAACTTTGATTGGTGATTGTGCAACTGTACATACAGCTAATAAATGGGATCTCATTATTAGTGA
TATGTACGACCCTAAGACTAAAAATGTTACAAAAGAAAATGACTCTAAAGAGGGTTTTTCACTTACATTTGTGGGTTTAT
ACAACAAAAGCTAGCTCTTGGAGGTTCCGTGGCTATAAAGATAACAGAACATTCTTGGAATGCTGATCTTTATAAGCTCAT
GGGACACTTCGCATGGTGGACAGCCTTTGTTACTAATGTGAATGCGTCATCATCTGAAGCATTTTTAATTGGATGTAATTA
TCTTGGCAAACCACGCGAACAAATAGATGGTTATGTCATGCATGCAAATTACATATTTTGGAGGAATACAAATCCAATTCA
GTTGTCTTCCTATTCTTTATTTGACATGAGTAAATTTCCCCTTAAATTAAGGGGTACTGCTGTTATGTCTTTAAAAGAAGGTC
AAATCAATGATATGATTTTATCTCTTCTTAGTAAAGGTAGACTTATAATTAGAGAAAACAACAGAGTTGTTATTTCTAGTGA
TGTTCTTGTTAACAACTAAACGAACAAT

**F1D**

ATTAAAGGTTTATACCTTCCCAGGTAACAAACCAACCAACTTTCGATCTCTTGTAGATCTGTTCTCTAAACGAACTTTAAAA
TCTGTGTGGCTGTCACTCGGCTGCATGCTTAGTGCACTCACGCAGTATAATTAATAACTAATTACTGTCGTTGACAGGACA
CGAGTAACTCGTCTATCTTCTGCAGGCTGCTTACGGTTTCGTCCGTGTTGCAGCCGATCATCAGCACATCTAGGTTTCGTCC
GGGTGTGACCGAAAGGTAAGATGGAGAGCCTTGTCCCTGGTTTCAACGAGAAAACACACGTCCAACTCAGTTTGCCTGTT
TTACAGGTTCGCGACGTGCTCGTACGTGGCTTTGGAGACTCCGTGGAGGAGGTCTTATCAGAGGCACGTCAACATCTTAA
AGATGGCACTTGTGGCTTAGTAGAAGTTGAAAAAGGCGTTTTGCCTCAACTTGAACAGCCCTATGTGTTCATCAAACGTTC
GGATGCTCGAACTGCACCTCATGGTCATGTTATGGTTGAGCTGGTAGCAGAACTCGAAGGCATTCAGTACGGTCGTAGTG
GTGAGACACTTGGTGTCCTTGTCCCTCATGTGGGCGAAATACCAGTGGCTTACCGCAAGGTTCTTCTTCGTAAGAACGGTA
ATAAAGGAGCTGGTGGCCATAGTTACGGCGCCGATCTAAAGTCATTTGACTTAGGCGACGAGCTTGGCACTGATCCTTAT
GAAGATTTTCAAGAAAACTGGAACACTAAACATAGCAGTGGTGTTACCCGTGAACTCATGCGTGAGCTTAACGGAGGGG
CATACACTCGCTATGTCGATAACAACTTCTGTGGCCCGGATGGCTACCCGCTGGAGTGCATTAAAGACCTGCTGGCACGG
GCGGGGAAAGCGAGCTGCACGCTGAGCGAACAACTGGACTTTATTGACACGAAGAGGGGGGTATACTGCTGCCGGGAA
CATGAGCATGAAATTGCGTGGTACACGGAACGGAGCGAAAAGAGCTATGAACTGCAGACACCGTTTGAAATTAAACTGG
CAAAGAAATTTGACACCTTCAATGGGGAATGTCCAAATTTTGTATTTCCCCTGAATAGCATAATCAAGACGATTCAACCAA
GGGTGGAAAAGAAAAAGCTGGATGGCTTTATGGGGAGAATTCGAAGCGTCTATCCAGTGGCGAGCCCAAATGAATGCA
ACCAAATGTGCCTGAGCACGCTGATGAAGTGTGATCATTGTGGGGAAACGAGCTGGCAGACGGGCGATTTTGTGAAAGC
CACGTGCGAATTTTGTGGCACGGAGAATCTGACGAAAGAAGGGGCCACGACGTGTGGGTACCTGCCCCAAAATGCGGTG
GTGAAAATTTATTGTCCAGCATGTCACAATAGCGAAGTAGGACCGGAGCATAGCCTGGCCGAATACCATAATGAAAGCG
GCCTGAAAACCATTCTGCGGAAGGGGGGGCGCACGATTGCCTTTGGAGGCTGTGTGTTCAGCTATGTGGGGTGCCATAA
CAAGTGTGCCTATTGGGTGCCACGGGCGAGCGCGAACATAGGGTGTAACCATACAGGGGTGGTGGGAGAAGGGAGCGA
AGGGCTGAATGACAACCTGCTGGAAATACTGCAAAAAGAGAAAGTCAACATCAATATTGTGGGGGACTTTAAACTGAAT
GAAGAGATCGCCATTATTCTGGCAAGCTTTAGCGCGAGCACAAGCGCGTTTGTGGAAACGGTGAAAGGGCTGGATTATA
AAGCATTCAAACAAATTGTGGAAAGCTGTGGGAATTTTAAAGTGACAAAAGGAAAAGCGAAAAAAGGGGCCTGGAATAT
TGGGGAACAGAAAAGCATACTGAGCCCGCTGTATGCATTTGCAAGCGAGGCGGCGCGGGTGGTACGAAGCATTTTCAGC
CGCACGCTGGAAACGGCGCAAAATAGCGTGCGGGTGCTGCAGAAGGCCGCGATAACAATACTGGATGGAATTAGCCAG
TATAGCCTGAGACTGATTGATGCGATGATGTTCACAAGCGATCTGGCGACGAACAATCTGGTGGTAATGGCCTACATTAC
AGGGGGGGTGGTGCAGCTGACGAGCCAGTGGCTGACGAACATCTTTGGCACGGTGTATGAAAAACTGAAACCCGTCCTG
GATTGGCTGGAAGAGAAGTTTAAGGAAGGGGTAGAGTTTCTGAGAGACGGGTGGGAAATTGTGAAATTTATCAGCACCT
GTGCGTGTGAAATTGTCGGGGGACAAATTGTCACCTGTGCAAAGGAAATTAAGGAGAGCGTGCAGACATTCTTTAAGCT
GGTAAATAAATTTCTGGCGCTGTGTGCGGACAGCATCATTATTGGGGGAGCGAAACTGAAAGCCCTGAATCTGGGGGAA
ACATTTGTCACGCACAGCAAGGGACTGTACAGAAAGTGTGTGAAAAGCAGAGAAGAAACGGGCCTGCTGATGCCGCTGA
AAGCCCCAAAAGAAATTATCTTCCTGGAGGGAGAAACACTGCCCACAGAAGTGCTGACAGAGGAAGTGGTCCTGAAAAC
GGGGGATCTGCAACCACTGGAACAACCGACGAGCGAAGCGGTGGAAGCGCCACTGGTGGGGACACCAGTGTGTATTAA
CGGGCTGATGCTGCTGGAAATCAAAGACACAGAAAAGTACTGTGCCCTGGCACCGAATATGATGGTAACAAACAATACCT
TCACACTGAAAGGCGGTGC

**F1E**

ATTAAAGGTTTATACCTTCCCAGGTAACAAACCAACCAACTTTCGATCTCTTGTAGATCTGTTCTCTAAACGAACTTTAAAA
TCTGTGTGGCTGTCACTCGGCTGCATGCTTAGTGCACTCACGCAGTATAATTAATAACTAATTACTGTCGTTGACAGGACA
CGAGTAACTCGTCTATCTTCTGCAGGCTGCTTACGGTTTCGTCCGTGTTGCAGCCGATCATCAGCACATCTAGGTTTCGTCC
GGGTGTGACCGAAAGGTAAGATGGAGAGCCTTGTCCCTGGTTTCAACGAGAAAACACACGTCCAACTCAGTTTGCCTGTT
TTACAGGTTCGCGACGTGCTCGTACGTGGCTTTGGAGACTCCGTGGAGGAGGTCTTATCAGAGGCACGTCAACATCTTAA
AGATGGCACTTGTGGCTTAGTAGAAGTTGAAAAAGGCGTTTTGCCTCAACTTGAACAGCCCTATGTGTTCATCAAACGTTC
GGATGCTCGAACTGCACCTCATGGTCATGTTATGGTTGAGCTGGTAGCAGAACTCGAAGGCATTCAGTACGGTCGTAGTG
GTGAGACACTTGGTGTCCTTGTCCCTCATGTGGGCGAAATACCAGTGGCTTACCGCAAGGTTCTTCTTCGTAAGAACGGTA
ATAAAGGAGCTGGTGGCCATAGTTACGGCGCCGATCTAAAGTCATTTGACTTAGGCGACGAGCTTGGCACTGATCCTTAT
GAAGATTTTCAAGAAAACTGGAACACTAAACATAGCAGTGGTGTTACCCGTGAACTCATGCGTGAGCTTAACGGAGGGG
CATACACTCGTTATGTTGATAATAATTTTTGTGGTCCTGATGGTTATCCTTTAGAGTGTATTAAAGATTTATTAGCTCGTGCT
GGTAAAGCTAGTTGTACTTTAAGTGAACAATTAGATTTTATTGATACTAAGCGTGGTGTTTATTGTTGTCGTGAACATGAG
CATGAAATTGCTTGGTATACTGAACGTAGTGAAAAGAGTTATGAATTACAGACTCCTTTTGAAATTAAATTAGCTAAGAAA
TTTGATACTTTTAATGGTGAATGTCCTAATTTTGTTTTTCCTTTAAATAGTATTATTAAGACTATTCAACCTCGTGTTGAAAA
GAAAAAGTTAGATGGTTTTATGGGTCGTATTCGTAGTGTTTATCCTGTTGCTAGTCCTAATGAATGTAATCAAATGTGTTTA
AGTACTTTAATGAAGTGTGATCATTGTGGTGAAACTAGTTGGCAGACTGGTGATTTTGTTAAAGCTACTTGTGAATTTTGT
GGTACTGAGAATTTAACTAAAGAAGGTGCTACTACTTGTGGTTATTTACCTCAAAATGCTGTTGTTAAAATTTATTGTCCTG
CTTGTCATAATAGTGAAGTTGGTCCTGAGCATAGTTTAGCTGAATATCATAATGAAAGTGGTTTAAAAACTATTTTACGTA
AGGGTGGTCGTACTATTGCTTTTGGTGGTTGTGTTTTTAGTTATGTTGGTTGTCATAATAAGTGTGCTTATTGGGTTCCTCG

TGCTAGTGCTAATATTGGTTGTAATCATACTGGTGTTGTTGGTGAAGGTAGTGAAGGTTTAAATGATAATTTATTAGAAAT
TTTACAAAAAGAGAAAGTTAATATTAATATTGTTGGTGATTTTAAATTAAATGAAGAGATTGCTATTATTTTAGCTAGTTTT
AGTGCTAGTACTAGTGCTTTTGTTGAAACTGTTAAAGGTTTAGATTATAAAGCTTTTAAACAAATTGTTGAAAGTTGTGGT
AATTTTAAAGTTACTAAAGGTAAAGCTAAAAAAGGTGCTTGGAATATTGGTGAACAGAAAAGTATTTTAAGTCCTTTATAT
GCTTTTGCTAGTGAGGCTGCTCGTGTTGTTCGTAGTATTTTTAGTCGTACTTTAGAAACTGCTCAAAATAGTGTTCGTGTTT
TACAGAAGGCTGCTATTACTATTTTAGATGGTATTAGTCAGTATAGTTTACGTTTAATTGATGCTATGATGTTTACTAGTGA
TTTAGCTACTAATAATTTAGTTGTTATGGCTTATATTACTGGTGGTGTTGTTCAGTTAACTAGTCAGTGGTTAACTAATATTT
TTGGTACTGTTTATGAAAAATTAAAACCTGTTTTAGATTGGTTAGAAGAGAAGTTTAAGGAAGGTGTTGAGTTTTTACGTG
ATGGTTGGGAAATTGTTAAATTTATTAGTACTTGTGCTTGTGAAATTGTTGGTGGTCAAATTGTTACTTGTGCTAAGGAAA
TTAAGGAGAGTGTTCAGACTTTTTTTAAGTTAGTTAATAAATTTTTAGCTTTATGTGCTGATAGTATTATTATTGGTGGTGC
TAAATTAAAAGCTTTAAATTTAGGTGAAACTTTTGTTACTCATAGTAAGGGTTTATATCGTAAGTGTGTTAAAAGTCGTGA
AGAAACTGGTTTATTAATGCCTTTAAAAGCTCCTAAAGAAATTATTTTTTTAGAGGGTGAAACTTTACCTACTGAAGTTTTA
ACTGAGGAAGTTGTTTTAAAAACTGGTGATTTACAACCTTTAGAACAACCTACTAGTGAAGCTGTTGAAGCTCCTTTAGTT
GGTACTCCTGTTTGTATTAATGGTTTAATGTTATTAGAAATTAAAGATACTGAAAAGTATTGTGCTTTAGCTCCTAATATGA
TGGTTACTAATAATACTTTTACTTTAAAAGGTGGTGC

**F2D**

GTGCACCAACAAAGGTGACGTTTGGGGATGACACGGTGATAGAAGTGCAAGGGTACAAGAGCGTGAATATCACGTTTGA
ACTGGATGAAAGGATTGATAAAGTACTGAATGAGAAGTGCAGCGCCTATACAGTGGAACTGGGGACAGAAGTAAATGA
GTTCGCCTGTGTGGTGGCAGATGCGGTCATAAAAACGCTGCAACCAGTAAGCGAACTGCTGACACCACTGGGCATTGATC
TGGATGAGTGGAGCATGGCGACATACTACCTGTTTGATGAGAGCGGGGAGTTTAAACTGGCGAGCCATATGTATTGTAG
CTTCTACCCGCCAGATGAGGATGAAGAAGAAGGGGATTGTGAAGAAGAAGAGTTTGAGCCAAGCACGCAATATGAGTAT
GGGACGGAAGATGATTACCAAGGGAAACCGCTGGAATTTGGGGCCACGAGCGCGGCGCTGCAACCGGAAGAAGAGCA
AGAAGAAGATTGGCTGGATGATGATAGCCAACAAACGGTGGGGCAACAAGACGGCAGCGAGGACAATCAGACAACGAC
GATTCAAACAATTGTGGAGGTGCAACCGCAACTGGAGATGGAACTGACACCAGTGGTGCAGACGATTGAAGTGAATAGC
TTTAGCGGGTATCTGAAACTGACGGACAATGTATACATTAAAAATGCAGACATTGTGGAAGAAGCGAAAAAGGTAAAAC
CAACAGTGGTGGTGAATGCAGCCAATGTGTACCTGAAACATGGAGGAGGGGTGGCAGGAGCCCTGAATAAGGCGACGA
ACAATGCCATGCAAGTGGAAAGCGATGATTACATAGCGACGAATGGACCACTGAAAGTGGGGGGGAGCTGTGTGCTGA
GCGGACACAATCTGGCGAAACACTGTCTGCATGTGGTCGGCCCAAATGTGAACAAAGGGGAAGACATTCAACTGCTGAA
GAGCGCGTATGAAAATTTTAATCAGCACGAAGTGCTGCTGGCACCACTGCTGAGCGCGGGGATTTTTGGGGCGGACCCG
ATACATAGCCTGAGAGTGTGTGTAGATACGGTGCGCACAAATGTCTACCTGGCGGTCTTTGATAAAAATCTGTATGACAA
ACTGGTGAGCAGCTTTCTGGAAATGAAGAGCGAAAAGCAAGTGGAACAAAAGATCGCGGAGATTCCGAAAGAGGAAGT
GAAGCCATTTATAACGGAAAGCAAACCGAGCGTGGAACAGAGAAAACAAGATGATAAGAAAATCAAAGCGTGTGTGGA
AGAAGTGACAACAACGCTGGAAGAAACGAAGTTCCTGACAGAAAACCTGCTGCTGTATATTGACATTAATGGCAATCTGC
ATCCAGATAGCGCCACGCTGGTGAGCGACATTGACATCACGTTCCTGAAGAAAGATGCGCCATATATAGTGGGGGATGT
GGTGCAAGAGGGGGTGCTGACGGCGGTGGTGATACCGACGAAAAAGGCGGGGGGCACGACGGAAATGCTGGCGAAAG
CGCTGAGAAAAGTGCCAACAGACAATTATATAACCACGTACCCGGGGCAGGGGCTGAATGGGTACACGGTAGAGGAGG
CAAAGACAGTGCTGAAAAAGTGTAAAAGCGCCTTTTACATTCTGCCAAGCATTATCAGCAATGAGAAGCAAGAAATTCTG
GGAACGGTGAGCTGGAATCTGCGAGAAATGCTGGCACATGCAGAAGAAACACGCAAACTGATGCCGGTCTGTGTGGAA
ACGAAAGCCATAGTGAGCACGATACAGCGGAAATATAAGGGGATTAAAATACAAGAGGGGGTGGTGGATTATGGGGCG
AGATTTTACTTTTACACCAGCAAAACAACGGTAGCGAGCCTGATCAACACACTGAACGATCTGAATGAAACGCTGGTGAC
AATGCCACTGGGCTATGTAACACATGGCCTGAATCTGGAAGAAGCGGCGCGGTATATGAGAAGCCTGAAAGTGCCAGCG
ACAGTGAGCGTGAGCAGCCCGGATGCGGTGACAGCGTATAATGGGTATCTGACGAGCAGCAGCAAAACACCGGAAGAA
CATTTTATTGAAACCATCAGCCTGGCGGGGAGCTATAAAGATTGGAGCTATAGCGGACAAAGCACACAACTGGGGATAG
AATTTCTTAAGAGAGGTGATAAAAGTGTATATTACACTAGTAATCCTACCACATTCCACCTAGATGGTGAAGTTATCACCTT
TGACAATCTTAAGACACTTCTTTCTTTGAGAGAAGTGAGGACTATTAAGGTGTTTACAACAGTAGACAACATTAACCTCCA
CACGCAAGTTGTGGACATGTCAATGACATATGGACAACAGTTTGGTCCAACTTATTTGGATGGAGCTGATGTTACTAAAAT
AAAACCTCATAATTCACATGAAGGTAAAACATTTTATGTTTTACCTAATGATGACACTCTACGTGTTGAGGCTTTTGAGTAC
TACCACACAACTGATCCTAGTTTTCTGGGTAGGTACATGTCAGCATTAAATCACACTAAAAAGTGGAAATACCCACAAGTT
AATGGTTTAACTTCTATTAAATGGGCAGATAACAACTGTTATCTTGCCACTGCATTGTTAACACTCCAACAAATAGAGTTGA
AGTTTAATCCACCTGCTCTACAAGATGCTTATTACAGAGCAAGGGCTGGTGAAGCTGCTAACTTTTGTGCACTTATCTTAG
CCTACTGTAATAAGACAGTAGGTGAGTTAGGTGATGTTAGAGA

**F2E**

GTGCTCCTACTAAGGTTACTTTTGGTGATGATACTGTTATTGAAGTTCAAGGTTATAAGAGTGTTAATATTACTTTTGAATT
AGATGAACGTATTGATAAAGTTTTAAATGAGAAGTGTAGTGCTTATACTGTTGAATTAGGTACTGAAGTTAATGAGTTTGC
TTGTGTTGTTGCTGATGCTGTTATTAAAACTTTACAACCTGTTAGTGAATTATTAACTCCTTTAGGTATTGATTTAGATGAGT

GGAGTATGGCTACTTATTATTTATTTGATGAGAGTGGTGAGTTTAAATTAGCTAGTCATATGTATTGTAGTTTTTATCCTCC
TGATGAGGATGAAGAAGAAGGTGATTGTGAAGAAGAAGAGTTTGAGCCTAGTACTCAATATGAGTATGGTACTGAAGAT
GATTATCAAGGTAAACCTTTAGAATTTGGTGCTACTAGTGCTGCTTTACAACCTGAAGAAGAGCAAGAAGAAGATTGGTT
AGATGATGATAGTCAACAAACTGTTGGTCAACAAGATGGTAGTGAGGATAATCAGACTACTACTATTCAAACTATTGTTG
AGGTTCAACCTCAATTAGAGATGGAATTAACTCCTGTTGTTCAGACTATTGAAGTTAATAGTTTTAGTGGTTATTTAAAATT
AACTGATAATGTTTATATTAAAAATGCTGATATTGTTGAAGAAGCTAAAAAGGTTAAACCTACTGTTGTTGTTAATGCTGCT
AATGTTTATTTAAAACATGGTGGTGGTGTTGCTGGTGCTTTAAATAAGGCTACTAATAATGCTATGCAAGTTGAAAGTGAT
GATTATATTGCTACTAATGGTCCTTTAAAAGTTGGTGGTAGTTGTGTTTTAAGTGGTCATAATTTAGCTAAACATTGTTTAC
ATGTTGTTGGTCCTAATGTTAATAAAGGTGAAGATATTCAATTATTAAAGAGTGCTTATGAAAATTTTAATCAGCATGAAG
TTTTATTAGCTCCTTTATTAAGTGCTGGTATTTTTGGTGCTGATCCTATTCATAGTTTACGTGTTTGTGTTGATACTGTTCGT
ACTAATGTTTATTTAGCTGTTTTTGATAAAAATTTATATGATAAATTAGTTAGTAGTTTTTTAGAAATGAAGAGTGAAAAGC
AAGTTGAACAAAAGATTGCTGAGATTCCTAAAGAGGAAGTTAAGCCTTTTATTACTGAAAGTAAACCTAGTGTTGAACAG
CGTAAACAAGATGATAAGAAAATTAAAGCTTGTGTTGAAGAAGTTACTACTACTTTAGAAGAAACTAAGTTTTTAACTGAA
AATTTATTATTATATATTGATATTAATGGTAATTTACATCCTGATAGTGCTACTTTAGTTAGTGATATTGATATTACTTTTTTA
AAGAAAGATGCTCCTTATATTGTTGGTGATGTTGTTCAAGAGGGTGTTTTAACTGCTGTTGTTATTCCTACTAAAAAGGCT
GGTGGTACTACTGAAATGTTAGCTAAAGCTTTACGTAAAGTTCCTACTGATAATTATATTACTACTTATCCTGGTCAGGGTT
TAAATGGTTATACTGTTGAGGAGGCTAAGACTGTTTTAAAAAAGTGTAAAAGTGCTTTTTATATTTTACCTAGTATTATTAG
TAATGAGAAGCAAGAAATTTTAGGTACTGTTAGTTGGAATTTACGTGAAATGTTAGCTCATGCTGAAGAAACTCGTAAATT
AATGCCTGTTTGTGTTGAAACTAAAGCTATTGTTAGTACTATTCAGCGTAAATATAAGGGTATTAAAATTCAAGAGGGTGT
TGTTGATTATGGTGCTCGTTTTTATTTTTATACTAGTAAAACTACTGTTGCTAGTTTAATTAATACTTTAAATGATTTAAATG
AAACTTTAGTTACTATGCCTTTAGGTTATGTTACTCATGGTTTAAATTTAGAAGAAGCTGCTCGTTATATGCGTAGTTTAAA
AGTTCCTGCTACTGTTAGTGTTAGTAGTCCTGATGCTGTTACTGCTTATAATGGTTATTTAACTAGTAGTAGTAAAACTCCT
GAAGAACATTTTATTGAAACTATTAGTTTAGCTGGTAGTTATAAAGATTGGAGTTATAGTGGTCAAAGTACTCAATTAGGT
ATTGAATTTCTTAAGAGAGGTGATAAAAGTGTATATTACACTAGTAATCCTACCACATTCCACCTAGATGGTGAAGTTATC
ACCTTTGACAATCTTAAGACACTTCTTTCTTTGAGAGAAGTGAGGACTATTAAGGTGTTTACAACAGTAGACAACATTAAC
CTCCACACGCAAGTTGTGGACATGTCAATGACATATGGACAACAGTTTGGTCCAACTTATTTGGATGGAGCTGATGTTACT
AAAATAAAACCTCATAATTCACATGAAGGTAAAACATTTTATGTTTTACCTAATGATGACACTCTACGTGTTGAGGCTTTTG
AGTACTACCACACAACTGATCCTAGTTTTCTGGGTAGGTACATGTCAGCATTAAATCACACTAAAAAGTGGAAATACCCAC
AAGTTAATGGTTTAACTTCTATTAAATGGGCAGATAACAACTGTTATCTTGCCACTGCATTGTTAACACTCCAACAAATAGA
GTTGAAGTTTAATCCACCTGCTCTACAAGATGCTTATTACAGAGCAAGGGCTGGTGAAGCTGCTAACTTTTGTGCACTTAT
CTTAGCCTACTGTAATAAGACAGTAGGTGAGTTAGGTGATGTTAGAGA
